# LDO proteins and Vac8 form a vacuole-lipid droplet contact site required for lipophagy in response to starvation

**DOI:** 10.1101/2023.04.21.537797

**Authors:** Irene Álvarez-Guerra, Emma Block, Filomena Broeskamp, Sonja Gabrijelčič, Ana de Ory, Lukas Habernig, Claes Andréasson, Tim P. Levine, Johanna L. Höög, Sabrina Büttner

## Abstract

Lipid droplets (LDs) are fat storage organelles critical for energy and lipid metabolism. Upon nutrient exhaustion, cells consume LDs via gradual lipolysis or via lipophagy, the *en bloc* uptake of LDs into the vacuole. Here, we show that LDs dock to the vacuolar membrane via a contact site that is required for lipophagy in yeast. The LD-localized LDO proteins carry an intrinsically disordered region that associates with vacuolar Vac8 to form vCLIP, the vacuolar-LD contact site. Nutrient limitation drives vCLIP formation, and its inactivation blocks lipophagy. Vac8 is sufficient to recruit LDs to cellular membranes. We establish a functional link between lipophagy and microautophagy of the nucleus, both requiring Vac8 to form respective contact sites upon metabolic stress. In sum, we unravel the molecular architecture of vCLIP, a contact site required for lipophagy, and find that Vac8 provides a platform for multiple and competing contact sites associated with autophagy.

## Introduction

Cellular adaptation to changing metabolic demands requires efficient communication between organelles and remodeling of subcellular structures and processes. Direct physical contact between organelles at dedicated membrane contact sites provides ancient communication routes and hubs for metabolic exchange present in organisms ranging from yeast to humans (Bohnert, 2020; Eisenberg-Bord et al., 2016; Scorrano et al., 2019). Membrane contact sites are established by an array of tethering proteins that bridge virtually all pairs of organelles, facilitating the bidirectional exchange of biochemical information in form of metabolites, ions and lipids (Eisenberg-Bord et al., 2016; Phillips and Voeltz, 2016; Prinz et al., 2020; Scorrano et al., 2019). These organelle contacts are key to intracellular signaling and are emerging as important sites for metabolic adaptation (Bohnert, 2020; Henne, 2016; Kohler and Büttner, 2021).

In yeast, a contact site that dynamically changes size and molecular architecture in response to nutrient availability is the nucleus-vacuole junction (NVJ). NVJ connects the main anabolic and catabolic cellular compartments by establishing proximity between the nuclear endoplasmic reticulum (nER) and the vacuole, the yeast counterpart of mammalian lysosomes (Hariri et al., 2018; Kvam and Goldfarb, 2007; Pan et al., 2000; Rogers et al., 2021; Tosal-Castano et al., 2021). When nutrients are depleted, NVJs expand and serve as a platform for the coordination of distinct aspects of lipid metabolism, including the subcellular organization of lipid droplets (LDs) (Barbosa and Siniossoglou, 2016; Hariri et al., 2018; Rogers et al., 2021). LDs are dynamic fat storage organelles with critical roles in lipid and energy metabolism that allow cells to adapt to changing nutritional cues (Olzmann and Carvalho, 2019; Walther and Farese, 2012). They consist of a core of neutral lipids, particularly triacylglycerides and sterol esters, delimited by a phospholipid monolayer that originates from the outer leaflet of the ER membrane and contains a number of integral and peripheral proteins (Olzmann and Carvalho, 2019). Due to the absence of a bilayer, LDs cannot engage in vesicle trafficking but rely on communication via membrane contact sites, established with several other organelles (Herker et al., 2021; Liao et al., 2022; Schuldiner and Bohnert, 2017; Shai et al., 2018; Thiam and Dugail, 2019; Wang et al., 2021).

Depending on the metabolic status, cells alternate between LD biogenesis at the ER and LD consumption via gradual enzymatic lipolysis or via lipophagy (Grabner et al., 2021; Graef, 2018; Shin, 2020; Singh et al., 2009). Lipophagy is based on *en bloc* import into the lysosome/vacuole and plays a key role in efficient LD consumption. In mammalian cells, uptake into the lysosome is mediated by macroautophagy, in which autophagosomes sequester LDs for subsequent delivery to the lysosome (Garcia et al., 2018; Zhang et al., 2018). Notably, an alternative lipophagic route for fatty acid mobilization from LDs has recently been described in hepatocytes, which are particularly rich in LDs (Schulze et al., 2020). Here, autophagosome intermediates are absent and LDs directly dock to lysosomes to be engulfed in a process that resembles microlipophagy, which is the main route of lipophagic LD consumption in yeast cells (Schott et al., 2022; Seo et al., 2017; van Zutphen et al., 2014; Vevea et al., 2015). During microlipophagy, LDs redistribute around the yeast vacuole and associate with the liquid-ordered (Lo) microdomains on the vacuolar membrane before being engulfed by the vacuole (Liao et al., 2021; Wang et al., 2014). Though contact formation via molecules that bridge the LDs and the vacuole/lysosome is anticipated to support the subsequent lipophagic LD engulfment, no such molecules have been identified. Still, proximity-based approaches employing LD-localized and vacuolar resident proteins suggest the existence of such a <u>v</u>a<u>c</u>uolar-<u>lip</u>id droplet contact site (vCLIP) (Shai et al., 2018).

Here, we uncover the molecular architecture of the contact site that anchors LDs to the vacuole to facilitate lipophagy in response to nutrient exhaustion (also see Diep *et al*., submitted in parallel to this study). We demonstrate that the <u>LD</u> <u>o</u>rganization proteins Ldo16 and Ldo45, encoded by overlapping genes and products of an alternative splicing event, accumulate at distinct foci on LDs to attach them to the vacuolar membrane via Vac8, forming the <u>v</u>a<u>c</u>uole-<u>lip</u>id droplet contact site (vCLIP). Genetic ablation of the LDO proteins or of Vac8 disrupted vCLIP formation and lipophagy. Ldo16 was found to be transcriptionally upregulated specifically upon nutrient depletion and to anchor LDs to Vac8 via its C-terminal intrinsically disordered region. Experimentally redirecting Vac8 to the nuclear membrane was sufficient to re-route LDs to juxtanuclear locations. In sum, we identify the molecular bridges that form vCLIP and show that this contact is critical for lipophagic LD consumption when nutrients become limiting.

## Results

### LDO proteins accumulate at the vacuole-LD interface in response to nutrient exhaustion

The shift from rapid proliferation in nutrient-rich conditions to stationary phase upon nutrient exhaustion is associated with a metabolic switch that involves the transition from LD biogenesis to storage and gradual LD consumption. To assess how this transition affects LD-localized proteins, we followed the subcellular distribution of endogenous GFP fusions of the fatty acyl-CoA synthetase Faa4, the ergosterol biosynthesis enzyme Erg1, the phosphatidylinositol transfer protein Pdr16 and the LD- organization proteins Ldo16/45. Ldo16/45 have been reported to associate with a specific LD subpopulation at the NVJs (Eisenberg-Bord et al., 2018; Teixeira et al., 2018) and are putative homologs of the mammalian Lipid droplet assembly factor 1 LDAF1/Promethin (Castro et al., 2019). We analyzed cells during exponential growth in glucose-rich conditions (8 h) as well as after prolonged incubation (48 h), which results in gradual exhaustion of glucose, followed by a switch to respiratory metabolism and entry into stationary phase (Tosal-Castano et al., 2021). In cells growing in glucose-rich conditions, all GFP fusion proteins decorated the LDs, visualized using the neutral lipid stain monodansylpentane (MDH) (Fig. 1A). As reported previously (Müllner et al., 2004), a portion of Erg1 was retained in the ER (Fig. 1A). In glucose-exhausted cells, Faa4 and Erg1 remained evenly distributed on the surface of enlarged LDs, while Pdr16 and in particular the LDO proteins formed distinct foci on LDs, mostly limited to one structure per LD (Fig. 1A). Simultaneous visualization of the vacuole using Vph1^mCherry^ revealed that these foci mark the interface between the vacuole and LDs, suggesting that Pdr16 and the LDO proteins are redirected to membrane contact sites between these two organelles when glucose is exhausted (Fig. 1B). The targeting of the LDO proteins to the vacuolar-LD interface was not affected by genetic ablation of Pdr16 (Fig. 1C). In contrast, the lack of both LDO proteins (ΔΔ*ldo*) completely prevented the accumulation of Pdr16 at these sites and resulted in its cytosolic distribution (Fig. 1C), suggesting that the LDO proteins are required to recruit Pdr16 to these contact sites. The two LDO proteins are encoded by overlapping open reading frames (Fig. 1D), and C-terminal tagging with GFP (referred to as LDO^GFP^) thus results in the simultaneous microscopic visualization of both proteins (Fig. 1A-C). Immunoblotting demonstrated that the expression of Ldo45 and Ldo16 was differentially affected by nutrient depletion (Fig. 1E, F). Ldo16 protein levels progressively increased with time spent in stationary phase, while the levels of Ldo45 slightly decreased (Fig. 1E, F). Likewise, *LDO16* mRNA levels (but not *LDO45* levels) increased over time, indicating transcriptional upregulation of specifically Ldo16 (Supplemental Figure S1A). Simultaneous visualisation of Faa4^mCherry^ and LDO^GFP^ in glucose-exhausted cells demonstrated that the LDO proteins were targeted selectively to the vacuole-LD interface, while Faa4 remained evenly distributed at the LD surface, independently of which fluorescent tags were used (Fig. 1G). Next, we ruled out that the starvation-induced targeting of LDO^GFP^ to the vacuole-LD interface was a simple consequence of increased LD size by expanding the LDs in exponentially growing cells on glucose-rich media using oleate supplementation (Exner et al., 2019). Although oleate prominently enlarged the LDs, the LDO proteins mainly remained distributed over the LD surface and only accumulated at the vacuole-LD interface in few, small foci when glucose was still available (Fig. 1H). Again, also under oleate supplemented conditions, entry into stationary phase resulted in the targeting of the LDO proteins to the prominently expanded contact sites between the enlarged LDs and the vacuole (Fig. 1H). Furthermore, subjecting cells to phosphate restriction or acute nitrogen depletion resulted in the partitioning of the LDO proteins to the vacuole-LD interface (Fig. 1I) and a specific increase of Ldo16 (Supplemental Fig. S1B-D). Collectively, this shows that limitation of the macronutrients glucose, nitrogen or phosphate induces Ldo16 expression and results in the specific targeting of the LDO proteins to the vacuole-LD interface, likely reflecting vCLIP that emerge in response to starvation.

**Figure 1:**
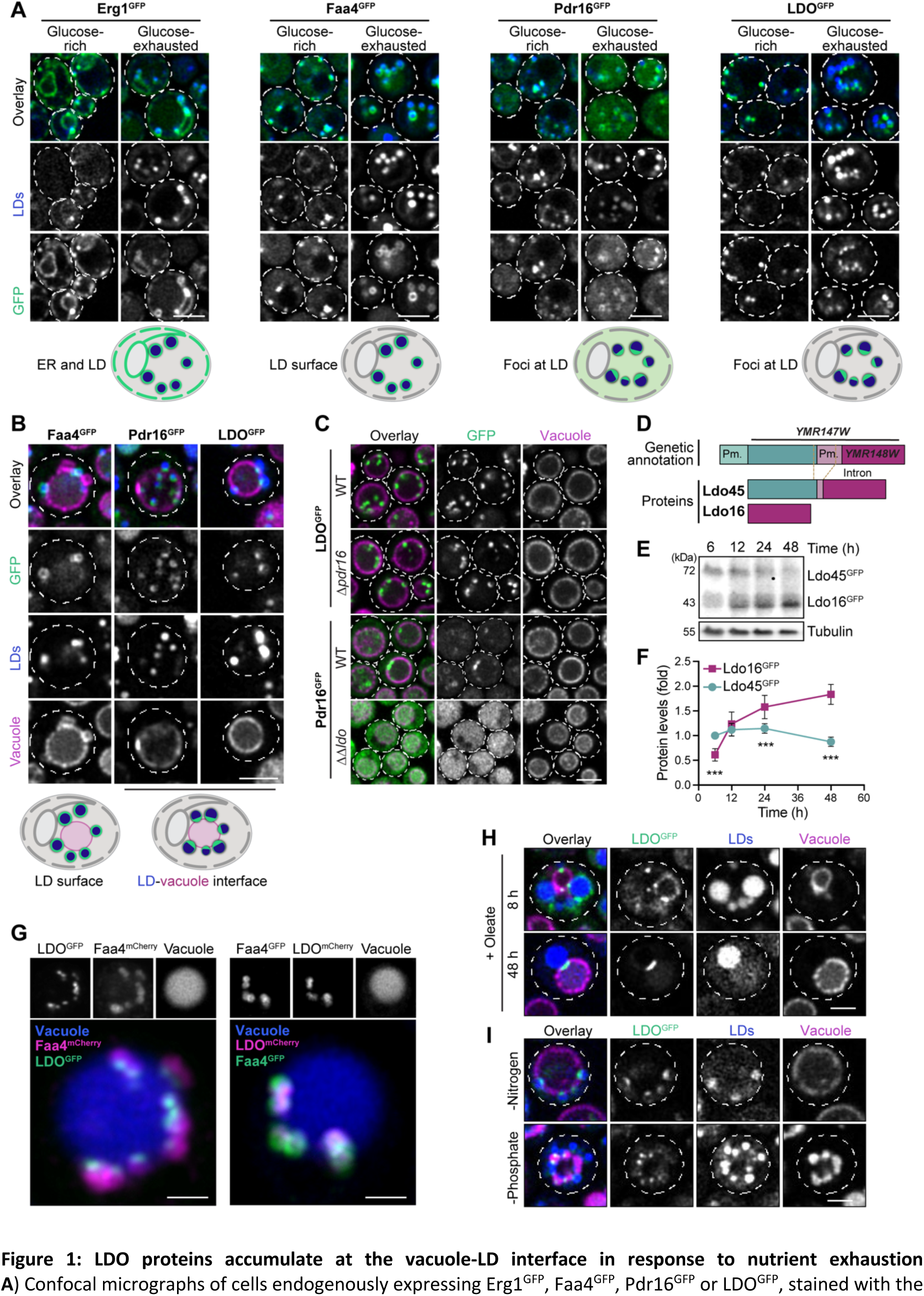
LDO proteins accumulate at the vacuole-LD interface in response to nutrient exhaustion. **A**) Confocal micrographs of cells endogenously expressing Erg1^GFP^, Faa4^GFP^, Pdr16^GFP^ or LDO^GFP^, stained with the LD dye MDH at 8 h (glucose-rich) and 48 h (glucose-exhausted) of culturing in standard glucose media. Scale bars: 3 μm. Schematics depict the subcellular distribution of GFP-tagged proteins at 48 h. **B**) Confocal micrographs of cells expressing LDO^GFP^, Pdr16^GFP^ or Faa4^GFP^ after growth into glucose exhaustion (48 h). Cells were in addition equipped with Vph1^mCherry^ and stained with MDH to visualize the vacuole and LDs, respectively. Scale bar: 3 μm. Schematics depict the distribution of GFP-tagged proteins on LDs and the vacuole. **C**) Confocal micrographs of glucose-exhausted (48 h) wild type (WT) cells and cells lacking either both LDO proteins (ΔΔ*ldo*) or Pdr16 (Δ*pdr16*), expressing either LDO^GFP^ or Pdr16^GFP^ and additionally carrying with Vph1^mCherry^ Scale bar: 3 μm**. D**) Schematic representation of the genomic loci of *YMR147W* and *YMR148W* and the respective promoters (Pm), encoding the LDO proteins Ldo16 and Ldo45. **E, F**) Immunoblot analysis of total protein extracts of cells endogenously expressing LDO^GFP^ collected after 6, 12, 24 and 48 h of culturing. Blots were probed with antibodies directed against GFP and tubulin as loading control. A representative blot (E) and corresponding densitometric quantification of Ldo45^GFP^ and Ldo16^GFP^ levels (F), normalized to tubulin and depicted as fold of Ldo45^GFP^ at 6 h, are shown; Data represent mean ± s.e.m.; *n* = 8. **G**) Z-projection of confocal microscopy stacks showing top view of glucose-exhausted cells (24 h) endogenously expressing either LDO^GFP^ and Faa4^mCherry^ or LDO^mCherry^ and Faa4^GFP^, stained with CMAC to visualize the vacuole. Scale bar: 1 μm. **H**) Confocal micrographs of cells expressing LDO^GFP^ and Vph1^mCherry^ grown for 8 h and 48 h in standard glucose media supplemented with 0.5% oleate or respective solvent control. Cells were stained with MDH to visualize LDs. Scale bar: 2 μm. **I**) Confocal micrographs of cells expressing LDO^GFP^ and Vph1^mCherry^ subjected to either nitrogen or phosphate depletion. Scale bar: 2 μm. *** p < 0.001. See Supplemental Table 4 for details on statistical analyses.

### LDO proteins target LDs to the vacuole to facilitate lipophagy

We directly tested whether LDO proteins are required for vCLIP to form. In wild type cells, LDs were targeted to the vacuolar membrane (visualized via Vph1^mCherry^) shortly after glucose exhaustion. Upon prolonged starvation, a large part of the LD population was gradually engulfed into the vacuole via lipophagy (Fig. 2A). In contrast, such starvation-induced lipophagy was absent in cells lacking both LDO proteins (ΔΔ*ldo*), which have already been proposed to contribute to lipophagy by yet unclear mechanisms (Teixeira et al., 2018). In ΔΔ*ldo* cells, LDs remained clustered at one side of the vacuole, likely reflecting the nER and its contact to the vacuole at the NVJs (Fig. 2A). Indeed, visualization of the NVJs employing Nvj1^GFP^ demonstrated that in glucose-exhausted ΔΔ*ldo* cells, a few LDs remained associated with the NVJs but most LDs clustered at the nER, while in wild type cells they surrounded the vacuole for subsequent microlipophagic engulfment (Fig. 2B; Supplemental Fig. S2A). Transmission electron microscopy demonstrated that in glucose-exhausted wild type cells, more than 80% of LDs were closely attached to the vacuolar membrane at contact sites that reflect vCLIP, often partly or already completely engulfed by the vacuole (Fig. 2C, D). In contrast, in ΔΔ*ldo* cells, only 10% of the LDs contacted the vacuole, while 80% remained associated with the ER, mostly nER (Fig. 2C-E). This suggests a critical function of the LDO proteins in vCLIP formation and supports the notion that contact formation between the LDs and the vacuole is a prerequisite for subsequent microlipophagy. Pdr16 as additional factor that is targeted to vCLIP (Fig. 1A-C) was not required for lipophagy (Supplemental Fig. 2B). To discriminate between Ldo16 and Ldo45 functions, we introduced selective deletion mutations to individually express either Ldo16 or Ldo45 from the chromosomal locus and assessed the subcellular distribution of LDs in glucose-exhausted cells. Confocal microscopy and automated image quantification revealed that the individual loss of Ldo16 (but not of Ldo45) caused a slight reduction of vacuolar LD engulfment, but not to the same extent as the simultaneous lack of both proteins (Fig. 2F, G). Moreover, in ΔΔ*ldo* cells but not the single deletion mutants LDs displayed a different morphology, being fewer in number and prominently enlarged, suggesting that the presence of either LDO protein is sufficient to adjust the number and size of LDs to cellular needs (Fig. 2H, I). In line with this, both Ldo16^GFP^ and Ldo45^GFP^ still efficiently accumulated at vCLIP to establish contact when expressed individually from their chromosomal locus (Fig. 2J). Furthermore, overexpression of either Ldo16 or Ldo45 triggered massive accumulation of LDs inside the vacuole, suggesting that increasing the levels of either one of these proteins efficiently induces lipophagy (Fig. 2K; Supplemental Fig. S2C). As the complete Ldo16 sequence is also present in Ldo45, we created a truncated Ldo45 variant that lacks the part shared with Ldo16. This Ldo45^ΔC148^ mutant was still able to target LDs but failed to attach LDs to the vacuole (Supplemental Fig. S2D, E). Taken together, both LDO proteins establish vCLIP by tethering the LDs to the vacuole via the shared Ldo16 domains, yet Ldo16 expression is selectively upregulated when cells enter stationary phase due to nutrient exhaustion.

**Figure 2:**
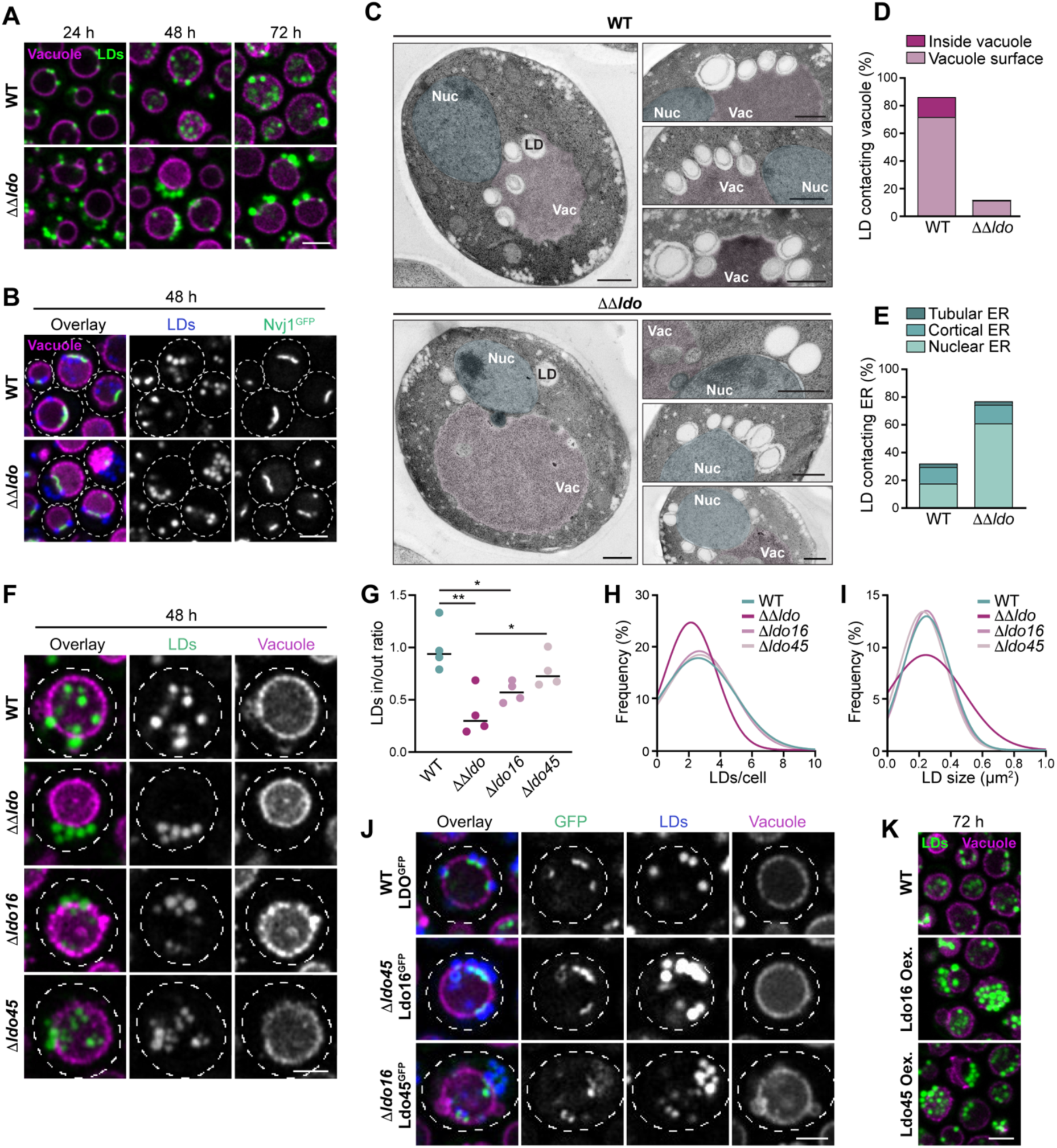
LDO proteins target LDs to the vacuole to facilitate lipophagy. **A)** Confocal micrographs of wild type (WT) cells or cells lacking both LDO proteins equipped with Vph1^mCherry^ to visualize the vacuole after 24, 48 and 72 h of culturing. LDs were stained with BODIPY 493/503. Scale bar: 3 μm. **B)** Confocal micrographs of WT and ΔΔ*ldo* cells endogenously expressing Nvj1^GFP^ and Vph1^mCherry^ after prolonged glucose exhaustion (48 h). LDs were stained with MDH. Scale bar: 3 μm. Corresponding images at 8 h and 24 h are shown in Supplemental Fig. S2A. **C**) Transmission electron micrographs of WT and ΔΔ*ldo* cells grown into glucose depletion (32 h). Colors indicate the vacuole (pink) and the nucleus (blue). Scale bars: 500 μm. **D, E**) Quantification of transmission electron micrographs shown in (C), depicting the number of LDs in contact with the vacuole or engulfed by the vacuole (D) as well as the number of LDs contacting the tubular, cortical or nuclear ER (E). Sections quantified: *n* = 94 (for WT) and *n* = 127 (for ΔΔ*ldo*). **F**) Confocal micrographs of WT and ΔΔ*ldo* cells as well as of the genomic single deletion mutants Δ*ldo16* and Δ*ldo45*, endogenously expressing either Ldo45 or Ldo16. Cells were additionally equipped with Vph1^mCherry^, stained with BODIPY 493/503 and analyzed upon growth into glucose exhaustion (48 h). Scale bar: 2 μm. **G**) Micrographs in (F) were used for automated quantification of the ratio of the BODIPY intensity inside and outside of the segmented vacuole, indicating lipophagic sequestration of LDs. Line represents mean; *n* = 4, with 200-500 cells quantified per *n*. (**H, I**) Micrographs in (F) were used for automated quantification of the number of LDs per cell frequencies (H) and of LD size (μm^2^) frequencies (I) fitted to a gaussian distribution. Between 800-1500 cells were evaluated per genotype. **J**) Confocal micrographs WT cells expressing LDO^GFP^ as well as the genomic single deletion mutants Δ*ldo45* and Δ*ldo16* endogenously expressing either Ldo16^GFP^ or Ldo45^GFP^. Cells were additionally equipped with Vph1^mCherry^, stained with MDH and analyzed upon growth into glucose exhaustion (48 h). Scale bar: 2 μm. **K**) Confocal micrographs of WT cells and ΔΔ*ldo* cells ectopically overexpressing either Ldo45 or Ldo16. Cells were additionally equipped with Vph1^mCherry^, stained with BODIPY 493/503 and analyzed upon prolonged glucose exhaustion (72 h). Scale bar: 3 μm. See Supplemental Fig. S2C for corresponding single channel micrographs. * p < 0.05 and ** p < 0.01. See Supplemental Table 4 for details on statistical analyses.

### The C-terminal disordered region of Ldo16 anchors LDs to the vacuole

Next, we generated a set of Ldo16 mutants to assign function to distinct protein regions. The N-terminus of Ldo16 likely corresponds to hydrophobic transmembrane α-helixes (Fig. 3A) and includes a proline-lysine-lysine-glutamine (PLLG) motif as typical helix breaker with ∼20 hydrophobics on either side. This might facilitate hairpin-like membrane insertion, characteristic for LD proteins that are targeted to LDs from the ER membrane (Kory et al., 2016; Olarte et al., 2022). In addition, Ldo16 is predicted to contain a cationic amphipathic helix (CAH; Fig. 3B), again a common motif to target both ER membrane-inserted and cytosolic, soluble proteins to LDs (Pataki et al., 2018; Prévost et al., 2018). The remainder of Ldo16 in its C-terminus is mostly predicted to be an intrinsically disordered region, ending with a short α-helix (Fig. 3C). We created a series of C- and N-terminally truncated Ldo16 variants as well as point mutants within the CAH, all expressed as GFP fusion proteins (Fig. 3C-E). Immunoblotting confirmed the expression of all Ldo16 mutants, though modification of the predicted CAH impaired protein stability (Fig. 3D). Assessing the subcellular localization of the different mutants demonstrated that the N-terminal hydrophobic region served as ER targeting signal, as its deletion prevented ER import (Ldo16^ΔN49^ and Ldo16^ΔN72^). In agreement with this, the Ldo16 variant containing only the N-terminal hydrophobic helixes (Ldo16^ΔC98^) was still imported into the ER. Importantly, this mutant failed to associate with LDs, while a slightly longer variant that still contained the CAH (Ldo16^ΔC54^) was fully redirected to LDs, demonstrating that the CAH is critical to target Ldo16 from the ER membrane to LDs in the so-called ERTOLD pathway (Fig. 3C) (Olarte et al., 2022). Exchanging 5 hydrophobic residues in the CAH with alanine did not prevent LD targeting, though accumulation of this Ldo16^5xA^ variant at the contact site was reduced (Fig. 3C, E). However, a stronger mutation of the CAH by inserting 2 glutamates resulted in severely compromised protein stability (Fig. 3D) and retention of the protein in the ER (Fig. 3C), supporting the notion that amphipathicity of the CAH contributes to Ldo16 targeting to LDs (Fig. 3C). Importantly, Ldo16^ΔC54^, which harbors a functional CAH but lacks the C-terminal disordered region, was efficiently redirected to LDs but decorated the complete LD surface instead of accumulating at vCLIP. In addition, the complete LD population remained attached to one side of the vacuole, likely reflecting the nER (Fig. 3C). These results demonstrate that the C-terminus of Ldo16 is critical for contact formation. Truncation of only 24 residues at the very C-terminus, removing only the short α-helix, did not prevent accumulation at vCLIP, indicating that it is the intrinsically disordered region that is essential for Ldo16 function in contact site formation (Fig. 3C, E). Collectively, this suggests that Ldo16 is targeted to the ER via its N-terminal hydrophobic region, is re-directed from the ER membrane to the LD surface via its CAH and attaches to the vacuolar membrane via the intrinsically disordered region in its C-terminus to establish vCLIP (Fig. 3F, G).

**Figure 3:**
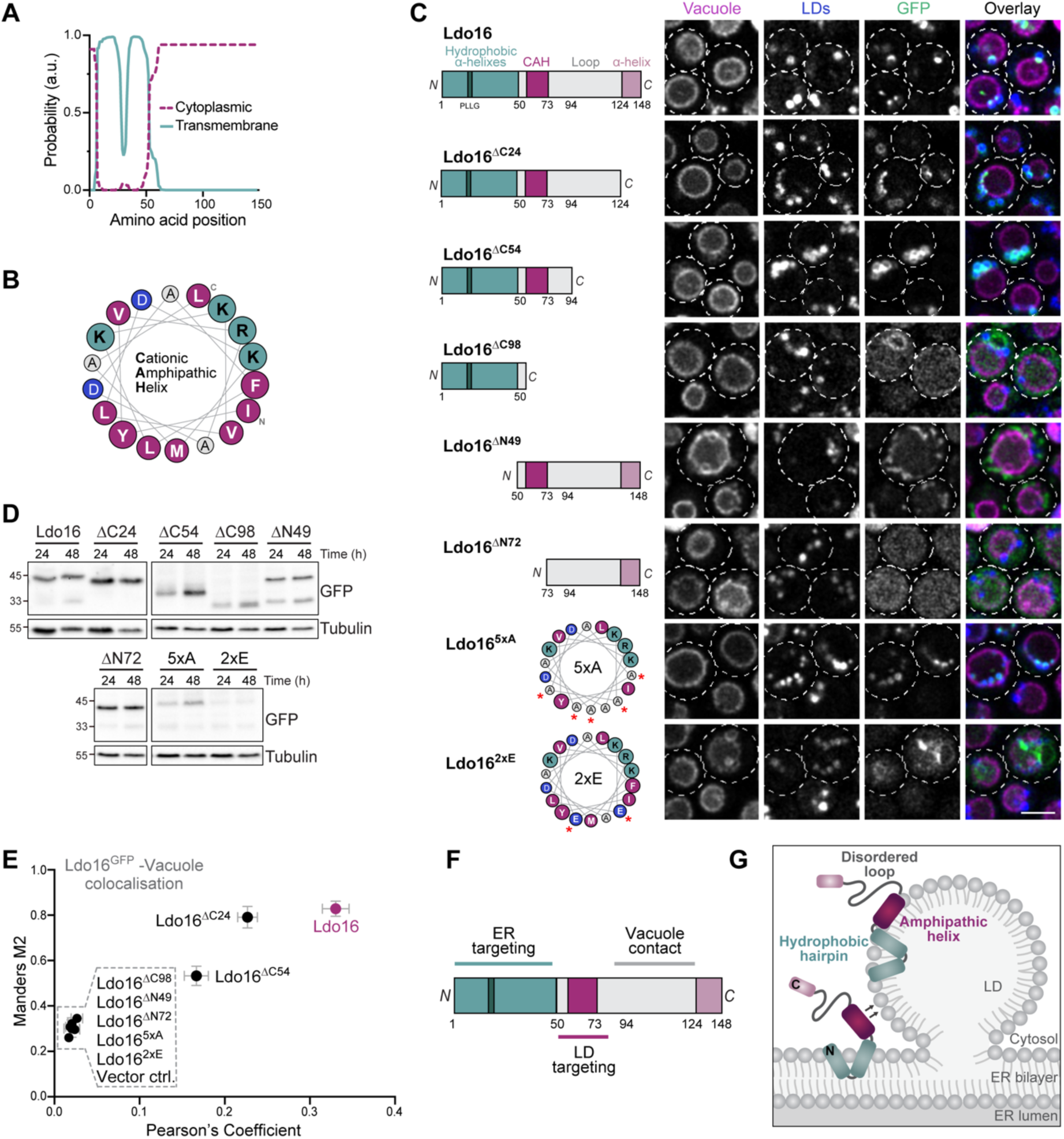
The C-terminal disordered region of Ldo16 anchors LDs to the vacuole. **A**) Hydrophobicity plot of the Ldo16 protein sequence created with Phobius. Probability of transmembrane or cytoplasmic topology per residue is depicted. **B**) α-Helix prediction by HeliQuest of putative Ldo16 cationic amphipathic helix (CAH). Hydrophobic residues are shown in magenta, positively charged residues in turquoise. **C)** Schematic representation of the generated Ldo16 mutants as well as confocal micrographs upon ectopic expression of the GFP-tagged Ldo16 mutants in ΔΔ*ldo* cells equipped with Vph1^mCherry^ and stained with MDH, analyzed after 48 h of culturing. Scale bar: 3 μm. **D**) Representative immunoblot of total protein extracts from ΔΔ*ldo* cells ectopically expressing Ldo16^GFP^ or indicated Ldo16 mutants, collected after 24 h and 48 h of growth. Membranes were probed with antibodies directed against GFP and tubulin. **E)** Colocalization analysis of wild type Ldo16^GFP^ and indicated GFP-tagged Ldo16 mutants with the vacuole (Vph1^mCherry^) from micrographs shown in (D). Manders M2 coefficient was plotted against Pearson’s coefficient. Data represent mean ± s.e.m.; *n* = 6-10, with at least 30 cells per *n*. **F, G)** Schematics of the Ldo16 domains required for ER import, LD targeting and contact formation with the vacuole (F) and of Ldo16 integration into the monolayer of a LD (G).

### Vac8 is required for vCLIP formation

To identify cellular processes and molecular determinants involved in Ldo16-mediated tethering of LDs to the vacuole, we examined LDO^GFP^ localization in a set of mutants with established functions in LD biosynthesis, macro- and microautophagy, the ESCRT machinery, NVJ formation and vacuolar membrane lipid composition, which included microdomain formation and phosphatidylinositol phosphate (PIP) metabolism. All mutants, expressing Vph1^mCherry^ to visualize the vacuole and stained with the LD dye MDH, were grown to glucose exhaustion and assessed microscopically for contact formation (Fig. 4A; Supplemental Fig. S3), scored as colocalization between LDO^GFP^ and Vph1^mCherry^ (Fig. 4B). The LDO proteins have previously been shown to interact with seipin (Sei1) at the ER membrane to support LD biogenesis in growing cells (Eisenberg-Bord et al., 2018; Teixeira et al., 2018). However, the lack of seipin only affected LD size and number but did not compromise LDO^GFP^ targeting to the contact sites. Likewise, neither genetic inactivation of autophagy nor impairment of endosomal protein sorting via deletion of genes coding for ESCRT components, which have been suggested to contribute to lipophagy (Garcia et al., 2021; Liao et al., 2021; Seo et al., 2017; van Zutphen et al., 2014), altered LDO-mediated tethering of LDs to the vacuole. The formation of microdomains on the vacuolar membrane has also been associated with stationary phase lipophagy (Liao et al., 2021; Wang et al., 2014), but defective microdomain formation, for instance due to compromised sterol incorporation into membranes due to the lack of the Niemann-Pick proteins Npc2 and Npr1 (Tsuji et al., 2017), did not prevent vCLIP formation. Similarly, changes in phosphatidylinositol metabolism had no effect on contact formation despite alterations in vacuolar morphology (Fig. 4A, B; Supplemental Fig. S3). Finally, we tested whether NVJs are required for LD tethering to the vacuole, as the LDO proteins have previously been shown to specifically decorate a LD subpopulation associated with the NVJs (Eisenberg-Bord et al., 2018). LDO-mediated tethering was unaffected in cells lacking either Nvj1 or Mdm1, bridging proteins required for formation of NVJs (Henne et al., 2015; Pan et al., 2000), or lacking the NVJ regulator Snd3 (Tosal-Castano et al., 2021). Importantly, one strain showed severely impaired vCLIP formation: the loss of Vac8 prevented LDO^GFP^ targeting to the vacuolar-LD interface, resulting instead in its spreading over the complete LD surface (Fig. 4C). Quantification of LDO^GFP^-Vph1^mCherry^ colocalization supported the absence of vCLIP in Δ*vac8* cells (Fig. 4B, D). In line with a critical function of Vac8 in recruiting LDs to the vacuole, cells lacking the palmitoyl transferase Pfa3, which lipidates Vac8 to attach it to the vacuolar membrane (Hou et al., 2005), resembled Δ*vac8* cells. LDO^GFP^ covered the complete LD surface, and LDO^GFP^-Vph1^mCherry^ colocalization was strongly decreased (Fig. 4C, D). The armadillo repeat protein Vac8 has previously been linked to lipophagy (van Zutphen et al., 2014) and contributes to multiple cellular processes via direct interaction with distinct proteins, including Nvj1 to tether the vacuole to the nER at NVJ (Pan et al., 2000), Vac17 to facilitate vacuole inheritance (Tang et al., 2003), as well as Atg11 and Atg13 to support phagophore assembly during selective and bulk autophagy, respectively (Hollenstein et al., 2021, 2019). Still, the lack of any of these Vac8 interactors did not preclude vCLIP formation, despite clear vacuolar defects such as massive vacuolar fragmentation in Δ*vac17* cells (Fig. 4F). This suggest that it is the absence of Vac8 *per se* rather than the resulting impairment of associated cellular processes that prevents the formation of vCLIP. Overall, these data demonstrate that Vac8 is required to recruit LDs to the vacuole.

**Figure 4:**
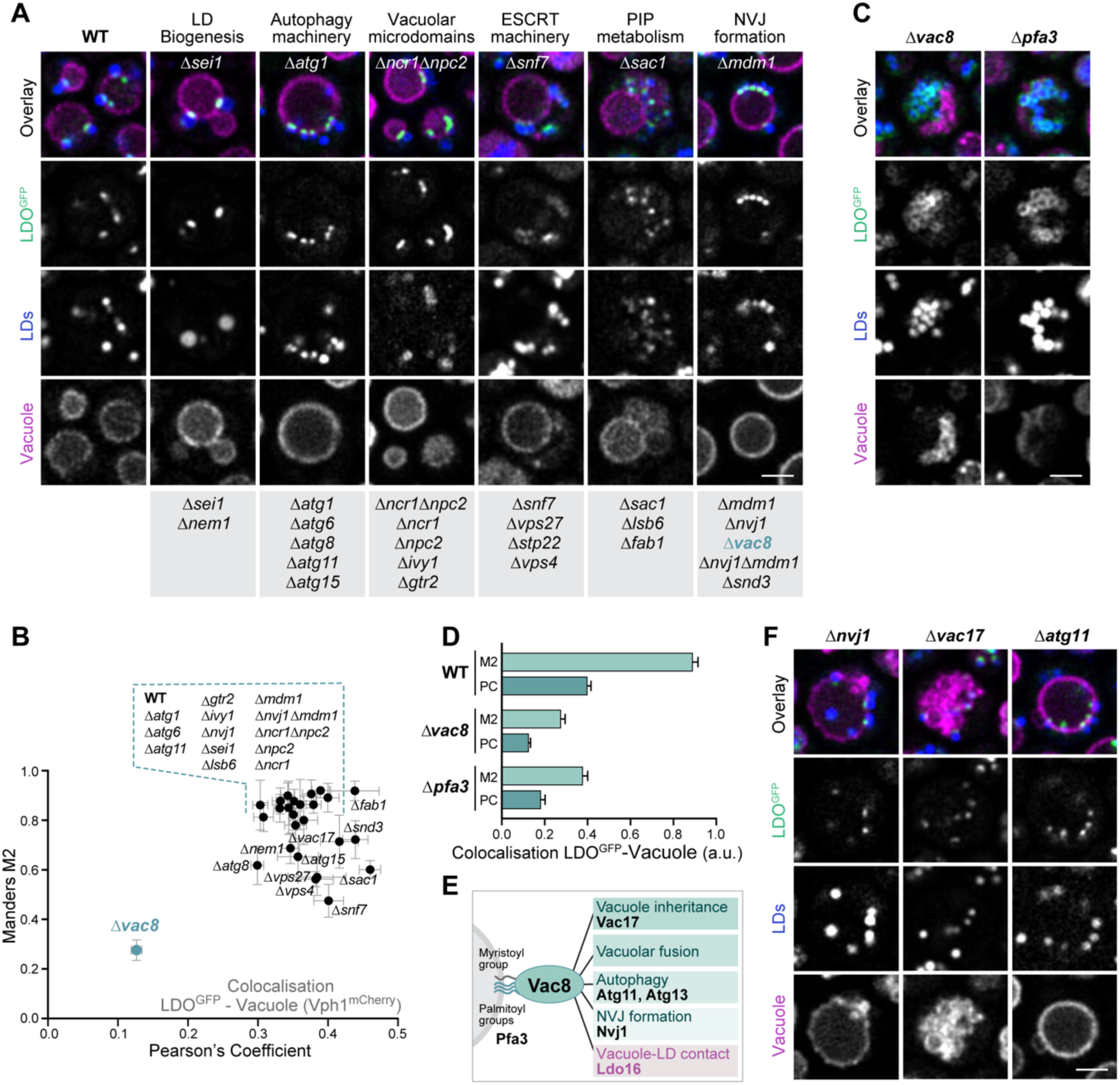
Vac8 is required for vCLIP formation. **A, B)** Confocal micrographs of wild type (WT) cells and indicated deletion mutants endogenously expressing LDO^GFP^ and Vph1^mCherry^, stained with MDH and analyzed upon growth into glucose exhaustion (48 h) (A) as well as corresponding colocalization analysis of Vph1^mCherry^ (vacuole) with LDO^GFP^. Manders M2 coefficient was plotted against Pearson’s coefficient. Data represent mean ± s.e.m.; *n* = 4-10, with at least 30 cells per *n*. Micrographs for the deletion mutants not shown but listed in (A) are shown in Supplemental Fig. S3. Scale bar: 2 μm. **C**) Confocal micrographs of Δ*vac8* and Δ*pfa3* cells expressing LDO^GFP^ and Vph1^mCherry^, stained with MDH and analyzed at 48 h. Scale bar: 2 μm. **D)** Colocalization analysis of Vph1^mCherry^ (vacuole) with LDO^GFP^ corresponding to micrographs shown in (C). Manders M2 coefficient and Pearson’s coefficient were plotted. Data represent mean ± s.e.m.; *n* = 5-10, with at least 30 cells per *n*. **E)** Schematic representation of Vac8 functions and associated interaction partners. **F)** Confocal micrographs of cells lacking selected Vac8 interacting proteins and endogenously expressing LDO^GFP^ and Vph1^mCherry^, stained with MDH and analyzed upon growth into glucose exhaustion (48 h). Scale bar: 2 μm.

### Modification of the C-terminus of Vac8 disrupts vCLIP formation

To test for co-localization between the LDO proteins and Vac8 at the vacuole-LD interface, we monitored the subcellular distribution of endogenously expressed LDO^GFP^ and Vac8^mScarlet^ (Fig. 5A). C-terminal tagging of Vac8 has extensively been used in the field, mostly without any impairment of Vac8 functions (Pan and Goldfarb, 1998), and we confirmed that mScarlet-tagging resulted in a functional protein based on growth assays (Supplemental Fig. S4A, B). Interestingly, the C-terminal tagging of Vac8 triggered the redistribution of LDO^GFP^ to cover the complete LD surface, a phenotype reminiscent of its distribution in Δ*vac8* cells (Fig. 5A and 4C). Formation of vCLIP was absent and the LDs failed to surround the vacuole (Fig. 5A). The use of HA as alternative epitope resulted in a similar disruption of contact formation and the localization of LDO^GFP^ over the complete LD surface (Supplemental Fig. S4C), while N-terminal tagging caused the loss of Vac8, most likely due to interference with its palmitoylation and thus defective anchoring to the vacuolar membrane (Fig. 5A). The interactions of Vac8 with Nvj1 and Atg13 have previously been studied at the structural level, showing that the last armadillo repeat at its C-terminus (ARM12) is not involved in the interaction (Jeong et al., 2017; Park et al., 2020). As C-terminal tagging selectively disrupted vCLIP formation in glucose-exhausted cells while not affecting other Vac8 functions, we created a series of genomically encoded C-terminal truncations of Vac8, successively deleting the α-helixes of ARM12 (Fig. 5B). We monitored LDO^GFP^ localization in combination with the vacuole (using FM4-64) and LDs (using MDH) (Fig. 5C) and scored for LDO^GFP^ – vacuole colocalisation (Fig. 5D) as well as for accumulation of LDO^GFP^ at the contact sites *versus* distribution across the entire LD surface (Fig. 5E). The truncation of the last α-helix (α3) only slightly compromised contact formation (Fig. 5D, E), although vacuolar morphology was already affected, leading to the formation of vacuolar sheets and compartments that partially surrounded the LDs (Fig. 5C). Further truncation of also α2 or the complete ARM12 disrupted contact formation, resulting in the spreading of LDO^GFP^ across the complete LD surface reminiscent of Δ*vac8* cells (Fig. 5C-E). In parallel, also vacuolar morphology with these truncations resembled the characteristic vacuolar fragmentation of cells completely lacking Vac8 (Fig. 5C). To assess whether these C-terminal truncations of Vac8 would also interfere with well-established interactions, we used Nvj1^GFP^ to monitor NVJ formation in cells harboring the Vac8 variants at the endogenous locus. This revealed a similar pattern of disruption of Vac8 binding to its partner, here Nvj1, with the complete loss of ARM12 causing a loss of NVJ formation and the redistribution of Nvj1 to the whole nER (Fig. 5F; Supplemental Fig. S4D). The progressive loss of Vac8 interactions upon truncation of the α-helixes of ARM12 correlated well with the extent of growth arrest (Fig. 5G; Supplemental Fig. S4E). As ARM12 does not directly contribute to the interaction interface of the Vac8-Nvj1 complex, determined by crystallography (Jeong et al., 2017; Park et al., 2020), it is likely that these C-terminal truncations impair a general aspect of protein structure. The regions of both Nvj1 and Atg13 that interact with Vac8 are disordered loops near their C-termini (Jeong et al., 2017; Park et al., 2020). Despite these sharing little sequence homology, both loops run as extended peptides along the minor groove of the armadillo repeat superhelix, anti-parallel to the armadillo repeats. We used ColabFold (Mirdita et al., 2022) to examine if an interaction surface in Vac8 could be predicted for Ldo16. Of the five best models obtained from full-length sequences of both proteins, all showed a portion of the disordered C-terminus of Ldo16 in contact with the minor groove of the Vac8 superhelix (Figure 5H). The extent of contact increased with rank of the model; with the top ranked model having the majority of the C-terminus of Ldo16 making multiple contacts with Vac8 (22 of the 34 residues between 104–137 had ≥8 contacts defined as atom centres ≤ 4Å apart). Notably, the predicted position of Ldo16 is the same as that previously found for Nvj1 (Fig. 5I). Thus, Ldo16 and Nvj1 might bind to the same groove of Vac8, potentially competing for Vac8 interaction when Ldo16-decorated LDs enrich at the NVJs shortly after glucose exhaustion. The elongated interaction surface along the minor groove of Vac8 is well conserved despite being spread across multiple ARM repeats (Figure 5J). The five ColabFold models aligned Ldo16 in the same direction, which was parallel to Vac8 (*i.e*., opposite to the direction of Nvj1 and Atg13). According to this prediction, the C-termini of Vac8 and Ldo16 would be in close proximity, and insertion of C-terminal tags simultaneously on both proteins might interfere with their interaction. To test this, we monitored LD localization in a strain equipped with Vac8^mScarlet^ and untagged LDO proteins. As opposed to the finding with both proteins tagged, Vac8^mScarlet^ indeed accumulated at sites where LDs made contact with the vacuole when the LDO proteins were untagged (Fig. 5K, L). Overall, these results suggest that (i) Vac8 is recruited to vCLIP and (ii) C-terminal tagging of both proteins at the same time interferes with vCLIP formation. Clustering of Vac8 at LDs was absent in ΔΔ*ldo* cells, while clear Vac8 patches at NVJs were visible (Fig. 5K), demonstrating that the LDO proteins are critical for Vac8 recruitment to the vacuole-LD contact sites.

**Figure 5:**
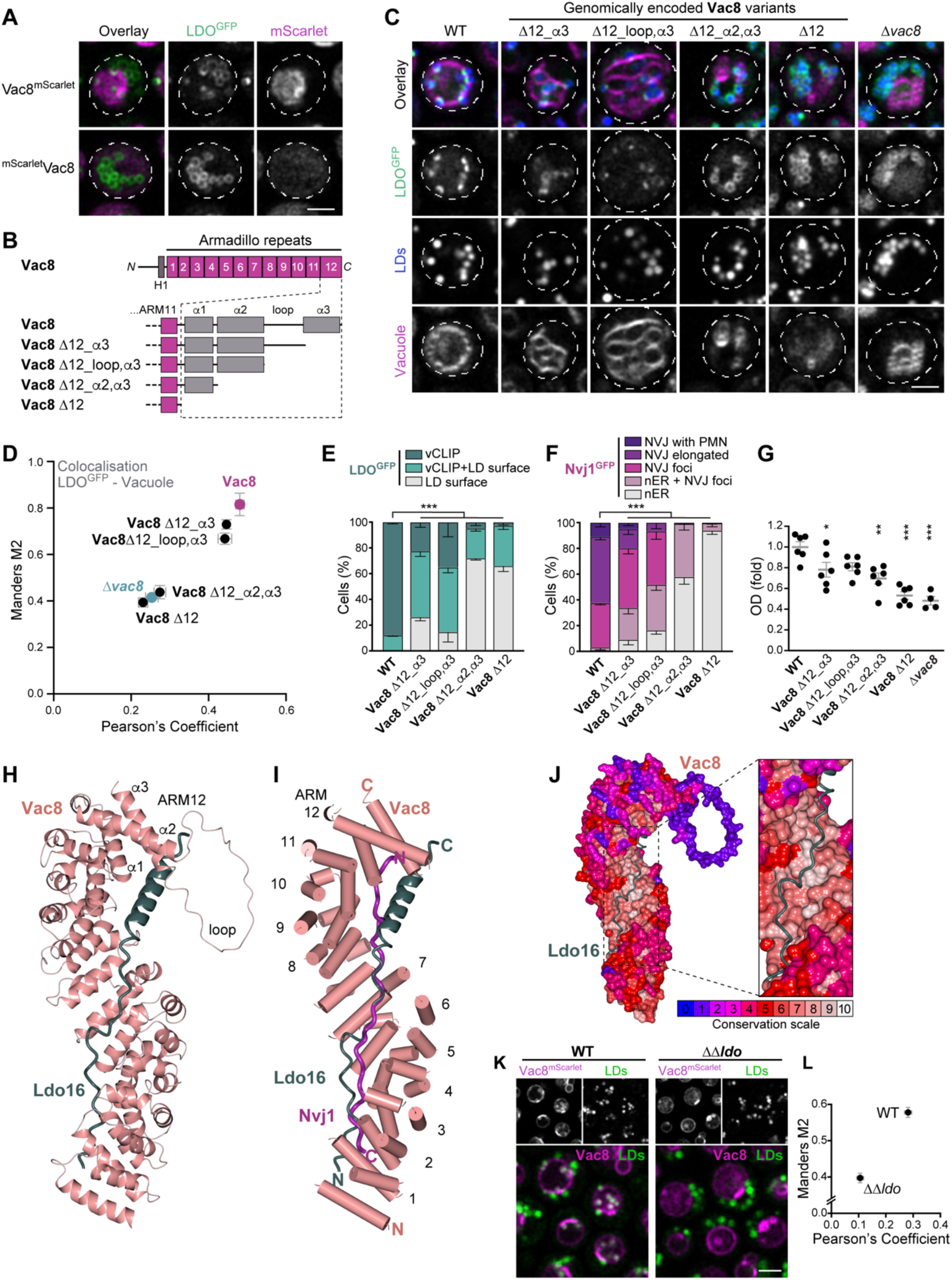
Modification of the C-terminus of Vac8 disrupts vCLIP formation. **A)** Confocal micrographs of cells expressing LDO^GFP^ and Vac8, C-terminally or N-terminally fused to mScarlet, after 48 h of culturing. Scale bar: 2 μm. **B)** Schematic representation of the armadillo (ARM) repeat protein Vac8, highlighting the ARM12 domain structure, as well as the generated Vac8 mutants, partly or completely lacking ARM12. **C)** Confocal micrographs of WT and Δ*vac8* cells as well as cells endogenously expressing the C-terminally truncated Vac8 variants shown in (B), additionally equipped with LDO^GFP^ and stained with MDH as well as with FM 4-64 to visualize the vacuolar membrane. Scale bar: 2 μm. **D)** Colocalization analysis of FM4-64 (vacuole) with LDO^GFP^ in cells endogenously expressing indicated Vac8 mutants as shown in (C). Manders M2 coefficient was plotted against Pearson’s coefficient, and data represent mean ± s.e.m.; *n* = 5-9, with at least 30 cells per *n*. **E)** Quantification of cells endogenously expressing indicated Vac8 mutants according to LDO^GFP^ distribution from micrographs shown in (C). At least 230 cells were evaluated per genotype. **F)** Quantification of cells endogenously expressing indicated Vac8 mutants according to Nvj1^GFP^ distribution, categorized as NVJ with piecemeal microautophagy of the nucleus (PMN), NVJ elongated, NVJ foci, nER +NVJ foci and nER from micrographs shown in Supplemental Fig. S4D. At least 90 cells were evaluated per genotype. **G)** Optical density, indicative of cellular growth, of WT and Δ*vac8* cells as well as cells carrying indicated Vac8 mutants, after 10 h of culturing. Data show mean ± s.e.m.; *n* = 6. Asterisks indicate significant differences compared to WT. See Supplemental Fig. S4E for corresponding growth curves. **H)** Predicted interaction model using the full-length sequences of Vac8 and Ldo16, created with ColabFold. **I)** Simplified predicted position of Ldo16 compared to the position previously found for Nvj1, spread across multiple ARM repeats. **J)** Structural model of Vac8 depicted in conservation scale to highlight the high degree of conservation of the minor groove of Vac8, the predicted interaction surface of Ldo16. **K)** Confocal micrographs of WT and ΔΔ*ldo* cells expressing Vac8^mScarlet^ and stained with BODIPY 493/503, analyzed upon glucose exhaustion (48 h). Scale bar: 2 μm. **L)** Colocalization analysis of Vac8^mSacrlet^ with LDs in WT and ΔΔ*ldo* cells corresponding to micrographs shown in (K). Manders M2 coefficient was plotted against Pearson’s coefficient. Data represent mean ± s.e.m.; *n* = 10-11, with at least 25 cells per *n*. * p < 0.05, ** p < 0.01 and *** p < 0.001. See Supplemental Table 4 for details on statistical analyses.

### Vac8 recruits LDs to cellular membranes

Next, we used electron microscopy and immunogold labeling to test for a specific targeting of native Vac8 to vCLIP in glucose-exhausted wild type cells. As expected, we detected Vac8 at the entire vacuolar membrane, with an enrichment at the interface between the vacuole and the nER at NVJs (Fig. 6A, B). Of note, Vac8 in addition clustered at the vacuolar membrane regions that made contact with LDs and that often were in the process of engulfing LDs, suggesting that Vac8 indeed accumulates at vCLIP (Fig. 6A, B). To test if Vac8 serves as a tether protein that recruits LDs to cellular membranes, we reconstituted this process at the nER, employing a previously established tethering system (Hollenstein et al., 2021). Fusion of the N-terminal half of Nvj1 to Vac8ΔN, a variant that lacks the domains necessary to attach it to the vacuolar membrane, drives the attachment of Vac8ΔN to the cytosolic phase of the nuclear envelope (Hollenstein et al., 2021). Importantly, ectopic expression of this nER-attached Vac8 in absence of endogenous, vacuolar Vac8 (Δ*vac8* cells) recruited LDs to the nER (Fig. 6C). The LDO proteins and Vac8 accumulated at concave nER patches in which attached LDs were buried, suggesting that Vac8 is the single necessary and sufficient component of the vacuolar membrane that recruits LDs (Fig. 6C, D). As a positive control to show the functionality of the chimera, Vac8ΔN was fused to Vph1 to retain it at the vacuolar membrane, which predicably resulted in colocalization of LDO^GFP^ and Vac8 at the vacuolar membrane and LD attachment to the vacuole (Fig. 6C, D). Collectively, these data suggest that Vac8 interacts with LDO proteins to establish vCLIP and is sufficient to recruit LDs to an organelle, thus fulfilling the formal criteria of a membrane contact site tether (Scorrano et al., 2019).

**Figure 6:**
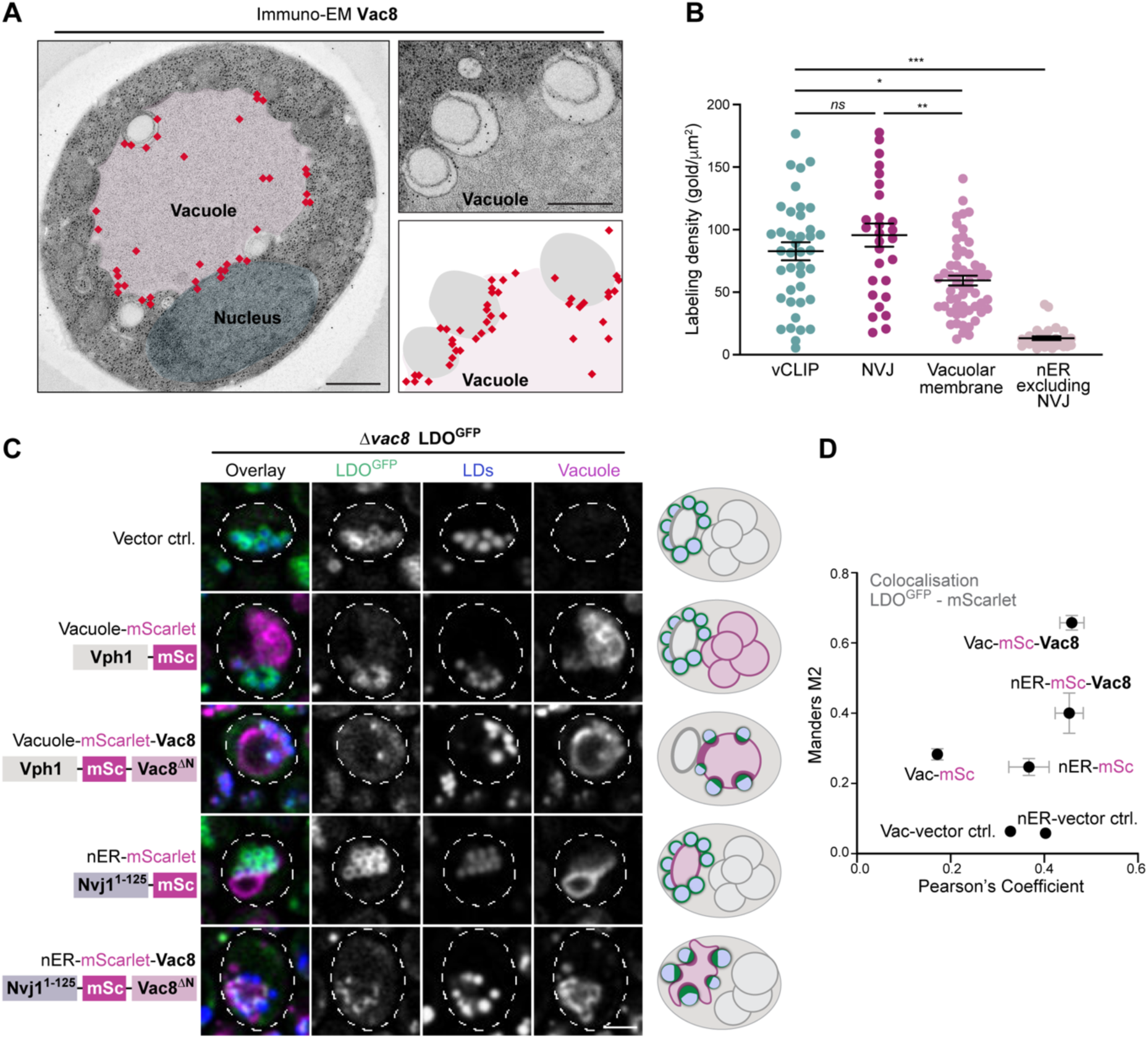
Vac8 recruits LDs to cellular membranes. **A)** Immunogold labelling of electron micrographs of glucose-exhausted WT cells (32 h) using a Vac8 antibody. Gold particles are indicated in red, and a respective labeling density model is shown. Scale bars: 500 μm. **B)** Quantification of Vac8 labeling density corresponding to electron micrographs shown in (A). Labeling density was quantified at the vacuolar membrane in direct contact with LDs (vCLIP), in direct contact with the nER (NVJs) and not in contact with other organelles (vacuolar membrane) as well as at the nER excluding the NVJ region. Individual data points, mean (line) and s.e.m. are shown. In total, 57 sections displaying vacuoles were quantified, with varying numbers of data points for the 4 different categories of membrane regions, depending on their presence in respective section. **C)** Confocal micrographs of Δ*vac8* cells endogenously expressing LDO^GFP^ and ectopically expressing depicted constructs, in which a cytosolic version of Vac8 (Vac8ΔN) was fused to either Vph1^mScarlet^, anchoring it to the vacuole, or truncated Nvj1^mScarlet^, anchoring it to the nER. Same constructs without Vac8ΔN served as controls. Observed cellular distribution of Vac8 and LDO is represented in the schematics. Scale bar: 2 μm. **D)** Colocalization analysis of LDO^GFP^ with the mScarlet signal, corresponding to micrographs shown in (C). Manders M2 coefficient was plotted against Pearson’s coefficient. Data represent mean ± s.e.m.; *n* = 6, with at least 30 cells per *n*.

### Proteolysis associated with lipophagy depends on vCLIP

We quantitatively assessed the impact of defective vCLIP formation on lipophagic activity associated with characteristic proteolytic breakdown of vacuolar cargo. To this end, GFP liberation from Faa4^GFP^ by vacuolar proteases was followed by immunoblotting. Faa4^GFP^ levels progressively decreased with time spent in glucose exhaustion, which was accompanied by the simultaneous increase of free GFP (Fig. 7A-C). Genetic ablation of the LDO proteins reduced the level of free GFP but not Faa4^GFP^, suggesting a drop in lipophagic activity (Fig. 7A-C). Notably, no GFP liberation was detectable in cells devoid of Vac8 (Fig. 7D), demonstrating that the absence of contact formation between LDs and the vacuole, achieved by disrupting Vac8, precluded lipophagic activity. Consistently, Δ*vac8* cells did not sequester LDs into the vacuole and displayed an increase in LD size comparable to ΔΔ*ldo* cells (Fig. 7E-G). Thus, anchoring LDs to the vacuole at vCLIP is required for subsequent lipophagy.

**Figure 7:**
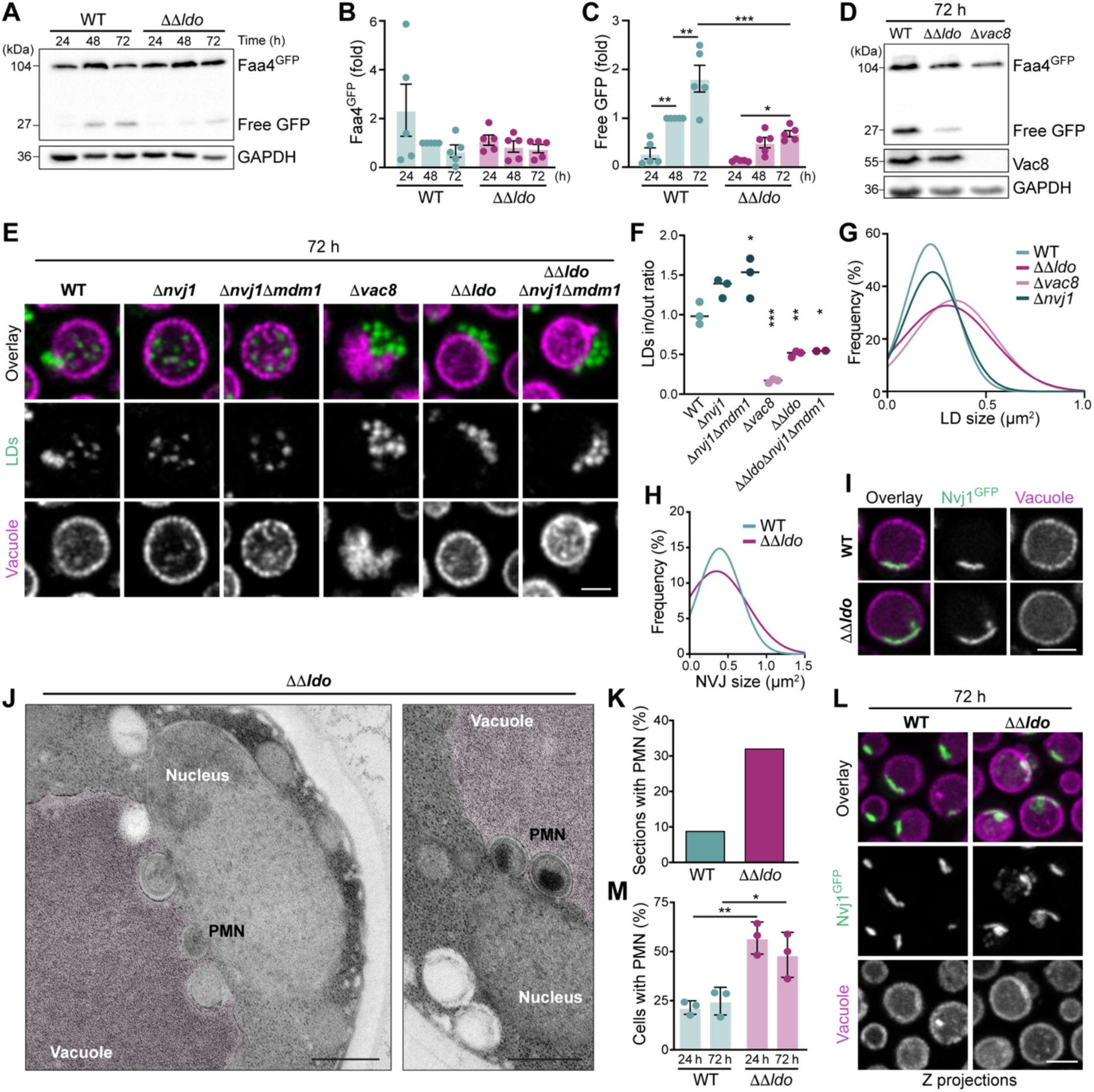
Disruption of vCLIP induces NVJ expansion and piecemeal microautophagy of the nucleus. **A-C)** Immunoblot analysis of total protein extracts from WT and ΔΔ*ldo* cells endogenously expressing Faa4^GFP^ collected after 24, 48 and 72 h of culturing. Membranes were probed with antibodies directed against GFP and GAPDH as loading control. Representative immunoblots (A) as well as corresponding densitometric quantification of full length Faa4^GFP^ (B) and free GFP liberated from Faa4^GFP^ (C), all normalized to GAPDH, are shown. Data represent mean ± s.e.m.; *n* = 6. **D)** Immunoblot analysis of total protein extracts from WT, ΔΔ*ldo*, and Δ*vac8* cells endogenously expressing Faa4^GFP^ collected after 72 h of culturing. Membranes were probed with antibodies directed against GFP, Vac8 and GAPDH as loading control. **E)** Confocal micrographs of WT and indicated deletion mutants expressing Vph1^mCherry^ and stained with BODIPY 493/503 after prolonged glucose exhaustion (72 h). Scale bar: 2 μm. **F)** Micrographs in (E) were used for quantification of the ratio of BODIPY intensity inside and outside of the segmented vacuole, indicating lipophagic sequestration of LDs. Line represents mean; *n* = 3, with at least 120 cells quantified per *n*. Asterisks indicate significant differences from WT. **G)** Micrographs in (G) were used for quantification of LD size (μm^2^) frequencies fitted to a gaussian distribution. At least 450 cells were evaluated per genotype. **H, I)** Microscopic analysis of NVJ in WT and cells endogenously expressing Nvj1^GFP^ and Vph1^mScarlet^. Quantification of NVJ size (μm^2^) frequencies fitted to a gaussian distribution (H) and representative confocal micrographs (I) are shown. Between 400-500 cells were evaluated per genotype. Scale bar: 2 μm. **J, K)** Electron micrographs of glucose-exhausted ΔΔ*ldo* cells (32 h) showing piecemeal microautophagy of the nucleus (PMN) (J) as well as corresponding quantification of sections from WT and ΔΔ*ldo* cells with PMN vesicles (K). Only sections with clearly defined vacuole and nucleus were evaluated. *n* = 56 (for WT) and *n* = 84 (for ΔΔ*ldo*). Scale bars: 500 μm. **L, M)** Z-projections from confocal microscopy stacks of WT and ΔΔ*ldo* cells expressing Nvj1^GFP^ and Vph1^mCherry^ analyzed after 72 h of culturing (L) as well as corresponding quantification of cells with PMN vesicles after 24 h and 72 h of culturing (M). Individual data points and mean ± s.e.m. is shown; *n* = 3, with at least 30 cells evaluated per *n*. Scale bar: 3 μm. * p < 0.05, ** p < 0.01 and *** p < 0.001. See Supplemental Table 4 for details on statistical analyses.

### Disruption of vCLIP induces NVJ expansion and piecemeal microautophagy of the nucleus

Vac8 is a key resident of both NVJs and vCLIP, raising the possibility of cross-talk via this shared and perhaps limiting component. Indeed, inhibition of NVJ formation by removing Nvj1 or both Nvj1 and Mdm1 resulted in increased uptake of LDs into the vacuole (Fig. 7E-F). Additional inactivation of Vac8 blocked this uptake. *Vice versa*, inactivating vCLIP in ΔΔ*ldo* cells affected NVJ size and shape, resulting in expanded NVJs that frequently appeared fragmented due to ongoing piecemeal microautophagy of the nucleus (PMN) (Fig. 7H, I). This special form of microautophagy occurs only at the NVJs and is characterized by vacuolar membrane invagination that leads to the pinching-off of vesicles carrying portions of nuclear cargo for subsequent degradation in the vacuole (Roberts et al., 2003). Electron microscopy demonstrated a 3-fold increase of sections with PMN vesicles upon disruption of vCLIP formation, and often multiple PMN vesicles formed simultaneously along the expanded NVJs (Fig. 7J, K). Confocal microscopy using Nvj1^GFP^ as marker for NVJs and PMN vesicles confirmed a prominent increase of PMN in cells lacking the LDO proteins (Fig. 7L, M). Conceptually, the mutually negative regulation of NVJs and vCLIP is likely caused by a thug-of-war for limiting Vac8, but may also involve compensatory cellular responses to maintain needed lipid flux to the vacuole. Collectively, these results point to a functional link between the NVJ and the vCLIP, both of which are established by Vac8 at the vacuolar membrane and metabolically regulated.

## Discussion

The mobilization of fat stored in LDs plays a central role in both physiological lipid metabolism and modern-day human pathology (Herker et al., 2021; Kounakis et al., 2019; Pressly et al., 2022). Lipophagy, one of two routes for LD consumption, enables eukaryotic cells to use lipids as an energy source under nutrient deprivation, critically contributing to the energetic balance of a cell. This route requires docking of LDs to the lysosome/vacuole before uptake of LDs. However, the means of LD attachment was unknown in both yeast and human cells. Here, we have identified the molecular machinery that tethers LDs to the vacuole to enable LD consumption via lipophagy in yeast. The LD-localized LDO proteins attach to the vacuolar armadillo repeat protein Vac8 to form vCLIP, the vacuolar-LD contact site. We demonstrate that vCLIP emerges specifically in response to nutrient exhaustion and is critical for efficient LD consumption via lipophagy.

The LDO proteins Ldo45 and Ldo16, encoded by overlapping genomic regions, were initially identified as accessory factors of yeast seipin, contributing to spatial organization of LD biogenesis at the ER (Eisenberg-Bord et al., 2018; Teixeira et al., 2018). Seipin is a conserved ER membrane protein with many functions in LD biogenesis that is frequently mutated in lipodystrophy (Fei et al., 2008; Klug et al., 2021; Magré et al., 2001; Salo et al., 2019). Similar to the LDO proteins in yeast, also LDAF1/Promethin, the putative mammalian LDO homolog, supports LD biogenesis by interacting with seipin and subsequently dissociates from seipin to target the surface of mature LDs (Castro et al., 2019; Chung et al., 2019). We demonstrate that at a later step, when nutrients are exhausted, the LDO proteins establish vCLIP by redistributing from the entire LD surface to defined Vac8-positive foci of adjacent vacuoles. LD-vacuole tethering is a prerequisite for subsequent LD consumption via microlipophagy, as genetic ablation of either the LDO proteins or Vac8 prevented not only vCLIP formation but also vacuolar sequestration of LDs. We also find that Pdr16, a cytosolic protein that is recruited to LDO-positive LDs emerging from the ER (Eisenberg-Bord et al., 2018), redistributes from the LD surface to vCLIP in nutrient-exhausted cells. However, Pdr16 was not needed for either vCLIP formation or lipophagy, indicating that this protein does not have an essential structural function at these contact sites. Interestingly, vCLIP formation occurs independently of several cellular processes and molecular determinants previously linked to lipophagy, including the formation of vacuolar microdomains, the ESCRT machinery and core autophagy regulators (Garcia et al., 2021; Liao et al., 2021; Oku et al., 2017; Tsuji et al., 2017; van Zutphen et al., 2014; Wang et al., 2014). Despite efficiently forming vCLIP, several of the analysed mutants were devoid of LDs inside the vacuole, suggesting that vCLIP formation is a prerequisite but not sufficient for successful lipophagy.

Our data demonstrate that both LDO proteins are able to form vCLIP and induce lipophagy. Their differential expression related to the metabolic state of the cell likely reflects different roles in LD biology. We find that only Ldo16 is transcriptionally upregulated upon nutrient depletion, supporting the notion that Ldo16 tethers LDs to the vacuole specifically to facilitate LD consumption in starved cells. In contrast, Ldo45 was mainly expressed in nutrient-rich conditions, consistent with the previous suggestion that it functions in LD proliferation in association with seipin (Teixeira et al., 2018). Supporting this, a truncated Ldo45 variant lacking the whole of Ldo16 failed to form vCLIP but was still targeted to LDs. Our series of Ldo16 mutants revealed that its N-terminal hydrophobic domains insert it into the ER, its cationic amphipathic helix redirect it to the LD surface and its C-terminal intrinsically disordered peptide region associates with Vac8 at the vacuolar surface. Formation of vCLIP was prevented by deleting *VAC8*, and phenocopied by loss of Pfa3, which palmitoylates Vac8 to anchor it to the vacuolar membrane. This resulted in a redistribution of the LDO proteins over the entire LD surface and prevented lipophagy. Notably, re-targeting Vac8 to the nuclear envelope was sufficient to re-route LDs to these sites, demonstrating that Vac8 is a vCLIP tether.

Multiple aspects of vacuolar homeostasis require the armadillo repeat protein Vac8, including vacuolar inheritance/fusion and the coordination of various macro- and microautophagic processes (Boutouja et al., 2019; Tang et al., 2003; Wang et al., 2001). Vac8 interacts with Atg13 and Atg11 to recruit the phagophore assembly site (PAS) to the vacuole to support bulk and selective autophagy, respectively, and interacts with Nvj1 to form the NVJ and facilitate microautophagic degradation of portions of the nucleus via PMN (Gatica et al., 2021; Hollenstein et al., 2021, 2019; Roberts et al., 2003). Crystallographic studies have demonstrated that both Atg13 and Nvj1 interact with Vac8 via disordered loops that associate with the minor groove of the armadillo repeat superhelix. According to ColabFold, the disordered loop of Ldo16 is predicted to occupy the same groove, which is the most highly conserved interface of Vac8, despite being spread across multiple armadillo repeats. Thus, Atg13, Nvj1 and Ldo16 might compete for binding, and Vac8 may switch from one binding partner to another to fine-tune multiple autophagic processes. Related to this, recruitment of the selective PAS to the vacuolar membrane has been proposed to derive from avidity-mediated Vac8–Atg11 interactions with low affinity that are stabilized by a high local concentration and limited diffusion of the interaction partners (Erlendsson and Teilum, 2020; Hollenstein et al., 2021). The same concept of a body to be autophagocytosed acting as a platform to concentrate Vac8 has been applied to the NVJ (Park et al., 2020), and now can be seen in the context of LDs. Once an Ldo16-decorated LD comes into proximity with the rim of the NVJ, it can access a pool of highly concentrated Vac8, which exchanges its partner from Nvj1 to Ldo16 to form vCLIP. This concept places Vac8 as a key regulator of autophagic processes during starvation, with all of macroautophagy, PMN and lipophagy being routed through the same final common pathway. In support of this, we find that the loss of vCLIP formation upon genetic ablation of the LDO proteins results in NVJ expansion and induction of PMN. Though a functional link between PMN and lipophagy remains to be explored, our findings establish Vac8 as critical regulator of the contact sites that enable delivery of diverse cargo to the vacuole.

## Acknowledgements

This work was supported by the Swedish Research Council Vetenskapsrådet (2019-05249 to SB, 2019-04004 to JLH and 2019-04052 to CA), the Knut and Alice Wallenberg foundation (2017.009 to CA, JLH and SB), Stiftelsen Olle Engkvist Byggmästare (207-0527 to SB), Cancerfonden (20045 to CA, 22 2488Pj to SB and 211865Pj to JLH) and the Alfonso Martín Escudero foundation (Postdoctoral fellowship to AdO). We thank Christian Ungermann for providing the Vac8 antibody, Claudine Kraft for sharing plasmids encoding Vac8ι1N targeted to the vacuole and the nER and Katharina Keuenhof, Vajradhar Acharya and Jake Croft for help with immuno-EM data quantification strategy.

## Author contributions

Conceptualization, IAG and SB; Methodology, IAG, EB, FB, SG, AdO, LH, CA, TPL, JLH and SB; Investigation, IAG, EB, FB, SG, AdO, LH and TPL; Formal Analysis, IAG, FB and SB; Resources, CA, TPL, JLH, and SB; Writing – Original Draft: IAG and SB; Writing – Review & Editing, TPL and SB; Supervision: CA, TPL, JLH and SB; Funding Acquisition, CA, JLH and SB.

## Figures and Figure legends

## Material and Methods

### Yeast strains and genetics

All experiments were performed in BY4742 (*MATα; his3*Δ1; *leu2*Δ0; *lys2*Δ0; *ura3*Δ0) or BY4741 (*MATa his3*Δ1; *leu2*Δ0; *met15*Δ0; *ura3*Δ0). Yeast transformations were conducted as previously described, and deletion or endogenous tagging of genes via homologous recombination was performed following standard procedure (Janke et al., 2004). All yeast strains and plasmids used in this study are listed in Supplemental Table 1 and 2, respectively. The plasmids encoding Ldo45 or GFP-tagged Ldo46 (pRS313-spLdo45 and pRS313-spLdo45^GFP^, respectively) were created by PCR amplification of respective fragments from the genome, excluding the intron region and including the Ldo45 promotor, followed by fragment joining by overlapping PCR (Heckman and Pease, 2007). For pRS313-spLdo45^GFP^ a yeGFP from pYM25 was C-terminally added. The final constructs were introduced into the pRS313 backbone using *BamH*I and *Xho*I restriction sites. The pRS313-Ldo16 and pRS313-Ldo16^GFP^ was constructed in the same way, amplifying the genomic *YMR148w* region. Truncation mutants of Ldo16 and Ldo45 (Ldo16^ΔC24^, Ldo16^ΔC54^, Ldo16^ΔC94^, Ldo16^ΔN49^, Ldo16^ΔN72^, Ldo45^ΔC148^) were created by amplifying the corresponding fragments from the pRS313-spLdo45^GFP^ or the pRS313-Ldo16^GFP^ and fusing them by overlapping PCR, preserving the promotor, the yeGFP sequence and the plasmid backbone. To create the Ldo16 variants carrying point mutations in the putative cationic amphipathic helix (pRS313-Ldo16^5xA^ and pRS313-Ldo16^2xE^) the point mutations were introduced in respective oligonucleotides, and the amplified fragments containing the mutated sites were joined with overlapping PCR. The genomic single deletion of *LDO16* (Δ*ldo16*) was created using the *delitto perfetto* approach (Storici and Resnick, 2006). Briefly, the open reading frame coding for Ldo45 and Ldo16 was first substituted with a cassette containing *KlURA3* and *kanMX4*. Then, the sequence corresponding to spliced *LDO45* was amplified from pRS313-spLdo45 and recombined to replace the cassette. All oligonucleotides used in this study are listed in Supplemental Table 3.

### Yeast culturing conditions

All strains were grown in baffled Erlenmeyer flasks at 28°C and shaking at 145 rpm in synthetic complete medium (SC), containing 0.17% yeast nitrogen base (BD Difco™), 0.5% (NH_4_)_2_SO_4_ (Carl Roth) and 30 mg/l of all amino acids (except 80 mg/l histidine and 200 mg/l leucine and, for the BY4742 strain, 120 mg/l lysine), 30 mg/l adenine and 320 mg/l uracil, with 2% glucose (SCD). Plasmid-containing strains were grown in SCD without histidine, uracil or leucine. Overnight cultures were incubated for 16-20 h in SCD and used to inoculate cultures to OD_600_ 0.1, followed by culturing into glucose exhaustion and stationary phase. For nitrogen starvation conditions, cells were pre-inoculated at 0.01 OD_600_ and grown in SCD for 12 h, washed with water and diluted 1:10 into SD without (NH_4_)_2_SO_4_ and amino acids (SD-N). For phosphate restriction, cells were inoculated in SCD prepared using YNB without phosphate (FORMEDIUM). Phosphate was supplemented as NaH_2_PO_4_ to a final concentration of 0.2 mM (instead of 7 mM as in standard SCD), allowing regular growth but leading to early entry into stationary phase due to phosphate exhaustion as described before (Ebrahimi et al., 2021; Peselj et al., 2022). For oleate supplementation, cultures were inoculated to OD_600_ 0.1 in SCD and grown for 5 h before addition of 0.5% sodium oleate (final concentration; dissolved in 0.1% Tween-20) (Sigma-Aldrich O3880) or of the respective solvent control. For deletion and tagging of genes, yeast cells were grown in rich medium (YPD) containing 20 g/l peptone (Gibco™ Bacto™ BD Biosciences), 10 g/l yeast extract (Bacto™ BD Biosciences) and 4% glucose. For subsequent selection of mutants, YPD plates containing hygromycin B (FORMEDIUM, HYG5000), nourseothricin sulphate (Jena Biosciences, AB102XL) or G418 (Sigma-Aldrich, A1720-5G) or SCD plates with all amino acids except for histidine, uracil or leucine, were used.

### Analysis of cell growth

Cells from overnight cultures were used to inoculate 250 μl SCD to OD600 0.1 in 96-well microplates with clear, flat bottom (Greiner Bio-ONE). Plates were shaking at 999 rpm and 28°C and growth was measured by monitoring OD600 every 2 h for 10 h using a plate reader (2300 EnSpire™, Perkin Elmer).

### Immunoblot Analysis

6 OD600 of cells were harvested by centrifugation, lysed with lysis buffer (1.85 M NaOH; 7.5% β-mercaptoethanol) and incubated on ice for 10 min. After addition of 55% TCA, samples were again incubated on ice for 10 min, centrifugated for 10 min at 4°C, and the pellets were resuspended in urea loading buffer (200 mM Tris-HCl; 8 M urea; 5% SDS; 1 mM EDTA; 0.02% bromophenol blue; 15 mM DTT; pH 6.8). Samples were incubated for 10 min at 65°C, centrifuged and the supernatant was loaded on 12.5% polyacrylamide gels for separation via SDS-PAGE, followed by blotting on PVDF membranes (ROTH). Membranes were blocked in 5% milk/TBS (500 mM Tris; 1.5 M NaCl; pH 7.4) and were fixed by incubating them shaking in acetone for 30 min at 4°C. Then membrane were dried at 50°C for another 30 min, reactivated with 99% ethanol, washed and probed with antibodies against the GFP-epitope (dilution 1:2500, mouse, Roche 181446001), α-Tubulin (dilution 1:10000, rabbit, Abcam, 184970), GAPDH (dilution 1:10000, mouse, Thermo Fischer, MA5-15738) and Vac8 (dilution 1:10000, rabbit, gift from Christian Ungermann) as well as respective peroxidase-conjugated secondary antibodies against mouse (dilution 1:10,000, rabbit, Sigma A9044) or rabbit (dilution 1:10000, goat, Sigma A0545). Clarity Western ECL Substrate (BIO-RAD) and a ChemiDoc XRS + Imaging System (BIO-RAD) were used for detection. Densitometric quantification was performed with Image Lab 5.2.1 Software (BIO-RAD).

### Quantitative Real-Time PCR

To assess gene expression via qRT-PCR, approximately 30 OD600 of cells were collected and total RNA was purified using the Ribopure-Yeast™ kit (Thermo Fisher, AM1926). Genomic DNA was digested using TURBO™ DNase (Invitrogen™ AM2238) according to the supplier’s protocol. 2 μg of total RNA was reverse transcribed with SuperScript transcriptase (Thermo Fisher, 18064014) following manufacturer’s instructions. qRT-PCR was performed in triplicates with the KAPA SYBR® Fast qPCR Master mix (Sigma-Aldrich, KK4600) using a Rotor-Gene Q (Qiagen) PCR cycler. Data are presented as fold changes using the comparative Ct method (ΔΔCT) (Livak and Schmittgen, 2001) and *UBC6* as housekeeping gene. All primers used for qRT-PCR are listed in Supplemental Table 3.

### Confocal Fluorescence Microscopy

1 OD600 of cells were harvested at indicated time points, stained using indicated dyes as described below and seeded on 3% agar/PBS (137 mM NaCl, 2.7 mM KCl, 10 mM Na2HPO4, 1.8mM KH2PO4; pH 7.4). The samples were imaged using a ZEISS LSM780 microscope equipped with an 63x/1.40 Oil M27 objective using the ZEN black software. For Figure 1G, a ZEISS LSM800 Airyscan microscope equipped with an 63x/1.40 Oil M27 objective and the ZEN blue software control was used. Appropriate laser and detector settings were applied to visualize endogenously tagged proteins (GFP, mCherry or mScarlet) or dyes. For LD stainings, wells were incubated for 10 min in the dark with BODIPY 493/503 (0.12 nM; Fischer Scientific, 11540326) or monodansylpentane (MDH) (100 μM; AUTODOT™ Abcepta, #SM1000b) and then washed with PBS prior to seeding on agar-coated slides and microscopic analysis. For vacuolar staining, cells were either incubated for 30 min in the dark with CellTracker™ Blue CMAC Dye (1 mM; Thermo Fisher, C2111) and washed with PBS before imaging, or supplemented with FM 4-64FX (Invitrogen™, 11574816) to a final concentration of 15 μM directly in the culture media until collecting the cells for microscopic analysis.

### Image analysis and quantification

The open-source software Fiji (Schindelin et al., 2012) was used to process and quantify confocal micrographs. To process the images, Gaussian filtering (s = 0.8-1) was applied, followed by background subtraction (rolling ball radius = 50–100 pixels) and Unsharp mask settings when required. Images from the same experiment were processed with similar settings unless stated otherwise in the respective figure legend. Brightness and contrast were adjusted for each channel equally in individual experiments for analysis. The ratio of ‘LDs in/out’ of the vacuole using BODIPY was calculated by automatically measuring Integrated Density (IntDen) ‘inside of vacuoles’ segmented with the Huang algorithm and divided by the IntDen of ‘outside of vacuoles’ (calculated by subtracting the IntDen ‘inside the vacuoles’ from the IntDen of the whole cell, segmented with YeastMate) (Bunk et al., 2022). All intensity measurements were performed in the original, unprocessed images. LD size was quantified by segmenting LDs with the Yen algorithm and measurement of the area. Number of LDs per cell was quantified by automated counting of the segmented LDs against the mask of segmented cells with Find Maxima. In both cases, the results were analyzed with a frequency distribution and were fitted to a gaussian curve. The colocalization analysis to obtain the Pearson and Mander’s M2 coefficients was performed with the JaCoP plugin, thresholding images with only Gaussian blur, subtract background and Unsharp Mask adjusted. Quantification of cells according to LDO^GFP^ distribution (Fig. 5E) as well as according to NVJ formation using Nvj1^GFP^ (Fig. 5F) in the Vac8 truncation mutants was performed manually using the Cell counter plugin.

### Transmission electron microscopy

For transmission electron microscopy, samples were prepared as described (Tosal-Castano et al., 2021). Briefly, a Wohlwend Compact 03 (M. Wohlwend GmbH, Sennwald, Switzerland) was used for high-pressure freezing of samples, and freeze-substitution was performed for 1 h in a Leica EM AFS2 (Leica Microsystems, Vienna, Austria) using the Leica reagent bath with a flow-through ring of 2% uranyl acetate dissolved in 90% acetone and 10% methanol at -90°C (Hawes et al., 2007). Samples were washed twice in acetone while the temperature was gradually raised (2.9°C per h to -50°C). Samples were infiltrated with increasing amounts of Lowicryl HM20 (Polysciences, Warrington, PA, 15924-1) mixed with acetone (1:4, 2:3, 1:1, 4:1 and 100% 3x) at -50°C and a duration of 2 h per step. UV light was used to induce polymerization for 72 h at -50°C, followed by 24 h at room temperature. A Reichert-Jung Ultracut E Ultramicrotome (C. Reichert, Vienna, Austria) with an ultra 45° diamond knife (Diatome, Biel, Switzerland) was used to cut sections of 70 nm, which were collected on copper slot grids coated with 1% Formvar (TAAB). 2% uranyl acetate in dH2O and Reynold’s lead citrate were applied for on-section contrast staining (Reynolds, 1963). A Tecnai T12 electron microscope equipped with a Ceta CMOS 16M camera (FEI Co., Eindhoven, the Netherlands) was used for imaging of samples at 120 kV. Quantification of LDs in direct physical contact with the vacuole or the ER as well as PMN events per section was performed manually.

### Immuno-electron microscopy

For immunogold labelling, the sample blocks were prepared and embedded in HM20 resin as described above, as this sample preparation has been shown to be amenable to immune-labelling (Panagaki et al., 2021). Grids with 70 nm-thick sections were fixed in 1% paraformaldehyde in PBS for 10 min and blocked with 0.1% fish skin gelatin and 0.8% BSA in PBS for 1 h. For primary antibody labeling, samples were incubated overnight with an antibody against Vac8 (dilution 1:30, rabbit, gift from Christian Ungermann), followed by 3x wash in PBS for 20 min and 1 h incubation with a secondary gold-conjugated antibody (anti-rabbit IgG 10 nm gold; EMS Electron Microscopy Sciences). After washing with PBS, glutaraldehyde (2.5%) was applied to the sections for postfixing for 1 h, followed by contrast staining using uranyl acetate and Reynold’s lead citrate as described above. The detection of gold particles was done automatically using the IMODfindbeads program (Kremer et al., 1996) inside of an area including 30 nm (the size of the antibody sandwich plus the gold particle) on either side of the membranes. Special drawing tools were used to efficiently model such areas in the IMOD suite of programs, followed by automated extraction of the quantification of the gold beads per area.

### Structural modelling

To model the conservation of the Vac8 surface, a multiple sequence alignment was made that focussed on Vac8 only using 2 rounds of HHblits, which searches into a nr30 database (non-redundant above 30% identity). After the first iteration 106 hits were included with e-value <10^-20^. After the second iteration, 294 proteins were included (e-value < 10^-42^). This excluded other Armadillo repeat proteins. Aligned sequences were submitted along with the solved structure of Vac8 to the ConSurf server (Ashkenazy et al., 2010) to obtain conservation scores for residues on the surface scaled between 0-10. To model the interaction of Vac8 with Ldo16, ColabFold (Mirdita et al., 2022) was seeded with Vac8 residues (560 residues, 19-578, missing the disordered N-terminus) and full length Ldo16 (148 residues). The structure shown in Fig. 5H is the rank 1 model, for which a version was obtained with side-chains positioned in relaxed conformations.

### Statistical analysis

Data are presented either as dot plots, showing individual data points, mean (line) and error bars representing standard error of mean (s.e.m.), or as line graphs with symbols depicting mean and s.e.m. (protein or mRNA level) or as line graphs depicting a gaussian non-linear regression fit histograms (frequency distribution). Sample size, referring to independent biological replicates, is indicated in the respective figure legends. Statistical analysis was performed using GraphPad Prism (v8.0). Shapiro-Wilk’s test and visual inspection of Q-Q-plots was used to check for normal distribution of data, and analysis of variance (ANOVA) with *Tukey’s post hoc* test was used for comparisons between multiple groups. The equality of group variances was checked with the Brown-Forsythe-Test. Due to unequal variances of data shown in Fig. 6B, a *Welch’s* ANOVA with *Dunnett T3 post hoc* test was used. Where appropriate, a two-way ANOVA, corrected with *Tukey’s* or *Bonferroni’s multiple comparisons test*, was applied. Significances are presented as ***p < 0.001, **p < 0.01, and *p < 0.05. Details for all statistical analyses performed are listed in Supplemental Table 4.

## Supplemental Information titles

**Supplemental Figure S1** (related to Figure 1)

**Expression of Ldo16 is induced in response to nutrient depletion**

**Supplemental Figure S2** (related to Figure 2)

**The LDO proteins target LDs to the vacuole via the shared Ldo16 region**

**Supplemental Figure S3** (related to Figure 4)

**Microscopic analysis of vCLIP formation in mutants linked to lipophagy or general LD biology**

**Supplemental Figure S4** (related to Figure 5)

**C-terminal modification of Vac8 affects growth and NVJ formation**

**Supplemental Table 1: Yeast strains used in this study.**

**Supplemental Table 2: Plasmids used in this study.**

**Supplemental Table 3: Oligonucleotides used in this study.**

**Supplemental Table 3: Details of statistical analyses**

