## Supplemental Material for "LDO proteins and Vac8 form a vacuole-lipid droplet contact site required for lipophagy in response to starvation"

**Including Supplemental Figures S1-S4 and Supplemental Tables 1-4**

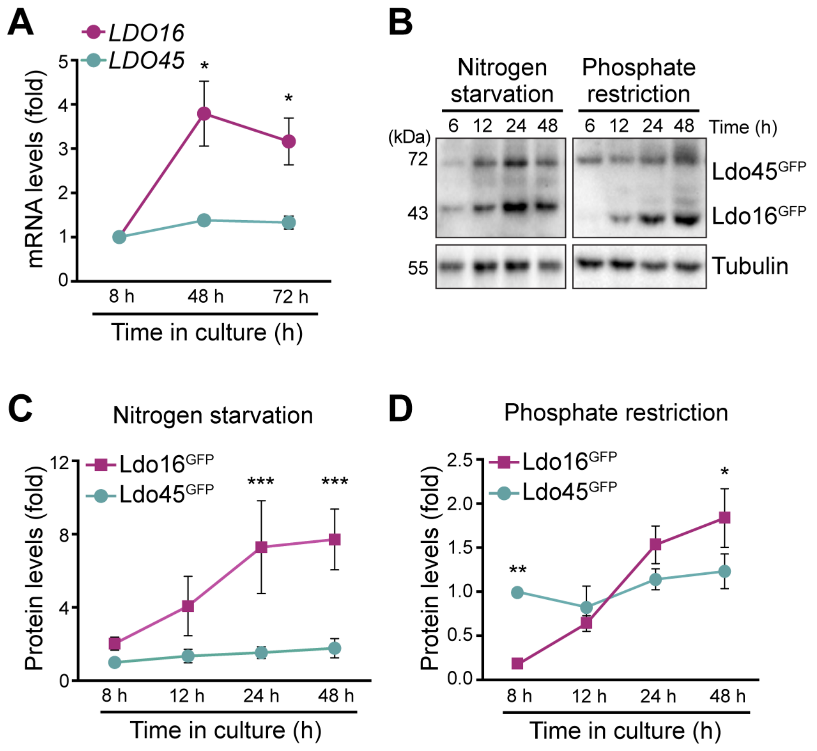


**Supplemental Figure S1** (related to Figure 1)

**Expression of Ldo16 is induced in response to nutrient depletion**

**A)** Quantification of *LDO16* and *LDO45* mRNA levels via RT-qPCR in wild type cells collected during exponential growth (8 h) in standard glucose media as well as upon growth into stationary phase upon glucose exhaustion (48 h and 72 h). Data are presented as fold changes using the comparative Ct method (ΔΔCT) and *UBC6* as housekeeping gene, depicted as fold of transcript levels at 8 h. Data represent mean ± s.e.m.; $n$ = 4. **B-D)** Immunoblot analysis of total protein extracts of cells endogenously expressing LDO$^{GFP}$ (allowing the simultaneous detection of both Ldo16$^{GFP}$ and Ldo45$^{GFP}$) collected after 8, 12, 24 and 48 h of culturing in nitrogen-depleted and phosphate-restricted media, respectively. Blots were probed with antibodies directed against GFP and tubulin as loading control. Representative blots (B) and corresponding densitometric quantification of Ldo45$^{GFP}$ and Ldo16$^{GFP}$ levels (C, D), normalized to tubulin and depicted as fold of Ldo45$^{GFP}$ at 8 h, are shown; Data represent mean ± s.e.m.; $n$ = 4 (for nitrogen starvation) and $n$ = 5 (for phosphate restriction); * $p < 0.05$, ** $p < 0.01$ and *** $p < 0.001$. See Supplemental Table 4 for details on statistical analyses.

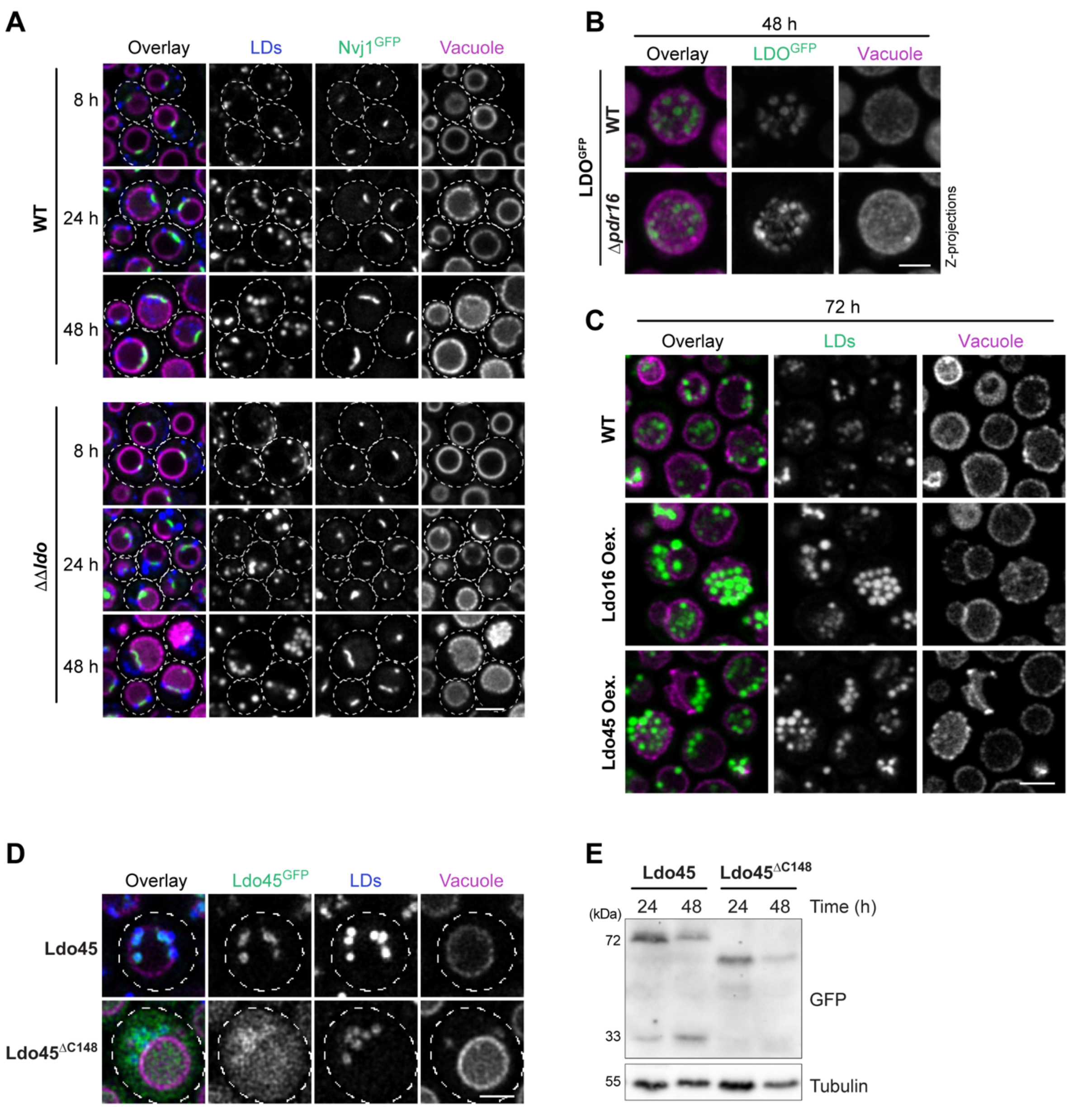


**Supplemental Figure S2** (related to Figure 2)

**The LDO proteins target LDs to the vacuole via the shared Ldo16 region**

**A)** Confocal micrographs of wild type (WT) and ΔΔ*ldo* cells endogenously expressing Nvj1$^{GFP}$ and Vph1$^{mCherry}$ in glucose-rich conditions (8h) and after growth into glucose exhaustion (24 h and 48 h). LDs were stained with MDH. Scale bar: 3 μm. A part of the microscopic analysis at 48 h is also shown in Figure 2B. **B)** Z-projections of confocal micrographs of glucose-exhausted (48 h) WT and Δ*pdr16* cells endogenously expressing either LDO$^{GFP}$ and additionally carrying with Vph1$^{mCherry}$. Scale bar: 2 μm. **C)** Extended data corresponding to the confocal microscopy shown in Figure 2K, now also including the single channel micrographs. WT cells as well as ΔΔ*ldo* cells ectopically overexpressing either Ldo45 or Ldo16, in addition equipped with Vph1$^{mCherry}$ and stained with BODIPY 493/503, were analyzed upon prolonged glucose exhaustion (72 h). Scale bar: 3 μm. **D)** Confocal micrographs of ΔΔ*ldo* cells ectopically expressing GFP-tagged Ldo45 or the C-terminally truncated variant Ldo45$^{\Delta C148}$, which lacks the complete Ldo16 region. The vacuole was visualized via Vph1$^{mCherry}$, the LDs were stained with MDH and cells were analyzed upon glucose exhaustion (48 h). Scale bar: 2 μm. **E)** Immunoblot analysis of total protein extracts of cells described in (D), collected after 24 h and 48 h of culturing. Blots were probed with antibodies directed against GFP to detect the Ldo45 variants and tubulin as loading control.

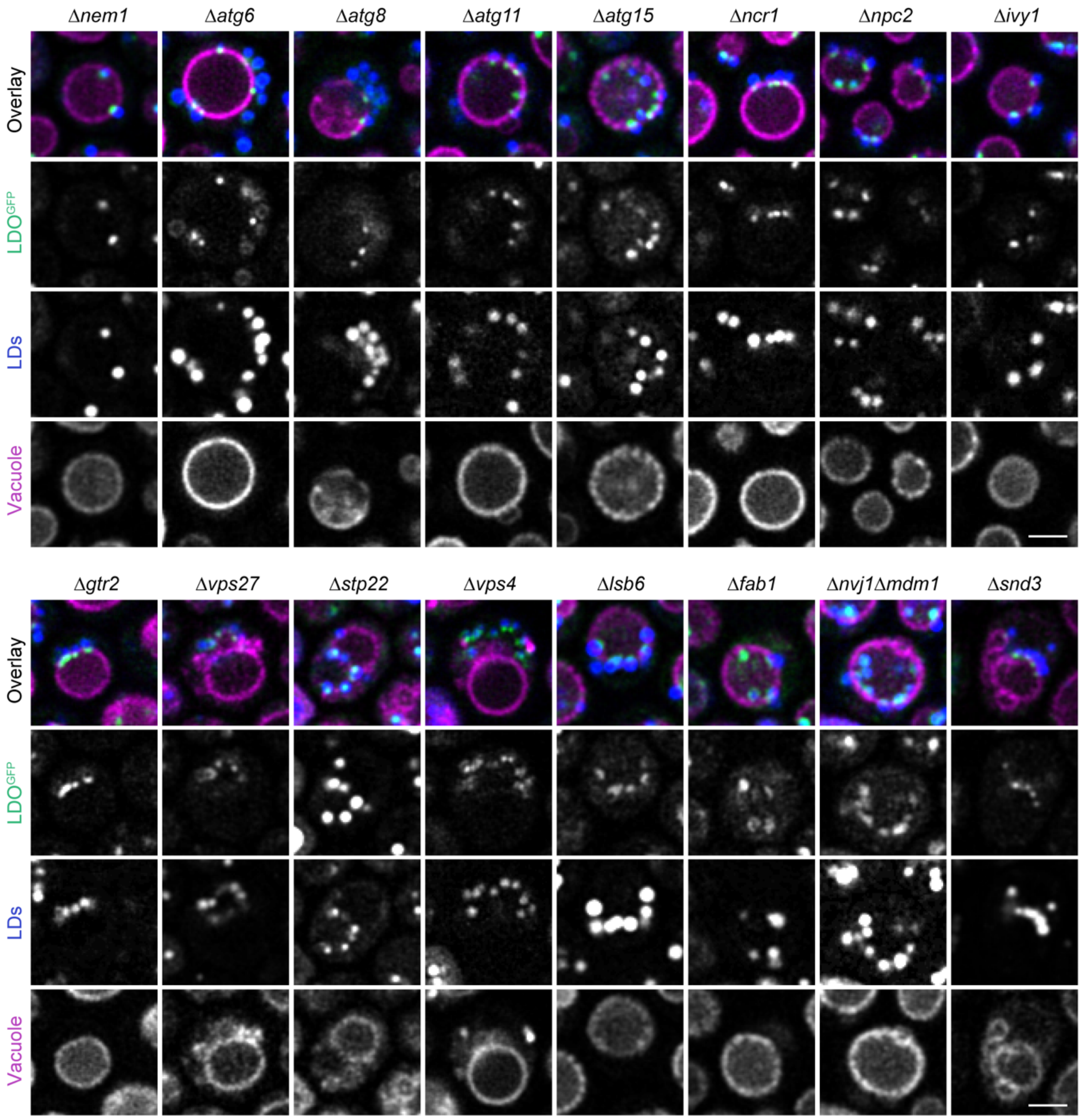


**Supplemental Figure S3** (related to Figure 4)

**Microscopic analysis of vCLIP formation in mutants linked to lipophagy or general LD biology**

Confocal micrographs of $LDO^{GFP}$ localization in indicated deletion mutants, all previously associated with LD biosynthesis, macro- and microautophagy, the ESCRT machinery, NVJ formation and vacuolar membrane lipid composition. All mutants expressed $LDO^{GFP}$ and $Vph1^{mCherry}$ and were stained with the LD stain MDH upon growth into glucose exhaustion (48 h). Selected mutants are shown in Figure 4. Scale bar: 2 μm.

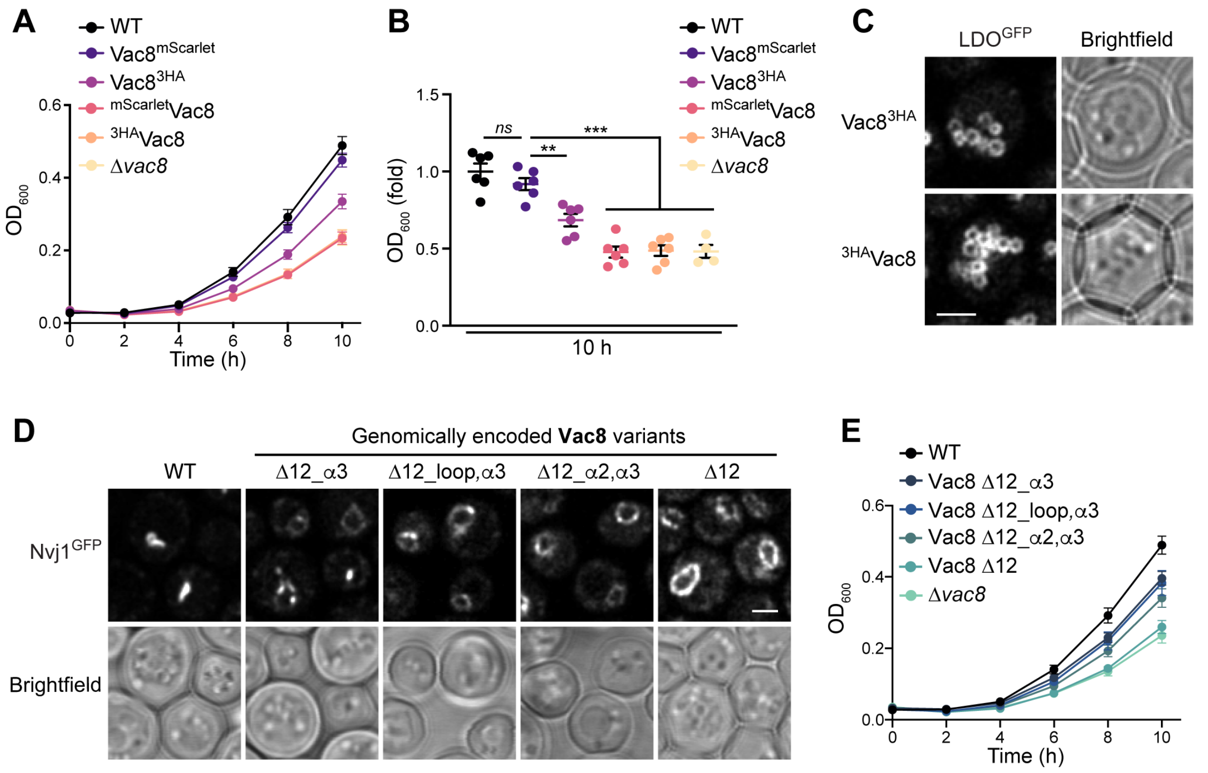


**Supplemental Figure S4** (related to Figure 5)

**C-terminal modification of Vac8 affects growth and NVJ formation**

**A, B)** Analysis of cellular growth of wild type (WT) and Δ*vac8* cells as well as of cells endogenously expressing C- and N-terminal fusions of Vac8 with mScarlet or 3HA as indicated. Optical density was determined every 2 h and data is shown as mean ± s.e.m. (A). To facilitate comparison, the 10 h time point is additionally depicted in (B), showing all data points as well as mean (line) and s.e.m.; $n = 4$ (for Δ*vac8*) and $n = 6$ (for all others). **C)** Confocal micrographs of cells expressing $LDO^{GFP}$ and Vac8, C-terminally or N-terminally fused to 3HA, after 48 h of culturing. Scale bar: 2 μm. **D)** Confocal micrographs of WT cells and cells endogenously expressing indicated C-terminally truncated Vac8 variants, additionally equipped with $Nvj1^{GFP}$ to visualize NVJ formation upon growth into glucose exhaustion (48 h). Corresponding quantification of cells according to NVJ morphology is shown in Figure 5F. **E)** Analysis of cellular growth of wild type (WT) and Δ*vac8* cells as well as of cells endogenously expressing indicated C-terminally truncated Vac8 variants. Optical density was determined every 2 h for 10 h and data is shown as mean ± s.e.m.; $n = 6$; ** $p < 0.01$ and *** $p < 0.001$. See Supplemental Table 4 for details on statistical analyses.

**Supplemental Table 1: Yeast strains used in this study.**

| Strain | Genotype | Source |
|---|---|---|
| BY4742 | MATα *his3Δ1 leu2Δ0 lys2Δ0 ura3Δ0* | Euroscarf |
| $Erg1^{GFP}$ | BY4741*, ERG1*-GFP[S65T]::HIS3 | Euroscarf |
| $Faa4^{GFP}$ | BY4742, *FAA4*-GFP::natNT2 | This study |
| $Pdr16^{GFP}$ | BY4742, *PDR16*-GFP::hphNT1 | This study |
| $LDO^{GFP}$ | BY4742, *YMR147/148W*-GFP::hphNT1 | This study |
| $Faa4^{GFP}$ $Vph1^{mCherry}$ | BY4742, *FAA4*-GFP::natNT2, *VPH1*-mCherry::kanMX | This study |
| $Pdr16^{GFP}$ $Vph1^{mCherry}$ | BY4742, *PDR16*-GFP::hphNT1, *VPH1*-mCherry::kanMX | This study |
| $LDO^{GFP}$ $Vph1^{mCherry}$ | BY4742, *YMR147/148W*-GFP::hphNT1, *VPH1*-mCherry::kanMX | This study |
| ΔΔ*ldo* $Pdr16^{GFP}$ $Vph1^{mCherry}$ | BY4742, *ymr148w*Δ::hphNT1, *PDR16*-GFP::hphNT1, *VPH1*-mCherry::kanMX | This study |
| Δ*pdr16* $LDO^{GFP}$ $Vph1^{mCherry}$ | BY4742, *pdr16*Δ::hphNT1, *YMR147/148W*-GFP::natNT2, *VPH1*-mCherry::kanMX | This study |
| $LDO^{GFP}$ $Faa4^{mCherry}$ | BY4742, *YMR147/148W*-GFP::hphNT1, *FAA4*-mCherry::kanMX | This study |
| $Faa4^{GFP}$ $LDO^{mCherry}$ | BY4742, *YMR147/148W*-GFP::hphNT1, *FAA4*-mCherry::kanMX | This study |
| $Nvj1^{GFP}$ $Vph1^{mCherry}$ | BY4742, *NVJ1*-GFP::natNT2, *VPH1*-mCherry::kanMX | This study |
| ΔΔ*ldo* $Nvj1^{GFP}$ $Vph1^{mCherry}$ | BY4742, *ymr148w*Δ::hphNT1, *NVJ1*-GFP::natNT2, *VPH1*-mCherry::kanMX | This study |
| ΔΔ*ldo* | BY4742, *ymr148w*Δ::hphNT1 | This study |
| $Vph1^{mCherry}$ | BY4742, *VPH1*-mCherry::kanMX | This study |
| ΔΔ*ldo* $Vph1^{mCherry}$ | BY4742, *ymr148w*Δ::hphNT1, *VPH1*-mCherry::kanMX | This study |
| Δ*ldo45* $Vph1^{mCherry}$ | BY4742, *ymr147w*Δ*0-448*::hphNT1, *VPH1*-mCherry::kanMX | This study |
| Δ*ldo16* $Vph1^{mCherry}$ | BY4742, *ymr147w*Δ *ymr148*Δ::*spLDO45*, *VPH1*-mCherry::kanMX | This study |
| Δ*ldo45* $Ldo16^{GFP}$ | BY4742, *ymr147w*Δ*0-448*::hphNT1, *LDO16*-GFP:: natNT2 | This study |
| Δ*ldo16* $Ldo45^{GFP}$ | BY4742, *ymr147w*Δ ymr148wΔ::*spLDO45*-GFP::hphNT1 | This study |
| $LDO^{GFP}$ $Vph1^{mCherry}$ | BY4742, *YMR147/148W*-GFP::hphNT1, *VPH1*-mCherry::kanMX | This study |
| Δ*ldo45* $Ldo16^{GFP}$ $Vph1^{mCherry}$ | BY4742, *ymr147w*Δ*0-448*::hphNT1, *YMR148W*-GFP::natNT2, *VPH1*-mCherry::kanMX | This study |
| Δ*ldo16* $Ldo45^{GFP}$ $Vph1^{mCherry}$ | BY4742, *ymr147w*Δ*ymr148w*Δ::*spLDO45*-GFP::hphNT1, *VPH1*-mCherry::kanMX | This study |
| ΔΔ*ldo* $Vph1^{mCherry}$ p-Empty | BY4742, *ymr148w*Δ::hphNT1, *VPH1*-mCherry::kanMX, pRS313-*HIS3* | This study |
| ΔΔ*ldo* $Vph1^{mCherry}$ p-Ldo16 | BY4742, *ymr148w*Δ::hphNT1, *VPH1*-mCherry::kanMX, pRS313-*HIS3* Ldo16 | This study |
| ΔΔ*ldo* $Vph1^{mCherry}$ p-Ldo45 | BY4742, *ymr148w*Δ::hphNT1, *VPH1*-mCherry::kanMX, pRS313-*HIS3* spLdo45 | This study |
| ΔΔ*ldo* $Vph1^{mCherry}$ p-$Ldo45^{GFP}$ | BY4742, *ymr148w*Δ::hphNT1, *VPH1*-mCherry::kanMX, pRS313-*HIS3* spLdo45-GFP | This study |
| ΔΔ*ldo* $Vph1^{mCherry}$ p-$Ldo45^{\Delta C148\text{-}GFP}$ | BY4742, *ymr148w*Δ::hphNT1, *VPH1*-mCherry::kanMX, pRS313-*HIS3* spLdo45Δ*C148*-GFP | This study |
| ΔΔ*ldo* $Vph1^{mCherry}$ p-$Ldo16^{GFP}$ | BY4742, *ymr148w*Δ::hphNT1, *VPH1*-mCherry::kanMX, pRS313-*HIS3* Ldo16-GFP | This study |
| ΔΔ*ldo* $Vph1^{mCherry}$ p-$Ldo16^{\Delta C24\text{-}GFP}$ | BY4742, *ymr148w*Δ::hphNT1, *VPH1*-mCherry::kanMX, pRS313-*HIS3* Ldo16Δ124-148-GFP | This study |
| ΔΔ*ldo* $Vph1^{mCherry}$ p-$Ldo16^{\Delta C54\text{-}GFP}$ | BY4742, *ymr148w*Δ::hphNT1, *VPH1*-mCherry::kanMX, pRS313-*HIS3* Ldo16Δ94-148-GFP | This study |

| | | |
|---|---|---|
| ΔΔ*ldo* Vph1$^{mCherry}$ p-Ldo16$^{\Delta C98-GFP}$ | BY4742, *ymr148w*Δ::hphNT1, *VPH1*-mCherry::kanMX, pRS313-*HIS3* Ldo16Δ*50-148*-GFP | This study |
| ΔΔ*ldo* Vph1$^{mCherry}$ p-Ldo16$^{\Delta N49-GFP}$ | BY4742, *ymr148w*Δ::hphNT1, *VPH1*-mCherry::kanMX, pRS313-*HIS3* Ldo16Δ1-49-GFP | This study |
| ΔΔ*ldo* Vph1$^{mCherry}$ p-Ldo16$^{\Delta N72-GFP}$ | BY4742, *ymr148w*Δ::hphNT1, *ymr148w*Δ::hphNT1, *VPH1*-mCherry::kanMX, pRS313-*HIS3* Ldo16Δ1-73-GFP | This study |
| ΔΔ*ldo* Vph1$^{mCherry}$ p-Ldo16$^{5xA-GFP}$ | BY4742, *ymr148w*Δ::hphNT1, *VPH1*-mCherry::kanMX, pRS313-*HIS3* Ldo16-F65A, L66A, V69A, L70A, M73A-GFP | This study |
| ΔΔ*ldo* Vph1$^{mCherry}$ p-Ldo16$^{2xE-GFP}$ | BY4742, *ymr148w*Δ::hphNT1, *VPH1*-mCherry::kanMX, pRS313-*HIS3* Ldo16-L66E, V69E-GFP | This study |
| LDO$^{GFP}$ Vph1$^{mCherry}$ Δ*nvj1* | BY4742, *YMR147/148W*-GFP::hphNT1, *VPH1*-mCherry::kanMX, *nvj*Δ*1*::URA3 | This study |
| LDO$^{GFP}$ *Vph1*$^{mCherry}$ Δ*mdm1* | BY4742, *YMR147/148W*-GFP::hphNT1, *VPH1*-mCherry::kanMX, *mdm*Δ*1*::URA3 | This study |
| LDO$^{GFP}$ Vph1$^{mCherry}$ Δ*atg1* | BY4742, *YMR147/148W*-GFP::hphNT1, *VPH1*-mCherry::kanMX, *atg*Δ*1*::URA3 | This study |
| LDO$^{GFP}$ Vph1$^{mCherry}$ Δ*atg6* | BY4742, *YMR147/148W* -GFP::hphNT1, *VPH1*-mCherry::kanMX, *atg6*Δ::natNT2 | This study |
| LDO$^{GFP}$ Vph1$^{mCherry}$ Δ*vps4* | BY4742, *YMR147/148W*-GFP::hphNT1, *VPH1*-mCherry::kanMX, *vps4*Δ::natNT2 | This study |
| LDO$^{GFP}$ Vph1$^{mCherry}$ Δ*ivy1* | BY4742, *YMR147/148W*-GFP::hphNT1, *VPH1*-mCherry::kanMX, *ivy1*Δ::natNT2 | This study |
| LDO$^{GFP}$ Vph1$^{mCherry}$ Δ*gtr2* | BY4742, *YMR147/148W*-GFP::hphNT1, *VPH1*-mCherry::kanMX, *gtr2*Δ::natNT2 | This study |
| LDO$^{GFP}$ Vph1$^{mCherry}$ Δ*vac8* | BY4742, *YMR147/148W*-GFP::hphNT1, *VPH1*-mCherry::kanMX, *vac8*Δ::natNT2 | This study |
| LDO$^{GFP}$ Vph1$^{mCherry}$ Δ*atg11* | BY4742, *YMR147/148W*-GFP::hphNT1, *VPH1*-mCherry::kanMX, *atg11*Δ::URA | This study |
| LDO$^{GFP}$ Vph1$^{mCherry}$ Δ*atg8* | BY4742, *YMR147/148W*-GFP::hphNT1, *VPH1*-mCherry::kanMX, *atg8*Δ::URA | This study |
| LDO$^{GFP}$ Vph1$^{mCherry}$ Δ*npc2* | BY4742, *YMR147/148W*-GFP::hphNT1, *VPH1*-mCherry::kanMX, *npc2*Δ::URA | This study |
| LDO$^{GFP}$ Vph1$^{mCherry}$ Δ*ncr1* | BY4742, *YMR147/148W*-GFP::hphNT1, *VPH1*-mCherry::kanMX, *ncr1*Δ::URA | This study |
| LDO$^{GFP}$ Vph1$^{mCherry}$ Δ*npc2* Δ*ncr1* | BY4742, *YMR147/148W*-GFP::hphNT1, *VPH1*-mCherry::kanMX, *npc2*Δ::URA, *ncr1*Δ::LEU | This study |
| LDO$^{GFP}$ Vph1$^{mCherry}$ Δ*snd3* | BY4742, *YMR147/148W*-GFP::hphNT1, *VPH1*-mCherry::kanMX, *snd3*Δ::URA | This study |
| LDO$^{GFP}$ Vph1$^{mCherry}$ Δ*sei1* | BY4742, *YMR147/148W* GFP::hphNT1, *VPH1*-mCherry::kanMX, *sei1*Δ::URA | This study |
| LDO$^{GFP}$ Vph1$^{mCherry}$ Δ*sac1* | BY4742, *YMR147/148W* GFP::hphNT1, *VPH1*-mCherry::kanMX, *sac1*Δ::natNT2 | This study |
| LDO$^{GFP}$ Vph1$^{mCherry}$ Δ*stp22* | BY4742, *YMR147/148W* GFP::hphNT1, *VPH1*-mCherry::kanMX, *stp22*Δ::natNT2 | This study |
| LDO$^{GFP}$ Vph1$^{mCherry}$ Δ*snf7* | BY4742, *YMR147/148W*-GFP::hphNT1, *VPH1*-mCherry::kanMX, *snf7*Δ::natNT2 | This study |
| LDO$^{GFP}$ Vph1$^{mCherry}$ Δ*lsb6* | BY4742, *YMR147/148W*-GFP::hphNT1, *VPH1*-mCherry::kanMX, *lsb6*Δ::natNT2 | This study |
| LDO$^{GFP}$ Vph1$^{mCherry}$ Δ*vac17* | BY4742, *YMR147/148W*-GFP::hphNT1, *VPH1*-mCherry::kanMX, *vac17*Δ::natNT2 | This study |
| LDO$^{GFP}$ Vph1$^{mCherry}$ Δ*vps27* | BY4742, *YMR147/148W*-GFP::hphNT1, *VPH1*-mCherry::kanMX, *vps27*Δ::natNT2 | This study |
| LDO$^{GFP}$ Vph1$^{mCherry}$ Δ*nvj1* Δ*mdm1* | BY4742, *YMR147/148W*-GFP::hphNT1, *VPH1*-mCherry::kanMX, *nvj1*Δ::HIS, *mdm1*Δ::natNT2 | This study |
| LDO$^{GFP}$ Vph1$^{mCherry}$ Δ*pfa3* | BY4742, *YMR147/148W*-GFP::hphNT1, *VPH1*-mCherry::kanMX, *pfa3*Δ::natNT2 | This study |
| LDO$^{GFP}$ Vph1$^{mCherry}$ Δ*atg15* | BY4742, *YMR147/148W*-GFP::hphNT1, *VPH1*-mCherry::kanMX, *atg15*Δ::natNT2 | This study |
| LDO$^{GFP}$ Vph1$^{mCherry}$ Δ*nem1* | BY4742, *YMR147/148W*-GFP::hphNT1, *VPH1*-mCherry::kanMX, *nem1*Δ::natNT2 | This study |

| | | |
|---|---|---|
| LDO$^{GFP}$ Vph1$^{mCherry}$ Δ*fab1* | BY4742, *YMR147/148W*-GFP::hphNT1, *VPH1*-mCherry::kanMX, *fab1*Δ::URA | This study |
| LDO$^{GFP}$ Vac8$^{mScarlet}$ | BY4742, *YMR147/148W*-GFP::hphNT1, *VAC8*-mScarlet::URA | This study |
| LDO$^{GFP}$ $^{mScarlet}$Vac8 | BY4742, *YMR147/148W* GFP::hphNT1, natNT2::CUP1-mScarlet-*VAC8* | This study |
| LDO$^{GFP}$ Vac8$^{3HA}$ | BY4742, *YMR147/148W*-GFP::hphNT1, *VAC8*-3HA::natNT2 | This study |
| LDO$^{GFP}$ $^{3HA}$Vac8 | BY4742, *YMR147/148W*-GFP::hphNT1, natNT2::ADH1-3HA-*VAC8* | This study |
| Vac8$^{mScarlet}$ | BY4742, *VAC8*-mScarlet::URA | This study |
| ΔΔ*ldo* Vac8$^{mScarlet}$ | BY4742, *ymr148w*Δ::hphNT1, *VAC8*-mScarlet::URA | This study |
| LDO$^{GFP}$ Vac8 Δ12 | BY4742, *YMR147/148W*-GFP::hphNT1, *VAC8*[Δaa495-578]::URA3 | This study |
| LDO$^{GFP}$ Vac8 Δ12_α2,α3 | BY4742, *YMR147/148W*-GFP::hphNT1, *VAC8*[Δaa507-578]::URA3 | This study |
| LDO$^{GFP}$ Vac8 Δ12_loop,α3 | BY4742, *YMR147/148W*-GFP::hphNT1, *VAC8*[Δaa533-578]::URA3 | This study |
| LDO$^{GFP}$ Vac8 Δ12_α3 | BY4742, *YMR147/148W*-GFP::hphNT1, *VAC8*[Δaa559-578::URA3 | This study |
| Nvj1$^{GFP}$ Vac8 Δ12 | BY4742, *NVJ1*-GFP::natNT2, *VAC8*[Δaa495-578]::URA3 | This study |
| Nvj1$^{GFP}$ Vac8 Δ12_α2,α3 | BY4742, *NVJ1*-GFP::natNT2, *VAC8*[Δaa507-578]::URA3 | This study |
| Nvj1$^{GFP}$ Vac8 Δ12_loop,α3 | BY4742, *NVJ1*-GFP::natNT2, *VAC8*[Δaa533-578]::URA3 | This study |
| Nvj1$^{GFP}$ Vac8 Δ12_α3 | BY4742, *NVJ1*-GFP::natNT2, *VAC8*[Δaa559-578]::URA3 | This study |
| LDO$^{GFP}$ Δ*vac8* Vph1-mSc | BY4742, *YMR147/48W*-GFP::hphNT1, pRS416-*URA3* Vph1-mScarlet, *vac8*Δ::HIS3 | This study |
| LDO$^{GFP}$ Δ*vac8* Vph1-mSc-Vac8ΔN | BY4742, *YMR147/48W*-GFP::hphNT1, pRS416-*URA3* Vph1-mScarlet -Vac8[19-578], *vac8*Δ::HIS3 | This study |
| LDO$^{GFP}$ Δ*vac8* Nvj1$^{1-125}$-mSc | BY4742, *YMR147/48W*-GFP::hphNT1, pRS416-*URA3* Nvj1[1-125]-mScarlet, *vac8*Δ::HIS3 | This study |
| LDO$^{GFP}$ Δ*vac8* Nvj1$^{1-125}$-mSc-Vac8ΔN | BY4742, *YMR147/48W*-GFP::hphNT1, pRS416-*URA3* Nvj1[1-125]-mScarlet-VAC8[19-578], *vac8*Δ::HIS3 | This study |
| LDO$^{GFP}$ Δ*vac8* pRS416-empty | BY4742, *YMR147/48W*-GFP::hphNT1, pRS416-*URA3*,*vac8*Δ::HIS3 | This study |
| Faa4$^{GFP}$ | BY4742, *FAA4*-GFP::natNT2 | This study |
| ΔΔ*ldo* Faa4$^{GFP}$ | BY4742, *ymr148w*Δ::hphNT1, *FAA4*-GFP::natNT2 | This study |
| Δ*vac8* Faa4$^{GFP}$ | BY4742, *vac8*Δ::hphNT1, *FAA4*-GFP::natNT2 | This study |
| Vph1$^{mCherry}$ Δ*nvj1* | BY4742, *VPH1*-mCherry::kanMX, *nvj1*Δ::natNT2 | This study |
| Vph1$^{mCherry}$ Δ*nvj1* Δ*mdm1* | BY4742, *VPH1*-mCherry::kanMX, *mdm1*Δ::natNT2, *nvj1*Δ::HIS3 | This study |
| Vph1$^{mCherry}$ Δ*vac8* | BY4742, *VPH1*-mCherry::kanMX, *vac8*Δ::natNT2 | This study |
| Vph1$^{mCherry}$ ΔΔ*ldo* Δ*nvj1* Δ*mdm1* | BY4742, *VPH1*-mCherry::kanMX, *ymr148w*Δ::hphNT1, *mdm1*Δ::natNT2, *nvj1*Δ::HIS3 | This study |

**Supplemental Table 2: Plasmids used in this study.**

| Name | Plasmid | Origin |
|---|---|---|
| Vector control | pRS313-*HIS3* | (Sikorski et al., 1989) |
| p-Ldo16 | pRS313-*HIS3* Ldo16 | This study |
| p-Ldo45 | pRS313-*HIS3* spLdo45 | This study |
| p-Ldo45$^{GFP}$ | pRS313-*HIS3* spLdo45-GFP | This study |
| p-Ldo45$^{\Delta C148\text{-}GFP}$ | pRS313-*HIS3* spLdo45ΔC148-GFP | This study |
| p-Ldo16$^{GFP}$ | pRS313-*HIS3* Ldo16-GFP | This study |
| p-Ldo16$^{\Delta C24\text{-}GFP}$ | pRS313-*HIS3* LDO16Δ124-148-GFP | This study |
| p-Ldo16$^{\Delta C54\text{-}GFP}$ | pRS313-*HIS3* Ldo16Δ94-148-GFP | This study |
| p-Ldo16$^{\Delta C98\text{-}GFP}$ | pRS313-*HIS3* Ldo16Δ50-148-GFP | This study |
| p-Ldo16$^{\Delta N49\text{-}GFP}$ | pRS313-*HIS3* Ldo16Δ1-49-GFP | This study |
| p-Ldo16$^{\Delta N72\text{-}GFP}$ | pRS313-*HIS3* Ldo16Δ1-73-GFP | This study |
| p-Ldo16$^{5xA\text{-}GFP}$ | pRS313-*HIS3* Ldo16-F65A, L66A, V69A, L70A, M73A-GFP | This study |
| p-Ldo16$^{2xE\text{-}GFP}$ | pRS313-*HIS3* Ldo16-L66E, V69E-GFP | This study |
| Vector control | pRS416-*URA3* | (Sikorski et al., 1989) |
| Vph1-mSc | pDH33 (pRS416-*URA3* Vph1-mScarlet) | (Hollenstein et al. 2019) |
| Vph1-mSc-Vac8ΔN | pDH39 (pRS416-*URA3* Vph1-mScarlet-Vac8[19-578]) | (Hollenstein et al. 2019) |
| Nvj1$^{1\text{-}125}$-mSc | pDH77 (pRS416-*URA3* Nvj1[1-125]-mScarlet) | (Hollenstein et al. 2021) |
| Nvj1$^{1\text{-}125}$-mSc-Vac8ΔN | pDH83 (pRS416-*URA3* Nvj1[1-125]-mScarlet-Vac8[19-578]) | (Hollenstein et al. 2021) |

**Supplemental Table 3: Oligonucleotides used in this study.**

| Modification | Oligonucleotides | PCR template |
|---|---|---|
| **Deletion and tagging of genes** | | |
| C-terminal tagging of *FAA4* | 5‘-CGTAGTGTTTATGAAGGGCAGGGGGGAAAGTAAA AAACTATGTCTTCCTTTAATCGATGAATTCGAGCTCG-3‘<br>5‘-TATTCTAGCGGCTGTCAAGCCAGATGTGGAAAG AGTTTAT AAAGAAAACACTCGTACGCTGCAGGTCGAC-3‘ | pSB56-GFP-natNT2 (This study)<br>pSB49-mCherry-kanMX (This study) |
| Control PCR *FAA4* tagging | 5‘-CGAGCTCGAATTCATCGAT-3‘<br>5‘-CGATGTTTCT TCGATAAAAGG-3‘ | |
| C-terminal tagging of *PDR16* | 5’-ATTATATATTATAGTGCATTATCATTATCTATCTAAATTTGCCT TTAATCGATGAATTCGAGCTCG-3’<br>5’-GAGGTACTCATGAAAAACTTCTTTACCCAGTAAAAT CGGAAA GCAGTACCGTGCGTACGCTGCAGGTCGAC-3’ | pSB56-GFP-natNT2 (This study) |
| Control PCR *PDR16* tagging | 5’-GACCTTGGGGTCGTGTTCGC-3’<br>5’-CGAGCTCGAATTCATCGAT-3’ | |
| C-terminal tagging of *YMR148w* | 5’-TTGACCTGCTAAACTTGCGAAAAATGTTTTTTTATTGC CGAGGTTAATCGATGAATTCGAGCTCG-3’<br>5’-CATTAGAGACTACTGCTAATAAAGCGGGTAATAAGTTCCA GCTCTCTCGTACGCTGCAGGTCGAC-3’ | pYM25-GFP (Janke et al., 2004)<br>pSB56-GFP-natNT2 (This study)<br>pSB49-mCherry-kanMX (This study) |
| Control PCR *YMR148w* tagging | 5’-CAATGGCGAAATTTACTTGAATCAC -3’<br>5’-CGAGCTCGAATTCATCGAT-3’ | |
| C-terminal tagging of *VPH1* | 5’-GAAGTACTTAAATGTTTCGCTTTTTTTAAAAGTCCTCAAAAT TTAATCGATGAATTCGAGCTCG-3’<br>5’-GACATGGAAGTCGCTGTTGCTAGTGCAAGCTCTT CCGCT TCAAGCCGTACGCTGCAGGTCGAC-3’ | pSB49-mCherry-kanMX (This study) |
| Control PCR *VPH1* tagging | 5’-CGAGCTCGAATTCATCGAT-3’<br>5’-GTATTCGAGGCCAATACTTG-3’ | |
| Deletion of *YMR148W* | 5‘-TCTCCTTTGCCATTGGACTTGTTATTCCGTGTTCCC TACTTTT TTTGATAATGCGTACGCTGCAGGTCGAC-3‘<br>5‘-TTGACCTGCTAAACTTGCGAAAAATGTTTTTTTATTGC CGAG GTTAATCGATGAATTCGAGCTCG-3‘ | pFA6a-hphNT1 (Janke et al., 2004)<br>pFA6a-natNT2 (Janke et al., 2004) |
| Control PCR *YMR148W* deletion | 5’-CGAGCTCGAATTCATCGAT-3’<br>5‘-CAATGGCGAAATTTACTTGAATCAC -3‘ | |
| Deletion of *PDR16* | 5‘-AACAAATAATACAAAACCACTTTATATAAAAAAA ATTACAAAA GCAAAAAAATGCGTACGCTGCAGGT CGAC-3‘<br>5‘-ATTATATATTATAGTGCATTATCATTATCTAT CTAAATTTGCCTTTAATCGATGAATTCGAGCTCG-3‘ | pFA6a-hphNT1 (Janke et al., 2004) |
| Control PCR *PDR16* deletion | 5’-CGAGCTCGAATTCATCGAT-3’<br>5‘-GACCTTGGGGTCGTGTTCGC-3‘ | |
| C-terminal tagging of *NVJ1* | 5’-GTTGTAAGTG ACGATGATAA CCGAGATGAC GGAAATATAG TACATTA ATCGATGAATTCGAGCTCG-3’<br>5’-CTAGATGCACAAGTGAACACTGAACAAGCATA CTCTCAACCA TTTAGATACCGTACGCTGCAGGTCGAC-3’ | pSB56-GFP-natNT2 (This study) |

| | | |
|---|---|---|
| Control PCR *NVJ1* tagging | 5'-CTATTGACCACATAATCCTTAG-3'<br>5'-CGAGCTCGAATTCATCGAT-3' | |
| Deletion of *YMR147WΔ1-448* | 5'-CGATTAAAATAAAAAGTGACATCTGAAAAACATCCA ATACTCC GATGCGTACGCTGCAGGTCGAC-3'<br>5'-AGTTTAAATAAATAAAAATAATCCAAAATACTT CGTACAAGCTATTTTGTAGAGCTCGATGAATTCGAGCTCG-3' | pFA6a-hphNT1 (Janke et al., 2004) |
| Control PCR *YMR147WΔ1-448* deletion | 5'-GTCGACCTGCAGCGTACG-3'<br>5'-CCTGGCCAGTTATCTAACG-3' | |
| *Delito perfetto* of *YMR147W* | 5'-CTTTCAAGACGATTAAAATAAAAAGTGACATCTGAAA AACATC CAATACTCCGGAGCTCGTTTTCGACACTGG-3'<br>5'-GATATTATTATTATTGACCTGCTAAACTTGCGAAAAATGTT TTTTTATTGCCGAGGTCCTTACCATTAAGTTGATC-3' | pCORE-*kl*URA3-kanaMX4 (Storici and Resnick, 2006) |
| Control PCR of *YMR147W delito perfetto* | 5'-GTCGACCTGCAGCGTACG-3'<br>5'-CCTGGCCAGTTATCTAACG-3' | |
| Deletion of *NVJ1* | 5'-TGTGCATAATATCAAAAAAGCTACAAATATAATTGTAA AATATAATAAGC ATG CGTACGCTGCAGGTCGAC-3'<br>5'-GTTGTAAGTGACGATGATAACCGAGATGACGGAAA TATAGTACATTAATCGATGAATTCGAGCTCG-3' | pFA6a-natNT2 (Janke et al., 2004) |
| Control PCR *NVJ1* deletion | 5'-TTGATAAGGCCTATTGTCGG-3'<br>5'-GTCGACCTGCAGCGTACG-3' | |
| Deletion of *MDM1* | 5'-GAAAGCGCCATAAGTGCGCGTGTTTGTGCCTTCTG ATATGA TATCGTATGCGTACGCTGCAGGTCGAC-3'<br>5'-CAATTACACTTTTTTTTTTAGATTGTTCGGTACTT AGTCAAGTT TTATTTTCAATCGATGAATTCGAGCTCG-3' | pFA6a-natNT2 (Janke et al., 2004) |
| Control PCR *MDM1* deletion | 5'-CGTCAAGGGTATCAGCAGAG-3'<br>5'-GTCGACCTGCAGCGTACG-3' | |
| Deletion of *ATG1* | 5'-ACCCCATATTTTCAAATCTCTTTTACAACA CCAGACGAGA AATTAAGAAA ATG CGTACGCTGCAGGTCGAC-3'<br>5'-ATATAGCAGGTCATTTGTACTTAATAAGAAAACC ATATT ATGCATCACTTAATCGATGAATTCGAGCTCG-3' | pSB36 pFA6a-URA3 (This study) |
| Control PCR *ATG1* deletion | 5'- GTAATGTAAGGAAAACCCAC-3'<br>5'- GTCGACCTGCAGCGTACG-3' | |
| Deletion of *ATG6* | 5'-GTTTTATGGCAGTCACTGTTTTCGCAAAGACTC CCAGAC ACGGGCATTAAAATGCGTACGCTGCAGGTCGAC-3'<br>5'-GAAATTTTCCCTTTATCACATTTATGAAAAAATG CATTTATAT GAACTACTTAATCGATGAATTCGAGCTCG-3' | pFA6a-natNT2 (Janke et al., 2004) |
| Control PCR *ATG6* deletion | 5'-CTCTTCGTTTTCACTCCAGTAC-3'<br>5'-CGAGCTCGAATTCATCGAT -3' | |
| Deletion of *VPS4* | 5'-ATGGAAGACAAAAATAAAGCAGCATAGAGTGCCTA TAGTAGATGGGGTACAAATGCGTACGCTGCAGGTCGAC-3'<br>5'-TTTTTTATTTTCATGTACACAAGAAATCTACATTAGCACGT TAATCAATTGACTAATCGATGAATTCGAGCTCG-3' | pFA6a-natNT2 Janke et al., 2004) |
| Control PCR *VPS4* deletion | 5'- CGAGCTCGAATTCATCGAT -3'<br>5'-CAGTCGCGCCAACGACCAGT-3' | |

| | | |
|---|---|---|
| Deletion of *IVY1* | 5‘-CCAAGGAATAAAATATTGTTAAAGAAAGTAACAGGA AGAGAAATCGGATATGCGTACGCTGCAGGTCGAC-3‘<br>5‘-GTTCACTTTCTCCATTTCTATATAAAAAGCATACA TAGAGTTAC AAATTTTAATCGATGAATTCGAGCTCG-3‘ | pFA6a-natNT2 (Janke et al., 2004) |
| Control PCR *IVY1* deletion | 5’-GACTTCAATCCCGTGCTTC-3’<br>5’-CGAGCTCGAATTCATCGAT -3’ | |
| Deletion of *GTR2* | 5‘-CACAGATTAACAAAACTCCAGGACAACGGTACTA ATACACA TACAACATGCGTACGCTGCAGGTCGAC-3‘<br>5‘-CATTTTCAATATGTATCTATATACCCTAATATTTTC ATGCC TTACGTCTTTCAATCGATGAATTCGAGCTCG-3‘ | pFA6a-natNT2 (Janke et al., 2004) |
| Control PCR *GTR2* deletion | 5’-CGAGCTCGAATTCATCGAT -3’<br>5’-GGAGTGGCTTTCCTCTTTTTTG-3’ | |
| Deletion of *VAC8* | 5‘-CTGAGCAAACTATAAGGGTGTTCTTTCTTCTGTA CTATATATACATTTGCAACTATGCGTACGCTGCAGGTCGAC-3‘<br>5‘-GAAAATTTTGATAAAAATTATAATGCCTAGTCC CGCTTTTGA AGAAAATCAATCGATGAATTCGAGCTCG-3 | pFA6a-natNT2 (Janke et al., 2004) |
| Control PCR *VAC8* deletion | 5’- CGAGCTCGAATTCATCGAT-3’<br>5’- GTCGACCTGCAGCGTACG-3’ | |
| Deletion of *ATG11* | 5‘-GTGTACTGTT GTTGTTCGGA AAGTACTTCT TTTATTTTCT TTTATACATC ATG CGTACGCTGCAGGTCGAC-3‘<br>5‘-GATACATAATTAAAATCTTGTCATTTGTGACAAACGT TTAGCACTGTTCA ATCGATGAATTCGAGCTCG-3‘ | pSB36 pFA6a-URA3 (This study) |
| Control PCR *ATG11* deletion | 5‘-GCTAGCATTC CTATATATCC-3‘<br>5‘-CGAGCTCGAATTCATCGAT-3‘ | |
| Deletion of *ATG8* | 5’CTAATAATTGTAAAGTTGAGAAAATCATAATAAAATAATTACTA GAGACATGCGTACGCTGCAGGTCGAC--3’<br>5’- CTATAATTTCGATTTTAGATGTTAACGCTTCATTTCT TTTC ATATAAAAGACTAATCGATGAATTCGAGCTCG-3’ | pSB36 pFA6a-URA3 (This study) |
| Control PCR *ATG8* deletion | 5’-CTAACTGTCTCCACCGATAATG-3’<br>5‘-CGAGCTCGAATTCATCGAT-3‘ | |
| Deletion of *NPC2* | 5’-CATAACCATATTAAATCTTCTCCTTCAAAGCTAG CACGCCTTC CAAATGCGTACGCTGCAGGTCGAC -3’<br>5’- GAACGAGAAGGGAAATAAACACGGATCAAT GAGTTGTATGAATCAGATCAATCGATGAATTCGAGC TCG-3’ | pSB36 pFA6a-URA3 (This study) |
| Control PCR *NPC2* deletion | 5’-CGCTTCTTGCCTAATGATAGG -3’<br>5’-CGAGCTCGAATTCATCGAT-3’ | |
| Deletion of *NCR1* | 5’-CTAAATTCATCTCCAAAAAGAACAAGAGCAGA ACTTCAATTAGTAAAACCATGCGTACGCTGCAGGTC GAC-3’<br>5’-GAATTTTTACCTATTTTTTCACTACGTAAAAT ATAGTATAA TCTGCTATGGCTAATCGATGAATTCGAGCTCG-3’ | pSB35 pFA6a-LEU2 (This study) |
| Control PCR *NCR1* deletion | 5’- CGAGCTCGAATTCATCGAT -3’<br>5’-CCTCTCTGAACAGGCCTG-3’ | |
| Deletion of *SND3* | 5‘-GAAACTATAACAGTATAACACAGCACAAGAGAACC GAGCAGCCCGCCATGCGTACGCTGCAGGTCGAC-3‘<br>5‘-ATACTTCGCTTTTGATCGAATCATTCAGCCTTAACA CCAGCGTTACCGGCATCGATGAATTCGAGCTCG-3‘ | pSB36 pFA6a-URA3 (This study) |

| | | |
|---|---|---|
| Control PCR *SND3* deletion | 5'- CGAGCTCGAATTCATCGAT -3'<br>5'- GCATAAAATCATATAGAC -3' | |
| Deletion of *SEI1* | 5'- CAAAATGTGAATCCAAGGTTTCAAGAAAATAA GATAAAG TGAATAGGAAGGATGCGTACGCTGCAGGTCGAC -3'<br>5'- CTTAGAAAATAACAGCTAGGTTTTAAAATTATATAGCGAG AAGTACAATTCTATCAATCGATGAATTCGAGCTCG -3' | pSB36 pFA6a-URA3 (This study) |
| Control PCR *SEI1* deletion | 5'- CGAGCTCGAATTCATCGAT -3'<br>5'- GCCACGTTATTTACACCTAC -3' | |
| Deletion of *SAC1* | 5'-CTATAACAGTAACGATAATATTTATATACACGTATA TTTTCTCGTCTAGATATGCGTACGCTGCAGGTCGAC-3'<br>5'-CGTTTTGGATTTACAATAATCATCATTTTATCAC ATATAGAAC TCATTAATCGATGAATTCGAGCTCG-3' | pFA6a-natNT2 (Janke et al., 2004) |
| Control PCR *SAC1* deletion | 5'- GTCGACCTGCAGCGTACG -3'<br>5'- CTGTTCCATCCCTCGAAACACTAGC-3' | |
| Deletion of *STP22* | 5'- CAGTTGGTATCTTAACGGCCAAGAAAAGAGA GAGAGTGAAGAGCAACGATGCGTACGCTGCAGGT CGAC -3'<br>5'- GTTAAAAAAATATTTTTTATGGCACTTCGGCGATG CGAAAG AAAGTGAGTCAATCGATGAATTCGAGCTCG -3' | pFA6a-natNT2 (Janke et al., 2004) |
| Control PCR *STP22* deletion | 5'- GCGTCTCGTTTCAAAGACCGTG -3'<br>5'- GTCGACCTGCAGCGTACG -3' | |
| Deletion of *SNF7* | 5'-CTCGGACGGAAGCAGCAGAAACATAACAGTATT GATAAATAAGGCATGCGTACGCTGCAGGTCGAC-3'<br>5'-GTAAGAACACCTTTTTTTTTCTTTCATCTAAACCG CATAGAACACGTTCAATCGATGAATTCGAGCTCG-3' | pFA6a-natNT2 (Janke et al., 2004) |
| Control PCR *SNF7* deletion | 5'-GCGGCTTCGTCGGAAATCAG-3'<br>5'- GTCGACCTGCAGCGTACG -3' | |
| Deletion of *LSB6* | 5'- CCGGGCATAAAGTGAACTAGACACTTTCAAGAAG CCAACC AAAGCATGCGTACGCTGCAGGTCGAC -3'<br>5'- GTTATGATTTCTTTATATTGAGTATGTATTGAATTATTTTCC<br>AAAAAATCAATCGATGAATTCGAGCTCG -3' | pFA6a-natNT2 (Janke et al., 2004) |
| Control PCR *LSB6* deletion | 5'- GTCGACCTGCAGCGTACG -3'<br>5'- CCACAGTTGTAGTCACGTGCG -3' | |
| Deletion of *VAC17* | 5'- GATAGATAAGAAACAGCTCGCATAAGGAAACAA GGACAC ATCGATTAATGCGTACGCTGCAGGTCGAC -3'<br>5'- GAATAAACATTTGGAGCAAAAGAAGAGTAG GTTAGGTAAAGGAGGCATTAATTAATCGATGAATTCGAGCTCG -3' | pFA6a-natNT2 (Janke et al., 2004) |
| Control PCR *VAC17* deletion | 5'- GTCGACCTGCAGCGTACG -3'<br>5'- GCATTACCGCCAGAACTAGCG -3' | |
| Deletion of *VPS27* | 5'-GATTTTTTTTTGCTAAGGTGAATGAGTAGTGAGTAAAG AACTAAGAACAGTATGCGTACGCTGCAGGTCGAC-3'<br>5'-GCGCTAGGTTTCTTTTTACAAATACATAGAAAAGGCTACAAT ATTAATCGATGAATTCGAGCTCG-3' | pFA6a-natNT2 (Janke et al., 2004) |

| | | |
|---|---|---|
| Control PCR *VPS27* deletion | 5’-CGGAGCCTACCTTTTAGCTTTTGC-3’<br>5’-GTCGACCTGCAGCGTACG-3’ | |
| Deletion of *PFA3* | 5’-CAAGACAAAGAAGCCGATCTTGGAATTCGATAGAATC TTTTTCGCAAATGCGTACGCTGCAGGTCGAC-3’<br>5’-GGATAATTTTCGTATTATCGTAACATTCATTCGTTCT AGTTT TATGACTAATCGATGAATTCGAGCTCG-3’ | pFA6a-natNT2 (Janke et al., 2004) |
| Control PCR *PFA3* deletion | 5’-GTCGACCTGCAGCGTACG-3’<br>5’-GCTCCTGATCATGTGTCTGTTGACG-3’ | |
| Deletion of *ATG15* | 5’- AACTGATCTAGGCATTACAATTAAAGGAAACAAGG GAAATA TTCTATTGAATGCGTACGCTGCAGGTCGAC -3’<br>5’- CGCATAGGCCCTAAAACAACACTAGGGTCAT AATAGAT GTATGGGTCTTAATCGATGAATTC GAGCTCG -3’ | pFA6a-natNT2 (Janke et al., 2004) |
| Control PCR *ATG15* deletion | 5’- TATAATAAGCATAACATCGG -3’<br>5’- GTCGACCTGCAGCGTACG -3’ | |
| Deletion of *NEM1* | 5’- CTCACTTTTCTTTATTGTTGTTAATTTCTGGAC ATTGTT TCATTAATTGAATGCGTACGCTGCAGGTCGAC-3’<br>5’- CGATTTATCACCTAAAATTCTCATCTTTTTTGGAAGGA CACAATAACAATTGTCAATCGATGAATTCGAGCTCG-3’ | pFA6a-natNT2 (Janke et al., 2004) |
| Control PCR *NEM1* deletion | 5’- CGAGCTCGAATTCATCGAT -3’<br>5’- CAAGTTGGGTTGAAAACACC-3’ | |
| Deletion of *FAB1* | 5’- CAAGGTAGCTTCCATCTTGCACACGCAAGACCGT CACACAG CATGCGTACGCTGCAGGTCGAC -3’<br>5’- ATAAAAAA AAGTTACAGA ATATAACTTG TACACGTTTA TGTATTA ATCGATGAATTCGAGCTCG -3’ | pSB36 pFA6a-URA3 (This study) |
| Control PCR *FAB1* deletion | 5’- CGAGCTCGAATTCATCGAT -3’<br>5’- CATTATGTTT GCAACAAAAT -3’ | |
| C-terminal tagging of *VAC8* | 5’- GAAAATTTTGATAAAAATTATAATGCCTAGTCCCGCTTTT GAAGAAAATCAATCGATGAATTCGAGCTCG -3’<br>5’- GCAAGTTTGGAATTGTATAATATTACTCAACAGA TTTTACA ATTTTTACATCGTACGCTGCAGGTCGAC -3’ | pSB22 pYM-C-mScarlet-URA3 (This study)<br>pYM17 3HA-natNT2 (Janke et al., 2004) |
| Control PCR *VAC8* C-terminal tagging | 5’- CGAGCTCGAATTCATCGAT -3’<br>5’- GTCGACCTGCAGCGTACG -3’ | |
| N-terminal tagging of *VAC8* | 5’- CTGAGCAAACTATAAGGGTGTTCTTTCTTCTGTA CTATATATACATTTGCAACTATGCGTACGCTGCAGGTCGAC -3’<br>5’- GTGAGACACTGGCCTCGTCTGAAGAATCTTTCAAGCAAC TACAACATGAACCCATCGATGAATTCTCTGTCG -3’ | pSB23 pYM-N-CUP1-mScarlet-natNT2 (This study)<br>pYM N8-ADH1-3HA-natNT2 (Janke et al., 2004) |
| Control PCR *VAC8* N-terminal tagging | 5’- GTCGACCTGCAGCGTACG-3’<br>5’- CGTATAAGCATACGGTAAGGAAGTGGG-3’ | |
| Truncation of Vac8$^{\Delta 495\text{-}578}$ | 5’-CTACCTTTGAACACATTGCGCTATGGACAATTTTA CAAT TGCTAGAAAGTTAACGTACGCTGCAGGTCGAC -3’<br>5’- GAAAATTTTGATAAAAATTATAATGCCTAGTCC CGCTTTTGA AGAAAATCAATCGATGAATTCGAGCTCG -3’ | pSB36 pFA6a-URA3 (This study) |
| Truncation of Vac8$^{\Delta 507\text{-}578}$ | 5’- CAATTGCTAGAAAGTCATAATGATAAAGTGGAAGAT TTGG TTAAAAATGATTAACGTACGCTGCAGGTCGAC-3’ | pSB36 pFA6a-URA3 (This study) |

| | | |
|---|---|---|
| | 5’- GAAAATTTTGATAAAAATTATAATGCCTAGTCC CGCTTTT GAAGAAAATCAATCGATGAATTCGAGCTCG -3’ | |
| Truncation of Vac8$^{\Delta 533\text{-}578}$ | 5’- GAAAAATGGCAGATGTGACCTTTGAGCGTTTACA AAGATCAGGAATTGATGTTAAATAACGTACGCTGCAGGTCGAC-3’<br>5’- GAAAATTTTGATAAAAATTATAATGCCTAGTCC CGCTTTTG AAGAAAATCAATCGATGAATTCGAG CTCG -3’ | pSB36 pFA6a-URA3 (This study) |
| Truncation of Vac8$^{\Delta 559\text{-}578}$ | 5’- GAATGATAACAACAGTAATAACAATGACACAGGATCTGAA CATCAACCTGTATAACGTACGCTGCAGGTCGAC-3’<br>5’- GAAAATTTTGATAAAAATTATAATGCCTAGTCCCGCTTTTG AAGAAAATCAATCGATGAATTCGAGCTCG -3’ | pSB36 pFA6a-URA3 (This study) |
| Control PCR of Vac8$^{\Delta 12}$ truncations | 5’- CGAGCTCGAATTCATCGAT -3’<br>5’- GTCGACCTGCAGCGTACG -3’ | |
| **RT-qPCR** | | |
| *LDO16* | 5‘-CCACAGGAGCCACTTTCCAC -3‘<br>5‘-TGGCGCTTTGAGCACTAACA -3‘ | |
| *LDO45* | 5‘-TTCCCCAATGTGGGTGACTA -3‘<br>5‘-CACGAAGTGCACGGAAATGA ACGTCT-3‘ | |
| *UBC6* | 5'-GGACGTTTCAAGCCCAACAC-3'<br>5'-TGAGACAGACCAGCCAGGAT-3' | |
| **Generation of LDO mutants** | | |
| p-Ldo16 | 5’-CTCCACCGCGGTGGCGGCCGCTCTAGAACTAGTGGATCCCTGGTT CCCCAATGTGGG-3’<br>5’-GGGAACAAAAGCTGGGTACCGGGCCCCCCCTCGAGGATCGGTAAAG CTAAGATATATGCG-3’ | genomic DNA pRS313 (Sikorski et al., 1989) |
| p-Ldo45 | 5’-CTCCACCGCGGTGGCGGCCGCTCTAGAACTAGTGGATCCCCGTTCAA CGAATACCAAAC-3’<br>5’-GCTATCTTGTTGTCCAAAAGTATTGCTCACATAGTCACCCACATTGG-3’<br>5’-CTACTGGTTCCCCAATGTGGGTGACTATGTGAGCAAT ACTTTTGGACAACAAG-3’<br>5’-GGGAACAAAAGCTGGGTACCGGGCCCCCCCTCGAGGA TCGGTAAAGCTAAGATATATGCG-3’ | genomic DNA pRS313 (Sikorski et al., 1989) |
| p-Ldo16$^{GFP}$ | 5’-ATCTGGATCCCTGGTTCCCCAATGTGGG-3’<br>5’-TCCAGCACCAGCACCAGCACCTGCTCCGGAAGAGAG CTGGAACTTATTAC-3’<br>5’-AATAAAGCGGGTAATAAGTTCCAGCTCTCTTCCGGAG CAGGTGCTGGT-3’<br>5’-AATAAAGCGGGTAATAAGTTCCAGCTCTCTTCCGGA GCAGGTGCTGGT-3’ | p-Ldo16 (This study) pRS313 (Sikorski et al., 1989) pYM25-yeGFP (Janke et al., 2004) |
| p-Ldo45$^{GFP}$ | 5’-ATCGGGATCCCGTTCAACGAATACCAAAC-3’<br>5’-TAATTCTTCACCTTTAGACAGAATTGCTCCAGAGA GCTGGAACTTATTAC-3’<br>5’-GCTGGAGCAATTCTGTCTAATAAAGCGGGTAATA AGTTCCAGCTCTCT-3’<br>5’-CGATCTCGAGCAGTTATTTGTACAATTCATCC-3’ | p-Ldo45 (This study) pRS313 (Sikorski et al., 1989) pYM25-yeGFP (Janke et al., 2004) |
| p-Ldo45$^{\Delta C148\text{-}GFP}$ | 5’-ATCGGGATCCCGTTCAACGAATACCAAAC-3’<br>5’-GTTATTCCGTGTTCCCTACTTTTTTTGATATCCGGAG CAGGTGCTG-3’<br>5’-TCCAGCACCAGCACCAGCACCTGCTCCGGATATCA | p-Ldo45 (This study) pRS313 (Sikorski et al., 1989) pYM25-yeGFP (Janke et al., 2004) |

| | | |
|---|---|---|
| | AAAAAAGTAGGGAACACGG-3’<br>5’-ATTATATCAAAAAAAGTAGGGAACACGG-3’<br>5’-ATCTCTCGAGTTATTTGTACAATTCATCCATAC-3’ | |
| p-Ldo16$^{\Delta C24\text{-}GFP}$ | 5’-ATCTGGATCCCTGGTTCCCCAATGTGGG-3’<br>5’-CGTCACCGAAGAAGATGTTATTTTCGAACCTGTTTCCGG AGCAGGTGCTGGTGC-3’<br>5’-TCCAGCACCAGCACCAGCACCTGCTCCGGAAACAGG TTCGAAAATAACATC-3’<br>5’-ATCTCTCGAGTTATTTGTACAATTCATCCATAC-3’ | p-Ldo16 (This study)<br>pRS313 (Sikorski et al., 1989)<br>pYM25-yeGFP (Janke et al., 2004) |
| p-Ldo16$^{\Delta C54\text{-}GFP}$ | 5’-ATCTGGATCCCTGGTTCCCCAATGTGGG-3’<br>5’-TCCAGCACCAGCACCAGCACCTGCTCCGGATGGCCT TAATGTGGAAAGTG-3’<br>5’-TCCAGCACCAGCACCAGCACCTGCTCCGGATGGCC TTAATGTGGAAAGTG-3’<br>5’-ATCTCTCGAGTTATTTGTACAATTCATCCATAC-3’ | p-Ldo16 (This study)<br>pRS313 (Sikorski et al., 1989)<br>pYM25-yeGFP (Janke et al., 2004) |
| p-Ldo16$^{\Delta C98\text{-}GFP}$ | 5’-ATCTGGATCCCTGGTTCCCCAATGTGGG-3’<br>5’-TCCAGCACCAGCACCAGCACCTGCTCCGGAGTTGCTG CAGAAGCCAAATG-3’<br>5’-GGCGTGGTATCATTTGGCTTCTGCAGCAACTCCGG AGCAGGTGCTGGT-3’<br>5’-ATCTCTCGAGTTATTTGTACAATTCATCCATAC-3’ | p-Ldo16 (This study)<br>pRS313 (Sikorski et al., 1989)<br>pYM25-yeGFP (Janke et al., 2004) |
| p-Ldo16$^{\Delta N49\text{-}GFP}$ | 5’-ATCTGGATCCCTGGTTCCCCAATGTGGG-3’<br>5’-ATAAATGAGTTGGGCCATTTTGAAACTCATTATCAAAA AAAGTAGGGAACAC-3’<br>5’-GTTATTCCGTGTTCCCTACTTTTTTTGATAATGAGTTTCAAA ATGGCCCA-3’<br>5’-TCCAGCACCAGCACCAGCACCTGCTCCGGAAGAGAG CTGGAACTTATTAC-3’<br>5’-AATAAAGCGGGTAATAAGTTCCAGCTCTCTTCCGGAG CAGGTGCTGGTGC-3’<br>5’-ATCTCTCGAGTTATTTGTACAATTCATCCATAC-3’ | p-Ldo16 (This study)<br>pRS313 (Sikorski et al., 1989)<br>pYM25-yeGFP (Janke et al., 2004) |
| p-Ldo16$^{\Delta N72\text{-}GFP}$ | 5’-ATCTGGATCCCTGGTTCCCCAATGTGGG-3’<br>5’-TAGTTGTGCCGGCTGTGTTTGTAGGGCCATTATCAAAA AAAGTAGGGAACAC-3’<br>5’-GTTATTCCGTGTTCCCTACTTTTTTTGATAATGGCCCTAC AAACACAGC-3’<br>5’-ATCTCTCGAGTTATTTGTACAATTCATCCATAC-3’ | p-Ldo16 (This study)<br>pRS313 (Sikorski et al., 1989)<br>pYM25-yeGFP (Janke et al., 2004) |
| p-Ldo16$^{5xA\text{-}GFP}$ | ATCTGGATCCCTGGTTCCCCAATGTGGG-3’<br>5’-TGCTTTGTCAGCAGCTTTTTTCGCGGCGGCGTCAGCTCGG ACATA-3’<br>5’-GCCGCGAAAAAAGCTGCTGACAAAGCAGCCCTACAAACA CAGCCG-3’<br>5’-ATCTCTCGAGTTATTTGTACAATTCATCCATAC-3’ | p-Ldo16 (This study)<br>pRS313 (Sikorski et al., 1989)<br>pYM25-yeGFP (Janke et al., 2004) |
| p-Ldo16$^{2xE\text{-}GFP}$ | 5’-ATCTGGATCCCTGGTTCCCCAATGTGGG-3’<br>5’-CCGAGCTGACGCCTTCGAGAAAAAAGAGCTCGACAAAA TGGCCC-3’<br>5’-GGGCCATTTTGTCGAGCTCTTTTTTCTCGAAGGCGTCA GCTCGG-3’<br>5’-ATCTCTCGAGTTATTTGTACAATTCATCCATAC-3’ | p-Ldo16 (This study)<br>pRS313 (Sikorski et al., 1989)<br>pYM25-yeGFP (Janke et al., 2004) |

### Supplemental Table 4: Details of statistical analyses.

| **Figure 1F** | | | |
|---|---|---|---|
| **Two-way ANOVA** | **P value** | **P value summary** | **F (DFn, DFd)** |
| **Row Factor** | <0.0001 | **** | F (3, 21) = 15.17 |
| **Column Factor** | 0.1244 | ns | F (1, 7) = 3.048 |
| **Row Factor x Column Factor** | <0.0001 | **** | F (3, 21) = 56.16 |

| **Tukey's multiple comparisons test** | **Mean Diff.** | **95.00% CI of diff.** | **Summary** | **Adjusted P Value** |
|---|---|---|---|---|
| 6:Ldo45$^{GFP}$ vs. 6:Ldo16$^{GFP}$ | 0.3871 | 0.1345 to 0.6397 | *** | 0.0009 |
| 6:Ldo45$^{GFP}$ vs. 12:Ldo45$^{GFP}$ | -0.1198 | -0.3724 to 0.1328 | ns | 0.7505 |
| 6:Ldo45$^{GFP}$ vs. 12:Ldo16$^{GFP}$ | -0.2359 | -0.4884 to 0.01670 | ns | 0.0784 |
| 6:Ldo45$^{GFP}$ vs. 24:Ldo45$^{GFP}$ | -0.1463 | -0.3989 to 0.1063 | ns | 0.5391 |
| 6:Ldo45$^{GFP}$ vs. 24:Ldo16$^{GFP}$ | -0.58 | -0.8326 to -0.3274 | **** | <0.0001 |
| 6:Ldo45$^{GFP}$ vs. 48:Ldo45$^{GFP}$ | 0.1247 | -0.1279 to 0.3772 | ns | 0.7134 |
| 6:Ldo45$^{GFP}$ vs. 48:Ldo16$^{GFP}$ | -0.8335 | -1.086 to -0.5809 | **** | <0.0001 |
| 6:Ldo16$^{GFP}$ vs. 12:Ldo45$^{GFP}$ | -0.5069 | -0.7594 to -0.2543 | **** | <0.0001 |
| 6:Ldo16$^{GFP}$ vs. 12:Ldo16$^{GFP}$ | -0.6229 | -0.8755 to -0.3704 | **** | <0.0001 |
| 6:Ldo16$^{GFP}$ vs. 24:Ldo45$^{GFP}$ | -0.5334 | -0.7860 to -0.2808 | **** | <0.0001 |
| 6:Ldo16$^{GFP}$ vs. 24:Ldo16$^{GFP}$ | -0.9671 | -1.220 to -0.7145 | **** | <0.0001 |
| 6:Ldo16$^{GFP}$ vs. 48:Ldo45$^{GFP}$ | -0.2624 | -0.5150 to -0.009828 | * | 0.0381 |
| 6:Ldo16$^{GFP}$ vs. 48:Ldo16$^{GFP}$ | -1.221 | -1.473 to -0.9680 | **** | <0.0001 |
| 12:Ldo45$^{GFP}$ vs. 12:Ldo16$^{GFP}$ | -0.1161 | -0.3687 to 0.1365 | ns | 0.7772 |
| 12:Ldo45$^{GFP}$ vs. 24:Ldo45$^{GFP}$ | -0.02654 | -0.2791 to 0.2260 | ns | >0.9999 |
| 12:Ldo45$^{GFP}$ vs. 24:Ldo16$^{GFP}$ | -0.4602 | -0.7128 to -0.2077 | *** | 0.0001 |
| 12:Ldo45$^{GFP}$ vs. 48:Ldo45$^{GFP}$ | 0.2444 | -0.008126 to 0.4970 | ns | 0.0624 |
| 12:Ldo45$^{GFP}$ vs. 48:Ldo16$^{GFP}$ | -0.7137 | -0.9663 to -0.4612 | **** | <0.0001 |
| 12:Ldo16$^{GFP}$ vs. 24:Ldo45$^{GFP}$ | 0.08956 | -0.1630 to 0.3421 | ns | 0.9263 |
| 12:Ldo16$^{GFP}$ vs. 24:Ldo16$^{GFP}$ | -0.3441 | -0.5967 to -0.09155 | ** | 0.0034 |
| 12:Ldo16$^{GFP}$ vs. 48:Ldo45$^{GFP}$ | 0.3605 | 0.1080 to 0.6131 | ** | 0.0021 |
| 12:Ldo16$^{GFP}$ vs. 48:Ldo16$^{GFP}$ | -0.5976 | -0.8502 to -0.3451 | **** | <0.0001 |
| 24:Ldo45$^{GFP}$ vs. 24:Ldo16$^{GFP}$ | -0.4337 | -0.6863 to -0.1811 | *** | 0.0002 |
| 24:Ldo45$^{GFP}$ vs. 48:Ldo45$^{GFP}$ | 0.271 | 0.01842 to 0.5236 | * | 0.0299 |
| 24:Ldo45$^{GFP}$ vs. 48:Ldo16$^{GFP}$ | -0.6872 | -0.9398 to -0.4346 | **** | <0.0001 |
| 24:Ldo16$^{GFP}$ vs. 48:Ldo45$^{GFP}$ | 0.7047 | 0.4521 to 0.9572 | **** | <0.0001 |
| 24:Ldo16$^{GFP}$ vs. 48:Ldo16$^{GFP}$ | -0.2535 | -0.5061 to -0.0009308 | * | 0.0487 |
| 48:Ldo45$^{GFP}$ vs. 48:Ldo16$^{GFP}$ | -0.9582 | -1.211 to -0.7056 | **** | <0.0001 |

| **Figure 2G** | |
|---|---|
| **ANOVA summary** | |
| **F** | 8.34 |
| **P value** | 0.0029 |
| **P value summary** | ** |
| **Significant diff. among means (P < 0.05)?** | Yes |
| **R squared** | 0.6759 |
| **F (DFn, DFd)** | 0.2727 (3, 12) |

| **Tukey's multiple comparisons test** | **Mean Diff.** | **95.00% CI of diff.** | **Summary** | **Adjusted P Value** |
|---|---|---|---|---|

| | | | | |
|---|---|---|---|---|
| WT vs. ΔΔ*ldo* | 0.6293 | 0.2370 to 1.022 | ** | 0.0022 |
| WT vs. Δ*ldo16* | 0.425 | 0.03269 to 0.8172 | * | 0.0325 |
| WT vs. Δ*ldo45* | 0.2235 | -0.1688 to 0.6157 | ns | 0.3694 |
| ΔΔ*ldo* vs. Δ*ldo16* | -0.2043 | -0.5966 to 0.1879 | ns | 0.4423 |
| ΔΔ*ldo* vs. Δ*ldo45* | -0.4058 | -0.7981 to -0.01356 | * | 0.0419 |
| Δ*ldo16* vs. Δ*ldo45* | -0.2015 | -0.5938 to 0.1908 | ns | 0.4537 |

**Figure 5E**

| **Two-way ANOVA** | **P value** | **P value summary** | **F (DFn, DFd)** |
|---|---|---|---|
| **Row Factor** | 0.3946 | ns | F (4, 8) = 1.164 |
| **Column Factor** | 0.4276 | ns | F (2, 4) = 1.058 |
| **Row Factor x Column Factor** | <0.0001 | **** | F (8, 16) = 54.85 |

| **Tukey's multiple comparisons test** | **Mean Diff.** | **95.00% CI of diff.** | **Summary** | **Adjusted P Value** |
|---|---|---|---|---|
| **LD surface** | | | | |
| WT vs. Vac8Δ12_α3 | -26.2 | -46.87 to -5.531 | ** | 0.0099 |
| WT vs. Vac8Δ12_loop,α3 | -14.65 | -35.32 to 6.016 | ns | 0.2393 |
| WT vs. Vac8Δ12_α2,α3 | -71.92 | -92.58 to -51.25 | **** | <0.0001 |
| WT vs. Vac8Δ12 | -65.94 | -86.60 to -45.27 | **** | <0.0001 |
| Vac8Δ12_α3 vs. Vac8Δ12_loop,α3 | 11.55 | -9.122 to 32.21 | ns | 0.4547 |
| Vac8Δ12_α3 vs. Vac8Δ12_α2,α3 | -45.72 | -66.39 to -25.05 | **** | <0.0001 |
| Vac8Δ12_α3 vs. Vac8Δ12 | -39.74 | -60.41 to -19.07 | *** | 0.0002 |
| Vac8Δ12_loop,α3 vs. Vac8Δ12_α2,α3 | -57.26 | -77.93 to -36.60 | **** | <0.0001 |
| Vac8Δ12_loop,α3 vs. Vac8Δ12 | -51.28 | -71.95 to -30.62 | **** | <0.0001 |
| Vac8Δ12_α2,α3 vs. Vac8Δ12 | 5.981 | -14.69 to 26.65 | ns | 0.8978 |
| **vCLIP + LD surface** | | | | |
| WT vs. Vac8Δ12_α3 | -39.05 | -59.72 to -18.38 | *** | 0.0002 |
| WT vs. Vac8Δ12_loop,α3 | -37.99 | -58.66 to -17.32 | *** | 0.0003 |
| WT vs. Vac8Δ12_α2,α3 | -10.99 | -31.66 to 9.681 | ns | 0.5015 |
| WT vs. Vac8Δ12 | -19.11 | -39.78 to 1.553 | ns | 0.077 |
| Vac8Δ12_α3 vs. Vac8Δ12_loop,α3 | 1.055 | -19.61 to 21.72 | ns | 0.9998 |
| Vac8Δ12_α3 vs. Vac8Δ12_α2,α3 | 28.06 | 7.393 to 48.73 | ** | 0.0057 |
| Vac8Δ12_α3 vs. Vac8Δ12 | 19.93 | -0.7347 to 40.60 | ns | 0.0614 |
| Vac8Δ12_loop,α3 vs. Vac8Δ12_α2,α3 | 27.01 | 6.338 to 47.67 | ** | 0.0078 |
| Vac8Δ12_loop,α3 vs. Vac8Δ12 | 18.88 | -1.790 to 39.55 | ns | 0.0822 |
| Vac8Δ12_α2,α3 vs. Vac8Δ12 | -8.128 | -28.80 to 12.54 | ns | 0.7488 |
| **vCLIP** | | | | |
| WT vs. Vac8Δ12_α3 | 65.25 | 44.58 to 85.91 | **** | <0.0001 |
| WT vs. Vac8Δ12_loop,α3 | 52.65 | 31.98 to 73.31 | **** | <0.0001 |
| WT vs. Vac8Δ12_α2,α3 | 82.9 | 62.24 to 103.6 | **** | <0.0001 |
| WT vs. Vac8Δ12 | 85.05 | 64.38 to 105.7 | **** | <0.0001 |
| Vac8Δ12_α3 vs. Vac8Δ12_loop,α3 | -12.6 | -33.27 to 8.067 | ns | 0.3721 |
| Vac8Δ12_α3 vs. Vac8Δ12_α2,α3 | 17.66 | -3.010 to 38.33 | ns | 0.1139 |
| Vac8Δ12_α3 vs. Vac8Δ12 | 19.8 | -0.8631 to 40.47 | ns | 0.0637 |
| Vac8Δ12_loop,α3 vs. Vac8Δ12_α2,α3 | 30.26 | 9.591 to 50.93 | ** | 0.003 |
| Vac8Δ12_loop,α3 vs. Vac8Δ12 | 32.41 | 11.74 to 53.07 | ** | 0.0016 |
| Vac8Δ12_α2,α3 vs. Vac8Δ12 | 2.147 | -18.52 to 22.82 | ns | 0.9975 |

| Figure 5F | | | |
|---|---|---|---|
| **Two-way ANOVA** | **P value** | **P value summary** | **F (DFn, DFd)** |
| **Row Factor** | >0.9999 | ns | F (4, 45) = 6.735e-006 |
| **Column Factor** | <0.0001 | **** | F (4, 45) = 101.3 |
| **Row Factor x Column Factor** | <0.0001 | **** | F (16, 45) = 106.3 |

| Tukey's multiple comparisons test | Predicted (LS) mean diff. | 95.00% CI of diff. | Summary | Adjusted P Value |
|---|---|---|---|---|
| **nER** | | | | |
| WT vs. Vac8Δ12_α3 | -8.3 | -18.53 to 1.930 | ns | 0.1621 |
| WT vs. Vac8Δ12_loop, α3 | -15.58 | -27.02 to -4.146 | ** | 0.0031 |
| WT vs. Vac8Δ12_α2,α3 | -57 | -67.23 to -46.77 | **** | <0.0001 |
| WT vs. Vac8Δ12 | -93.03 | -103.3 to -82.80 | **** | <0.0001 |
| Vac8Δ12_α3 vs. Vac8Δ12_loop,α3 | -7.283 | -18.72 to 4.154 | ns | 0.381 |
| Vac8Δ12_α3 vs. Vac8Δ12_α2,α3 | -48.7 | -58.93 to -38.47 | **** | <0.0001 |
| Vac8Δ12_α3 vs. Vac8Δ12 | -84.73 | -94.96 to -74.50 | **** | <0.0001 |
| Vac8Δ12_loop,α3 vs. Vac8Δ12_α2,α3 | -41.42 | -52.85 to -29.98 | **** | <0.0001 |
| Vac8Δ12_loop,α3 vs. Vac8Δ12 | -77.45 | -88.89 to -66.01 | **** | <0.0001 |
| Vac8Δ12_α2,α3 vs. Vac8Δ12 | -36.03 | -46.26 to -25.80 | **** | <0.0001 |
| **nER + NVJ foci** | | | | |
| WT vs. Vac8Δ12_α3 | -22.23 | -32.46 to -12.00 | **** | <0.0001 |
| WT vs. Vac8Δ12_loop,α3 | -32.98 | -44.42 to -21.55 | **** | <0.0001 |
| WT vs. Vac8Δ12_α2,α3 | -38.9 | -49.13 to -28.67 | **** | <0.0001 |
| WT vs. Vac8Δ12 | -3.367 | -13.60 to 6.864 | ns | 0.8817 |
| Vac8Δ12_α3 vs. Vac8Δ12_loop,α3 | -10.75 | -22.19 to 0.6878 | ns | 0.0746 |
| Vac8Δ12_α3 vs. Vac8Δ12_α2,α3 | -16.67 | -26.90 to -6.436 | *** | 0.0003 |
| Vac8Δ12_α3 vs. Vac8Δ12 | 18.87 | 8.636 to 29.10 | **** | <0.0001 |
| Vac8Δ12_loop,α3 vs. Vac8Δ12_α2,α3 | -5.917 | -17.35 to 5.521 | ns | 0.5869 |
| Vac8Δ12_loop,α3 vs. Vac8Δ12 | 29.62 | 18.18 to 41.05 | **** | <0.0001 |
| Vac8Δ12_α2,α3 vs. Vac8Δ12 | 35.53 | 25.30 to 45.76 | **** | <0.0001 |
| **NVJ foci** | | | | |
| WT vs. Vac8Δ12_α3 | -12.1 | -22.33 to -1.870 | * | 0.0131 |
| WT vs. Vac8Δ12_loop,α3 | -7.183 | -18.62 to 4.254 | ns | 0.395 |
| WT vs. Vac8Δ12_α2,α3 | 33.93 | 23.70 to 44.16 | **** | <0.0001 |
| WT vs. Vac8Δ12 | 34.37 | 24.14 to 44.60 | **** | <0.0001 |
| Vac8Δ12_α3 vs. Vac8Δ12_loop,α3 | 4.917 | -6.521 to 16.35 | ns | 0.7391 |
| Vac8Δ12_α3 vs. Vac8Δ12_α2,α3 | 46.03 | 35.80 to 56.26 | **** | <0.0001 |
| Vac8Δ12_α3 vs. Vac8Δ12 | 46.47 | 36.24 to 56.70 | **** | <0.0001 |
| Vac8Δ12_loop,α3 vs. Vac8Δ12_α2,α3 | 41.12 | 29.68 to 52.55 | **** | <0.0001 |
| Vac8Δ12_loop,α3 vs. Vac8Δ12 | 41.55 | 30.11 to 52.99 | **** | <0.0001 |
| Vac8Δ12_α2,α3 vs. Vac8Δ12 | 0.4333 | -9.797 to 10.66 | ns | >0.9999 |
| **NVJ elongated** | | | | |
| WT vs. Vac8Δ12_α3 | 36.8 | 26.57 to 47.03 | **** | <0.0001 |
| WT vs. Vac8Δ12_loop,α3 | 44.95 | 33.51 to 56.39 | **** | <0.0001 |
| WT vs. Vac8Δ12_α2,α3 | 51.17 | 40.94 to 61.40 | **** | <0.0001 |
| WT vs. Vac8Δ12 | 51.23 | 41.00 to 61.46 | **** | <0.0001 |
| Vac8Δ12_α3 vs. Vac8Δ12_loop,α3 | 8.15 | -3.288 to 19.59 | ns | 0.2711 |

| | | | | |
|---|---|---|---|---|
| Vac8Δ12_α3 vs. Vac8Δ12_α2,α3 | 14.37 | 4.136 to 24.60 | ** | 0.0021 |
| Vac8Δ12_α3 vs. Vac8Δ12 | 14.43 | 4.203 to 24.66 | ** | 0.002 |
| Vac8Δ12_loop,α3 vs. Vac8Δ12_α2,α3 | 6.217 | -5.221 to 17.65 | ns | 0.5399 |
| Vac8Δ12_loop,α3 vs. Vac8Δ12 | 6.283 | -5.154 to 17.72 | ns | 0.5295 |
| Vac8Δ12_α2,α3 vs. Vac8Δ12 | 0.06667 | -10.16 to 10.30 | ns | >0.9999 |
| **NVJ with PMN** | | | | |
| WT vs. Vac8Δ12_α3 | 5.867 | -4.364 to 16.10 | ns | 0.487 |
| WT vs. Vac8Δ12_loop,α3 | 10.8 | -0.6378 to 22.24 | ns | 0.0725 |
| WT vs. Vac8Δ12_α2,α3 | 10.8 | 0.5697 to 21.03 | * | 0.0339 |
| WT vs. Vac8Δ12 | 10.8 | 0.5697 to 21.03 | * | 0.0339 |
| Vac8Δ12_α3 vs. Vac8Δ12_loop,α3 | 4.933 | -6.504 to 16.37 | ns | 0.7367 |
| Vac8Δ12_α3 vs. Vac8Δ12_α2,α3 | 4.933 | -5.297 to 15.16 | ns | 0.6494 |
| Vac8Δ12_α3 vs. Vac8Δ12 | 4.933 | -5.297 to 15.16 | ns | 0.6494 |
| Vac8Δ12_loop,α3 vs. Vac8Δ12_α2,α3 | -2.842E-14 | -11.44 to 11.44 | ns | >0.9999 |
| Vac8Δ12_loop,α3 vs. Vac8Δ12 | -2.842E-14 | -11.44 to 11.44 | ns | >0.9999 |
| Vac8Δ12_α2,α3 vs. Vac8Δ12 | 0 | -10.23 to 10.23 | ns | >0.9999 |

| **Figure 5G** | |
|---|---|
| **ANOVA summary** | |
| **F** | 8.34 |
| **P value** | 0.0029 |
| **P value summary** | ** |
| **Significant diff. among means (P < 0.05)?** | Yes |
| **R squared** | 0.6759 |
| **F (DFn, DFd)** | 0.2727 (3, 12) |

| **Tukey's multiple comparisons test** | **Mean Diff.** | **95.00% CI of diff.** | **Summary** | **Adjusted P Value** |
|---|---|---|---|---|
| WT vs. Vac8Δ12 | 0.468 | 0.2514 to 0.6846 | **** | <0.0001 |
| WT vs. Vac8Δ12_α2,α3 | 0.3031 | 0.08652 to 0.5198 | ** | 0.0025 |
| WT vs. Vac8Δ12_α3 | 0.218 | 0.001366 to 0.4346 | * | 0.0479 |
| WT vs. Vac8Δ12_loop,α3 | 0.1899 | -0.02673 to 0.4065 | ns | 0.1117 |
| WT vs. Δ*vac8* | 0.5177 | 0.2755 to 0.7599 | **** | <0.0001 |
| Vac8Δ12 vs. Vac8Δ12_α2,α3 | -0.1649 | -0.3815 to 0.05177 | ns | 0.2176 |
| Vac8Δ12 vs. Vac8Δ12_α3 | -0.25 | -0.4666 to -0.03338 | * | 0.0166 |
| Vac8Δ12 vs. Vac8Δ12_loop,α3 | -0.2781 | -0.4947 to -0.06148 | ** | 0.0062 |
| Vac8Δ12 vs. Δ*vac8* | 0.04973 | -0.1925 to 0.2919 | ns | 0.988 |
| Vac8Δ12_α2,α3 vs. Vac8Δ1_α3 | -0.08515 | -0.3018 to 0.1315 | ns | 0.8325 |
| Vac8Δ12_α2,α3 vs. Vac8Δ12_loop,α3 | -0.1132 | -0.3299 to 0.1034 | ns | 0.6068 |
| Vac8Δ12_α2,α3 vs. Δvac8 | 0.2146 | -0.02761 to 0.4568 | ns | 0.1053 |
| Vac8Δ12_α3 vs. Vac8Δ12_loop,α3 | -0.0281 | -0.2447 to 0.1885 | ns | 0.9986 |
| Vac8Δ12_α3 vs. Δ*vac8* | 0.2997 | 0.05754 to 0.5419 | ** | 0.0088 |
| Vac8Δ12_loop,α3 vs. Δ*vac8* | 0.3278 | 0.08564 to 0.5700 | ** | 0.0036 |

| **Figure 6B** | |
|---|---|
| **Welch´s ANOVA summary** | |
| **P value** | <0.0001 |
| **P value summary** | **** |

| | |
|---|---|
| **Significant diff. among means (P < 0.05)?** | Yes |
| **W (DFn, DFd)** | 84.13 (3.000. 69.33) |

| **Dunnett's T3 multiple comparisons test** | **Mean Diff.** | **95.00% CI of diff.** | **Summary** | **Adjusted P Value** |
|---|---|---|---|---|
| vCLIP vs. NVJ | -12.94 | -48.01 to 22.13 | ns | 0.8481 |
| vCLIP vs. Vacuolar membrane | 23.37 | -0.9541 to 47.70 | * | 0.0345 |
| vCLIP vs. nER | 69.54 | 47.30 to 91.79 | **** | <0.0001 |
| NVJ vs. Vacuolar membrane | 36.32 | 5.665 to 66.97 | ** | 0.0054 |
| NVJ vs. nER | 82.49 | 53.35 to 111.6 | **** | <0.0001 |
| Vacuolar membrane vs. nER | 46.17 | 33.81 to 58.53 | **** | <0.0001 |

**Figure 7B**

| **Two-way ANOVA** | **P value** | **P value summary** | **F (DFn, DFd)** |
|---|---|---|---|
| **Row Factor** | 0.1077 | ns | F (2, 8) = 2.983 |
| **Column Factor** | 0.2551 | ns | F (1, 4) = 1.762 |
| **Row Factor x Column Factor** | 0.2627 | ns | F (2, 8) = 1.587 |

| **Tukey's multiple comparisons test** | **Mean Diff.** | **95.00% CI of diff.** | **Summary** | **Adjusted P Value** |
|---|---|---|---|---|
| 24 h:WT vs. 24 h:ΔΔ*ldo* | 1.227 | -0.8252 to 3.279 | ns | 0.3371 |
| 24 h:WT vs. 48 h:WT | 1.347 | -0.7052 to 3.399 | ns | 0.2602 |
| 24 h:WT vs. 48 h:ΔΔ*ldo* | 1.492 | -0.5606 to 3.544 | ns | 0.1876 |
| 24 h:WT vs. 72 h:WT | 1.676 | -0.3760 to 3.728 | ns | 0.1219 |
| 24 h:WT vs. 72 h:ΔΔ*ldo* | 1.572 | -0.4798 to 3.625 | ns | 0.1556 |
| 24 h:ΔΔ*ldo* vs. 48 h:WT | 0.12 | -1.932 to 2.172 | ns | >0.9999 |
| 24 h:ΔΔ*ldo* vs. 48 h:ΔΔ*ldo* | 0.2646 | -1.788 to 2.317 | ns | 0.996 |
| 24 h:ΔΔ*ldo* vs. 72 h:WT | 0.4491 | -1.603 to 2.501 | ns | 0.9597 |
| 24 h:ΔΔ*ldo* vs. 72 h:ΔΔ*ldo* | 0.3453 | -1.707 to 2.398 | ns | 0.9867 |
| 48 h:WT vs. 48 h:ΔΔ*ldo* | 0.1446 | -1.908 to 2.197 | ns | 0.9998 |
| 48 h:WT vs. 72 h:WT | 0.3292 | -1.723 to 2.381 | ns | 0.9892 |
| 48 h:WT vs. 72 h:ΔΔ*ldo* | 0.2254 | -1.827 to 2.278 | ns | 0.9981 |
| 48 h:ΔΔ*ldo* vs. 72 h:WT | 0.1846 | -1.868 to 2.237 | ns | 0.9993 |
| 48 h:ΔΔ*ldo* vs. 72 h:ΔΔ*ldo* | 0.08076 | -1.971 to 2.133 | ns | >0.9999 |
| 72 h:WT vs. 72 h:ΔΔ*ldo* | -0.1038 | -2.156 to 1.948 | ns | >0.9999 |

**Figure 7C**

| **Two-way ANOVA** | **P value** | **P value summary** | **F (DFn, DFd)** |
|---|---|---|---|
| **Row Factor** | 0.0001 | *** | F (2, 8) = 34.45 |
| **Column Factor** | 0.0136 | * | F (1, 4) = 17.70 |
| **Row Factor x Column Factor** | 0.003 | ** | F (2, 8) = 13.09 |

| **Tukey's multiple comparisons test** | **Mean Diff.** | **95.00% CI of diff.** | **Summary** | **Adjusted P Value** |
|---|---|---|---|---|
| 24 h:WT vs. 24 h:ΔΔ*ldo* | 0.1479 | -0.3528 to 0.6487 | ns | 0.8769 |
| 24 h:WT vs. 48 h:WT | -0.7179 | -1.219 to -0.2172 | ** | 0.0069 |
| 24 h:WT vs. 48 h:ΔΔ*ldo* | -0.2183 | -0.7191 to 0.2824 | ns | 0.6237 |
| 24 h:WT vs. 72 h:WT | -1.529 | -2.029 to -1.028 | **** | <0.0001 |
| 24 h:WT vs. 72 h:ΔΔ*ldo* | -0.4018 | -0.9025 to 0.09900 | ns | 0.1307 |
| 24 h:ΔΔ*ldo* vs. 48 h:WT | -0.8658 | -1.367 to -0.3651 | ** | 0.0021 |

| | | | | |
|---|---|---|---|---|
| 24 h:ΔΔ*ldo* vs. 48 h:ΔΔ*ldo* | -0.3662 | -0.8670 to 0.1345 | ns | 0.1836 |
| 24 h:ΔΔ*ldo* vs. 72 h:WT | -1.677 | -2.177 to -1.176 | **** | <0.0001 |
| 24 h:ΔΔ*ldo* vs. 72 h:ΔΔ*ldo* | -0.5497 | -1.050 to -0.04891 | * | 0.0313 |
| 48 h:WT vs. 48 h:ΔΔ*ldo* | 0.4996 | -0.001178 to 1.000 | ns | 0.0506 |
| 48 h:WT vs. 72 h:WT | -0.8108 | -1.312 to -0.3100 | ** | 0.0032 |
| 48 h:WT vs. 72 h:ΔΔ*ldo* | 0.3162 | -0.1846 to 0.8169 | ns | 0.2912 |
| 48 h:ΔΔ*ldo* vs. 72 h:WT | -1.31 | -1.811 to -0.8096 | *** | 0.0001 |
| 48 h:ΔΔ*ldo* vs. 72 h:ΔΔ*ldo* | -0.1834 | -0.6842 to 0.3173 | ns | 0.7591 |
| 72 h:WT vs. 72 h:ΔΔ*ldo* | 1.127 | 0.6262 to 1.628 | *** | 0.0003 |

| **Figure 7F** | |
|---|---|
| **ANOVA summary** | |
| **F** | 42.97 |
| **P value** | <0.0001 |
| **P value summary** | **** |
| **Significant diff. among means (P < 0.05)?** | Yes |
| **R squared** | 0.9513 |
| **F (DFn, DFd)** | 1.298 (5, 11) |

| **Tukey's multiple comparisons test** | **Mean Diff.** | **95.00% CI of diff.** | **Summary** | **Adjusted P Value** |
|---|---|---|---|---|
| WT vs. Δ*nvj1* | -0.3361 | -0.7136 to 0.04150 | ns | 0.0907 |
| WT vs. Δ*nvj1*Δ*mdm1* | -0.4744 | -0.8519 to -0.09684 | * | 0.0123 |
| WT vs. Δ*vac8* | 0.8419 | 0.4644 to 1.219 | *** | 0.0001 |
| WT vs. ΔΔ*ldo* | 0.5011 | 0.1235 to 0.8786 | ** | 0.0084 |
| WT vs. ΔΔ*ldo*Δ*nvj1*Δ*mdm1* | 0.4645 | 0.04242 to 0.8867 | * | 0.0288 |
| Δ*nvj1* vs. Δ*nvj1*Δ*mdm1* | -0.1383 | -0.5159 to 0.2392 | ns | 0.8047 |
| Δ*nvj1* vs. Δ*vac8* | 1.178 | 0.8004 to 1.556 | **** | <0.0001 |
| Δ*nvj1* vs. ΔΔ*ldo* | 0.8371 | 0.4596 to 1.215 | *** | 0.0001 |
| Δ*nvj1* vs. ΔΔ*ldo*Δ*nvj1*Δ*mdm1* | 0.8006 | 0.3785 to 1.223 | *** | 0.0005 |
| Δ*nvj1*Δ*mdm1* vs. Δ*vac8* | 1.316 | 0.9387 to 1.694 | **** | <0.0001 |
| Δ*nvj1*Δ*mdm1* vs. ΔΔ*ldo* | 0.9754 | 0.5979 to 1.353 | **** | <0.0001 |
| Δ*nvj1*Δ*mdm1* vs. ΔΔ*ldo*Δ*nvj1*Δ*mdm1* | 0.9389 | 0.5168 to 1.361 | *** | 0.0001 |
| Δ*vac8* vs. ΔΔ*ldo* | -0.3409 | -0.7184 to 0.03670 | ns | 0.0847 |
| Δ*vac8* vs. ΔΔ*ldo*Δ*nvj1*Δ*mdm1* | -0.3774 | -0.7995 to 0.04475 | ns | 0.0888 |
| ΔΔ*ldo* vs. ΔΔ*ldo*Δ*nvj1*Δ*mdm1* | -0.03651 | -0.4586 to 0.3856 | ns | 0.9996 |

| **Figure 7B** | | | |
|---|---|---|---|
| **Two-way ANOVA** | P value | P value summary | F (DFn, DFd) |
| **Time x Column Factor** | 0.2595 | ns | F (1, 4) = 1.724 |
| **Time** | 0.5889 | ns | F (1, 4) = 0.3443 |
| **Column Factor** | 0.0035 | ** | F (1, 4) = 38.04 |
| **Subject** | 0.4551 | ns | F (4, 4) = 1.127 |

| **Bonferroni's multiple comparisons test** | **Mean Diff.** | **95.00% CI of diff.** | **Summary** | **Adjusted P Value** |
|---|---|---|---|---|
| WT - *ΔΔldo* | | | | |
| 24 h | -35.37 | -53.42 to -17.31 | ** | 0.0013 |
| 72 h | -23.55 | -41.61 to -5.500 | * | 0.0142 |

| Figure S1A | | | |
|---|---|---|---|
| **Two-way ANOVA** | P value | P value summary | F (DFn, DFd) |
| **Row Factor** | 0.0042 | ** | F (2, 6) = 15.62 |
| **Column Factor** | 0.0478 | * | F (1, 3) = 10.50 |
| **Row Factor x Column Factor** | 0.0289 | * | F (2, 6) = 6.773 |

| Tukey's multiple comparisons test | Mean Diff. | 95.00% CI of diff. | Summary | Adjusted P Value |
|---|---|---|---|---|
| 8 h:*LDO16* vs. 8 h:*LDO45* | 0 | -1.908 to 1.908 | ns | >0.9999 |
| 8 h:*LDO16* vs. 48 h:*LDO16* | -2.795 | -4.703 to -0.8873 | ** | 0.0084 |
| 8 h:*LDO16* vs. 48 h:*LDO45* | -0.453 | -2.361 to 1.455 | ns | 0.92 |
| 8 h:*LDO16* vs. 72 h:*LDO16* | -2.166 | -4.074 to -0.2583 | * | 0.0287 |
| 8 h:*LDO16* vs. 72 h:*LDO45* | -0.2498 | -2.158 to 1.658 | ns | 0.9931 |
| 8 h:*LDO45* vs. 48 h:*LDO16* | -2.795 | -4.703 to -0.8873 | ** | 0.0084 |
| 8 h:*LDO45* vs. 48 h:*LDO45* | -0.453 | -2.361 to 1.455 | ns | 0.92 |
| 8 h:*LDO45* vs. 72 h:*LDO16* | -2.166 | -4.074 to -0.2583 | * | 0.0287 |
| 8 h:*LDO45* vs. 72 h:*LDO45* | -0.2498 | -2.158 to 1.658 | ns | 0.9931 |
| 48 h:*LDO16* vs. 48 h:*LDO45* | 2.342 | 0.4342 to 4.250 | * | 0.02 |
| 48 h:*LDO16* vs. 72 h:*LDO16* | 0.6289 | -1.279 to 2.537 | ns | 0.7714 |
| 48 h:*LDO16* vs. 72 h:*LDO45* | 2.545 | 0.6375 to 4.453 | * | 0.0134 |
| 48 h:*LDO45* vs. 72 h:*LDO16* | -1.713 | -3.621 to 0.1947 | ns | 0.0774 |
| 48 h:*LDO45* vs. 72 h:*LDO45* | 0.2032 | -1.704 to 2.111 | ns | 0.9973 |
| 72 h:*LDO16* vs. 72 h:*LDO45* | 1.916 | 0.008538 to 3.824 | * | 0.0491 |

| Figure S1C | | | |
|---|---|---|---|
| **Two-way ANOVA** | **P value** | **P value summary** | **F (DFn, DFd)** |
| **Row Factor** | 0.0011 | ** | F (3, 9) = 13.56 |
| **Column Factor** | 0.0051 | ** | F (1, 3) = 54.47 |
| **Row Factor x Column Factor** | 0.0023 | ** | F (3, 9) = 11.06 |

| Tukey's multiple comparisons test | Mean Diff. | 95.00% CI of diff. | Summary | Adjusted P Value |
|---|---|---|---|---|
| 6:Ldo45$^{GFP}$ vs. 6:Ldo16$^{GFP}$ | -1.027 | -3.789 to 1.735 | ns | 0.825 |
| 6:Ldo45$^{GFP}$ vs. 12:Ldo45$^{GFP}$ | -0.3489 | -3.111 to 2.413 | ns | 0.9995 |
| 6:Ldo45$^{GFP}$ vs. 12:Ldo16$^{GFP}$ | -3.077 | -5.839 to -0.3149 | * | 0.0275 |
| 6:Ldo45$^{GFP}$ vs. 24:Ldo45$^{GFP}$ | -0.5394 | -3.301 to 2.223 | ns | 0.9922 |
| 6:Ldo45$^{GFP}$ vs. 24:Ldo16$^{GFP}$ | -6.291 | -9.053 to -3.529 | *** | 0.0002 |
| 6:Ldo45$^{GFP}$ vs. 48:Ldo45$^{GFP}$ | -0.7799 | -3.542 to 1.982 | ns | 0.9451 |
| 6:Ldo45$^{GFP}$ vs. 48:Ldo16$^{GFP}$ | -6.706 | -9.468 to -3.944 | *** | 0.0001 |
| 6:Ldo16$^{GFP}$ vs. 12:Ldo45$^{GFP}$ | 0.6779 | -2.084 to 3.440 | ns | 0.9726 |
| 6:Ldo16$^{GFP}$ vs. 12:Ldo16$^{GFP}$ | -2.05 | -4.812 to 0.7119 | ns | 0.193 |
| 6:Ldo16$^{GFP}$ vs. 24:Ldo45$^{GFP}$ | 0.4874 | -2.275 to 3.249 | ns | 0.9957 |
| 6:Ldo16$^{GFP}$ vs. 24:Ldo16$^{GFP}$ | -5.264 | -8.026 to -2.502 | *** | 0.0007 |
| 6:Ldo16$^{GFP}$ vs. 48:Ldo45$^{GFP}$ | 0.2469 | -2.515 to 3.009 | ns | >0.9999 |
| 6:Ldo16$^{GFP}$ vs. 48:Ldo16$^{GFP}$ | -5.679 | -8.442 to -2.917 | *** | 0.0004 |
| 12:Ldo45$^{GFP}$ vs. 12:Ldo16$^{GFP}$ | -2.728 | -5.490 to 0.03402 | ns | 0.0534 |
| 12:Ldo45$^{GFP}$ vs. 24:Ldo45$^{GFP}$ | -0.1905 | -2.953 to 2.572 | ns | >0.9999 |
| 12:Ldo45$^{GFP}$ vs. 24:Ldo16$^{GFP}$ | -5.942 | -8.704 to -3.180 | *** | 0.0003 |
| 12:Ldo45$^{GFP}$ vs. 48:Ldo45$^{GFP}$ | -0.431 | -3.193 to 2.331 | ns | 0.9979 |

| | | | | |
|---|---|---|---|---|
| 12:Ldo45$^{GFP}$ vs. 48:Ldo16$^{GFP}$ | -6.357 | -9.119 to -3.595 | *** | 0.0002 |
| 12:Ldo16$^{GFP}$ vs. 24:Ldo45$^{GFP}$ | 2.538 | -0.2245 to 5.300 | ns | 0.0769 |
| 12:Ldo16$^{GFP}$ vs. 24:Ldo16$^{GFP}$ | -3.214 | -5.976 to -0.4521 | * | 0.0213 |
| 12:Ldo16$^{GFP}$ vs. 48:Ldo45$^{GFP}$ | 2.297 | -0.4650 to 5.059 | ns | 0.1217 |
| 12:Ldo16$^{GFP}$ vs. 48:Ldo16$^{GFP}$ | -3.629 | -6.391 to -0.8672 | ** | 0.01 |
| 24:Ldo45$^{GFP}$ vs. 24:Ldo16$^{GFP}$ | -5.752 | -8.514 to -2.990 | *** | 0.0004 |
| 24:Ldo45$^{GFP}$ vs. 48:Ldo45$^{GFP}$ | -0.2405 | -3.003 to 2.522 | ns | >0.9999 |
| 24:Ldo45$^{GFP}$ vs. 48:Ldo16$^{GFP}$ | -6.167 | -8.929 to -3.405 | *** | 0.0002 |
| 24:Ldo16$^{GFP}$ vs. 48:Ldo45$^{GFP}$ | 5.511 | 2.749 to 8.273 | *** | 0.0005 |
| 24:Ldo16$^{GFP}$ vs. 48:Ldo16$^{GFP}$ | -0.4151 | -3.177 to 2.347 | ns | 0.9984 |
| 48:Ldo45$^{GFP}$ vs. 48:Ldo16$^{GFP}$ | -5.926 | -8.688 to -3.164 | *** | 0.0003 |

**Figure S1D**

| Two-way ANOVA | P value | P value summary | F (DFn, DFd) |
|---|---|---|---|
| **Row Factor** | 0.0013 | ** | F (3, 12) = 10.19 |
| **Column Factor** | 0.9727 | ns | F (1, 4) = 0.001330 |
| **Row Factor x Column Factor** | <0.0001 | **** | F (3, 12) = 19.28 |

| Tukey's multiple comparisons test | Mean Diff. | 95.00% CI of diff. | Summary | Adjusted P Value |
|---|---|---|---|---|
| 6 h:Ldo16$^{GFP}$ vs. 6 h:Ldo45$^{GFP}$ | -0.827 | -1.356 to -0.2979 | ** | 0.0019 |
| 6 h:Ldo16$^{GFP}$ vs. 12 h:Ldo16$^{GFP}$ | -0.4663 | -0.9954 to 0.06286 | ns | 0.1001 |
| 6 h:Ldo16$^{GFP}$ vs. 12 h:Ldo45$^{GFP}$ | -0.6526 | -1.182 to -0.1235 | * | 0.0125 |
| 6 h:Ldo16$^{GFP}$ vs. 24 h:Ldo16$^{GFP}$ | -1.358 | -1.888 to -0.8293 | **** | <0.0001 |
| 6 h:Ldo16$^{GFP}$ vs. 24 h:Ldo45$^{GFP}$ | -0.9681 | -1.497 to -0.4390 | *** | 0.0005 |
| 6 h:Ldo16$^{GFP}$ vs. 48 h:Ldo16$^{GFP}$ | -1.663 | -2.192 to -1.134 | **** | <0.0001 |
| 6 h:Ldo16$^{GFP}$ vs. 48 h:Ldo45$^{GFP}$ | -1.059 | -1.588 to -0.5295 | *** | 0.0002 |
| 6 h:Ldo45$^{GFP}$ vs. 12 h:Ldo16$^{GFP}$ | 0.3607 | -0.1684 to 0.8899 | ns | 0.2931 |
| 6 h:Ldo45$^{GFP}$ vs. 12 h:Ldo45$^{GFP}$ | 0.1744 | -0.3548 to 0.7035 | ns | 0.9195 |
| 6 h:Ldo45$^{GFP}$ vs. 24 h:Ldo16$^{GFP}$ | -0.5314 | -1.061 to -0.002261 | * | 0.0488 |
| 6 h:Ldo45$^{GFP}$ vs. 24 h:Ldo45$^{GFP}$ | -0.1411 | -0.6703 to 0.3880 | ns | 0.9714 |
| 6 h:Ldo45$^{GFP}$ vs. 48 h:Ldo16$^{GFP}$ | -0.8363 | -1.365 to -0.3071 | ** | 0.0017 |
| 6 h:Ldo45$^{GFP}$ vs. 48 h:Ldo45$^{GFP}$ | -0.2316 | -0.7608 to 0.2975 | ns | 0.751 |
| 12 h:Ldo16$^{GFP}$ vs. 12 h:Ldo45$^{GFP}$ | -0.1864 | -0.7155 to 0.3428 | ns | 0.8919 |
| 12 h:Ldo16$^{GFP}$ vs. 24 h:Ldo16$^{GFP}$ | -0.8921 | -1.421 to -0.3630 | *** | 0.001 |
| 12 h:Ldo16$^{GFP}$ vs. 24 h:Ldo45$^{GFP}$ | -0.5019 | -1.031 to 0.02729 | ns | 0.0678 |
| 12 h:Ldo16$^{GFP}$ vs. 48 h:Ldo16$^{GFP}$ | -1.197 | -1.726 to -0.6679 | **** | <0.0001 |
| 12 h:Ldo16$^{GFP}$ vs. 48 h:Ldo45$^{GFP}$ | -0.5924 | -1.122 to -0.06321 | * | 0.0246 |
| 12 h:Ldo45$^{GFP}$ vs. 24 h:Ldo16$^{GFP}$ | -0.7058 | -1.235 to -0.1766 | ** | 0.0069 |
| 12 h:Ldo45$^{GFP}$ vs. 24 h:Ldo45$^{GFP}$ | -0.3155 | -0.8446 to 0.2137 | ns | 0.4342 |
| 12 h:Ldo45$^{GFP}$ vs. 48 h:Ldo16$^{GFP}$ | -1.011 | -1.540 to -0.4815 | *** | 0.0003 |
| 12 h:Ldo45$^{GFP}$ vs. 48 h:Ldo45$^{GFP}$ | -0.406 | -0.9351 to 0.1232 | ns | 0.1887 |
| 24 h:Ldo16$^{GFP}$ vs. 24 h:Ldo45$^{GFP}$ | 0.3903 | -0.1389 to 0.9194 | ns | 0.2208 |
| 24 h:Ldo16$^{GFP}$ vs. 48 h:Ldo16$^{GFP}$ | -0.3049 | -0.8340 to 0.2243 | ns | 0.4721 |
| 24 h:Ldo16$^{GFP}$ vs. 48 h:Ldo45$^{GFP}$ | 0.2998 | -0.2294 to 0.8289 | ns | 0.4909 |
| 24 h:Ldo45$^{GFP}$ vs. 48 h:Ldo16$^{GFP}$ | -0.6952 | -1.224 to -0.1660 | ** | 0.0078 |
| 24 h:Ldo45$^{GFP}$ vs. 48 h:Ldo45$^{GFP}$ | -0.0905 | -0.6197 to 0.4386 | ns | 0.9978 |
| 48 h:Ldo16$^{GFP}$ vs. 48 h:Ldo45$^{GFP}$ | 0.6047 | 0.07551 to 1.134 | * | 0.0214 |

| Figure S4C | |
|---|---|
| **ANOVA summary** | |
| **F** | 33.01 |
| **P value** | <0.0001 |
| **P value summary** | **** |
| **Significant diff. among means (P < 0.05)?** | Yes |
| **R squared** | 0.855 |
| **F (DFn, DFd)** | 0.4838 (5, 28) |

| Tukey's multiple comparisons test | Mean Diff. | 95.00% CI of diff. | Summary | Adjusted P Value |
|---|---|---|---|---|
| WT vs. $Vac8^{mScarlet}$ | 0.08191 | -0.09053 to 0.2544 | ns | 0.6963 |
| WT vs. $Vac8^{3HA}$ | 0.3152 | 0.1428 to 0.4877 | **** | <0.0001 |
| WT vs. $^{mScarlet}Vac8$ | 0.522 | 0.3495 to 0.6944 | **** | <0.0001 |
| WT vs. $^{3HA}Vac8$ | 0.5126 | 0.3402 to 0.6850 | **** | <0.0001 |
| WT vs. *Δvac8* | 0.5177 | 0.3249 to 0.7105 | **** | <0.0001 |
| $Vac8^{mScarlet}$ vs. $Vac8^{3HA}$ | 0.2333 | 0.06087 to 0.4058 | ** | 0.0036 |
| $Vac8^{mScarlet}$ vs. $^{mScarlet}Vac8$ | 0.4401 | 0.2676 to 0.6125 | **** | <0.0001 |
| $Vac8^{mScarlet}$ vs. $^{3HA}Vac8$ | 0.4307 | 0.2582 to 0.6031 | **** | <0.0001 |
| $Vac8^{mScarlet}$ vs. *Δvac8* | 0.4358 | 0.2430 to 0.6286 | **** | <0.0001 |
| $Vac8^{3HA}$ vs. $^{mScarlet}Vac8$ | 0.2067 | 0.03430 to 0.3792 | * | 0.0119 |
| $Vac8^{3HA}$ vs. $^{3HA}Vac8$ | 0.1974 | 0.02494 to 0.3698 | * | 0.0178 |
| $Vac8^{3HA}$ vs. *Δvac8* | 0.2025 | 0.009691 to 0.3953 | * | 0.0353 |
| $^{mScarlet}Vac8$ vs. $^{3HA}Vac8$ | -0.009366 | -0.1818 to 0.1631 | ns | >0.9999 |
| $^{mScarlet}Vac8$ vs. *Δvac8* | -0.004257 | -0.1971 to 0.1885 | ns | >0.9999 |
| $^{3HA}Vac8$ vs. *Δvac8* | 0.005109 | -0.1877 to 0.1979 | ns | >0.9999 |
